# Structural Basis for Intermodular Communication in Assembly-Line Polyketide Biosynthesis

**DOI:** 10.1101/2024.05.05.592269

**Authors:** Dillon P. Cogan, Alexander M. Soohoo, Muyuan Chen, Yan Liu, Krystal L. Brodsky, Chaitan Khosla

## Abstract

Assembly-line polyketide synthases are large multienzyme systems with considerable potential for genetic reprogramming. To investigate the mechanisms by which reactive biosynthetic intermediates are directionally channeled across a defined sequence of active sites in a naturally occurring assembly line, we employed a bifunctional reagent to crosslink transient domain-domain interfaces of the 6-deoxyerythronolide B synthase. Structural resolution of these crosslinked states by single-particle cryogenic electron microscopy (cryo-EM) together with statistical per-particle image analysis of cryo-EM data revealed ketosynthase – acyl carrier protein (KS-ACP) interactions that discriminate between intra- and inter-modular communication, while reinforcing the relevance of asymmetric conformations during the catalytic cycle. Our findings provide a new foundation for the structure-based design of hybrid polyketide synthases comprised of biosynthetic modules from different assembly lines.

## INTRODUCTION

Assembly-line polyketide synthases (PKSs) are homodimeric multienzyme systems that synthesize many structurally complex and medicinally important natural products^1–4^. Their modular architecture implies the existence of an inbuilt mechanism for ensuring that a growing polyketide chain is channeled along a uniquely defined sequence of enzymatic active sites, each of which is used only once in the overall catalytic cycle of the PKS^5^. Engineering these systems holds enormous potential for the biosynthesis of user-defined chemical structures^6^. To this end, we have studied 6-deoxyerythronolide B synthase (DEBS; Fig. S1) as a paradigm for decoding the principles of assembly-line polyketide biosynthesis^7^. For a representative catalytic cycle of the first module in this assembly line, see Supplementary Figure S2.

Recently, the application of single-particle cryo-EM techniques has not only opened the door to structural analysis of intact PKS modules at near-atomic resolution, but has also highlighted the conformationally dynamic nature of these modules as they proceed through their catalytic cycles^8–13^ (see Ref. 14 for a recent review on this topic). In this study, we have combined cryo-EM with a largely overlooked covalent crosslinking method to visualize functionally critical domain-domain interfaces in key transient states of the catalytic cycles of representative DEBS modules^15,16^. In turn, these structural insights explain how reactive intermediates are efficiently channeled between successive modules while reaffirming the functional relevance of asymmetry during the PKS’s catalytic cycle.

## RESULTS

### Rapid and site-selective crosslinking of KS and ACP thiols of PKS modules

Previously, we observed an asymmetric conformation (*State 1*) of homodimeric Module 1 (M1) of DEBS by single-particle cryo-EM^12^. (We specifically employed a version of M1, referred to as M1TE, that was N-terminally fused with a docking domain for antibody complexation and C-terminally fused with a thioesterase (TE) domain for improved expression of the recombinant protein.) In this structure, the 4′-phosphopantetheine (Ppant) arm of an acyl carrier protein (ACP) domain from one subunit had been inserted into the active site of the ketosynthase (KS) domain from the other subunit. This conformation corroborated prior biochemical evidence supporting the involvement of inter-subunit KS-ACP interactions during the elongation of a growing polyketide chain by an assembly-line PKS^17^. The cryo-EM structure of *State 1* also led us to propose that the elongation reaction is catalyzed asynchronously by the two equivalent KS-ACP pairs of a homodimeric PKS module. A related study of the lasalocid A PKS independently arrived at the same conclusion^13^.

Based on the observation that the catalytic Cys of the KS domain and the Ppant thiol of the ACP were only ∼4 Å apart in *State 1*^12^, we attempted to covalently crosslink the proximal KS and ACP thiols using 1,3-dibromoacetone (DBA) as a thiol-reactive bis-electrophile. Given the rapid reversibility of Ppant arm docking into the KS active site, we anticipated that a subset of KS and ACP partners in a population of homodimeric modules would each become pre-modified with one equivalent of DBA and therefore unable to crosslink upon KS-ACP docking (Fig. 1A). We therefore added DBA in slight excess (1.25 equivalents) to *holo*-form DEBS M1TE while also providing citrate (≥50 mM), a known promoter of polyketide chain elongation, to the reaction mixture^12,18^. A large excess of β-mercaptoethanol (BME) was added following DBA addition to quench any unreacted electrophiles.

**Fig. 1.**
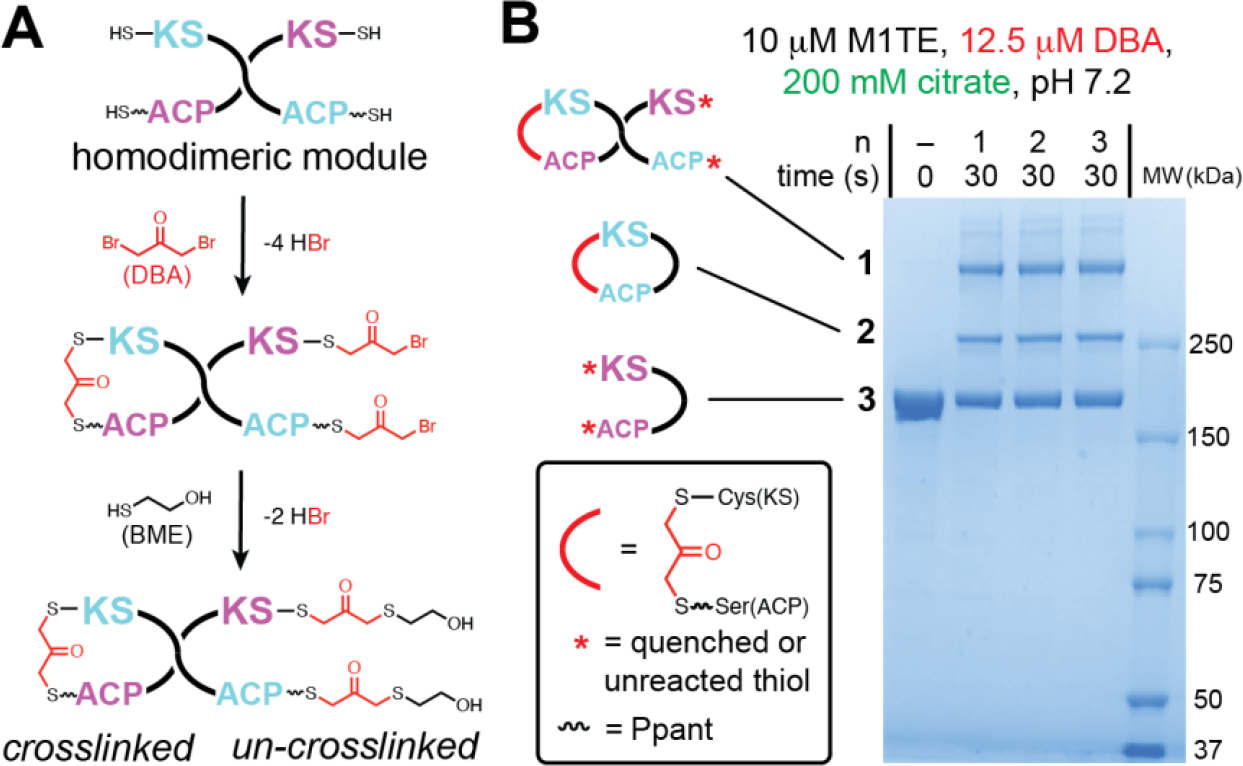
(**A**) Crosslinking a PKS module with 1,3-dibromoacetone (DBA). Only one crosslinking outcome corresponding to the asymmetrically crosslinked *State 1* is shown. (See Fig. S5A for other possible products.) (**B**) Addition of DBA to DEBS M1TE resulted in the formation of two major products as judged by SDS-PAGE (bands **1** and **2**). The un-crosslinked monomer is denoted as band **3**. Three replicate 30 s reactions (n) were analyzed alongside the unreacted M1TE (–). Ppant = 4′-phosphopantetheine. For gel source data, see Supplementary Raw Data.

Product analysis of these crosslinking reactions by SDS-PAGE revealed two putatively crosslinked species (bands **1** and **2**) with decreased electrophoretic mobilities relative to monomeric M1TE (band **3**, Fig. 1B). Similar products were also observed when DBA was added to the DEBS Module 3 analog of M1TE (i.e., M3TE; Fig. S3A). The formation of species **1** and **2** was rapid (reaching ∼50% completion within 5 s) and citrate dependent, suggesting that DBA crosslinking captured conformations related to polyketide chain elongation (Fig. S3B). Absence of the Ppant arm from the ACP domain or mutagenesis of the catalytic Cys residue of the KS greatly abolished crosslinking, highlighting DBA selectivity for KS and ACP thiols (Fig. S4). The decreased electrophoretic mobilities of bands **1** and **2** implied that one of them may correspond to a crosslinked *State 1*, although the origin of the other band was unclear.

### Fluorescence spectroscopic analysis of crosslinked PKS modules

To gain further insight into the identities of bands **1** and **2**, we considered four possible KS-ACP crosslinked species (Fig. S5) based on a previous DBA crosslinking analysis of the mammalian fatty acid synthase (mFAS) which shares the homodimeric architecture and domain organization of assembly-line PKS modules^15^. The putative species contain either one or two KS-ACP crosslinks that occur either within or between PKS subunits, and each was expected to migrate with reduced electrophoretic mobility relative to the un-crosslinked band **3** monomer. Given that one of the observed species probably mimicked the elongation state of a module, we hypothesized that it may also be possible for DBA to capture a state mimicking intermodular chain translocation. If so, then a stand-alone form of the cognate upstream ACP might serve as a probe to crosslink with unoccupied KS domains in bands **1, 2**, or **3**. Based on structural data, it has been proposed that a translocation event might be permitted to occur concomitantly with elongation during polyketide biosynthesis^13^.

To test this hypothesis, ACP2(2) was expressed as a stand-alone protein, purified, and tagged with a fluorophore. ACP2(2) is the cognate substrate donor of DEBS M3; it harbors the C-terminal docking domain of DEBS M2 (denoted by the parenthetical “2”) which is required for native intermolecular interactions between DEBS M2 and M3^19–21^. Recombinant ACP2(2) was tagged with fluorescein isothiocyanate to prepare ACP2(2)-**FL**. To confirm that the *N*-linked fluorescein did not impair ACP binding to its KS partner, ACP2(2)-**FL** was shown to efficiently crosslink to the homodimeric KS-AT didomain fragment of M3 (KS3AT3) in the presence of DBA (Fig. S6).

Upon addition of the ACP2(2)-**FL** probe or a buffer control to purified M3TE prior to DBA crosslinking, two major fluorescent adducts were observed: one that migrated between bands **2** and **3**, consistent with ACP2(2)-**FL** labeling of un-crosslinked M3TE (i.e., band **3** + ACP2(2)-**FL**), and another with a slightly decreased mobility relative to band **1**, supporting the assignment of band **1** as the crosslinked *State 1* (Figs. 1B and S7). Notably, in contrast to extensive conversion of band **1** to its labeled counterpart, no fluorescent band indicative of labeling of band **2** was observed. Not only do these findings attest to the assignment of band **2** as a self-crosslinked monomer (Figs. 1B, S5, and S7), but they also suggest that the unoccupied KS of *State 1* is capable of crosslinking with its translocation ACP partner. Our proposed assignment of bands **1** and **2** is also consistent with previous crosslinking of the mFAS^15^. In that study, two crosslinked species were observed, with the intra-polypeptide crosslinked species also migrating faster than the inter-polypeptide crosslinked species. Further verification of our assignments of bands **1** and **2** was obtained by limited trypsinolysis (Figs. S8–S9), although given that in-gel proteolysis was conducted on denatured DBA-treated M3TE, it was not possible to establish whether band **2** existed as a monomer or dimer prior to denaturation.

### Structure of a prototypical module-module interface during polyketide translocation

Motivated by our observation that KS3AT3 can be efficiently crosslinked to its upstream ACP2(2) partner (Fig. S6), we sought to optimize these crosslinking conditions to facilitate structural analysis of this prototypical module-module interface. Crosslinking of these two proteins was not as strongly dependent on citrate as that between KS3AT3 and its elongation partner ACP3, nor could it be driven to completion even in the presence of a large excess of ACP (Figs. 2 and S10–S11). To compare the crosslinking efficiency of the two ACP partners of KS3AT3, both ACP3 and ACP2(2) were co-incubated with homodimeric KS3AT3 under varying conditions prior to DBA addition (Fig. 2). Conditions were established under which both the elongation and translocation crosslinking events were generally competitive (see Supplementary Methods). A preparative-scale crosslinking reaction was performed with KS3AT3, ACP3, and ACP2(2), and the major protein complex was isolated by SEC for single-particle cryo-EM analysis (Fig. S12).

**Fig. 2.**
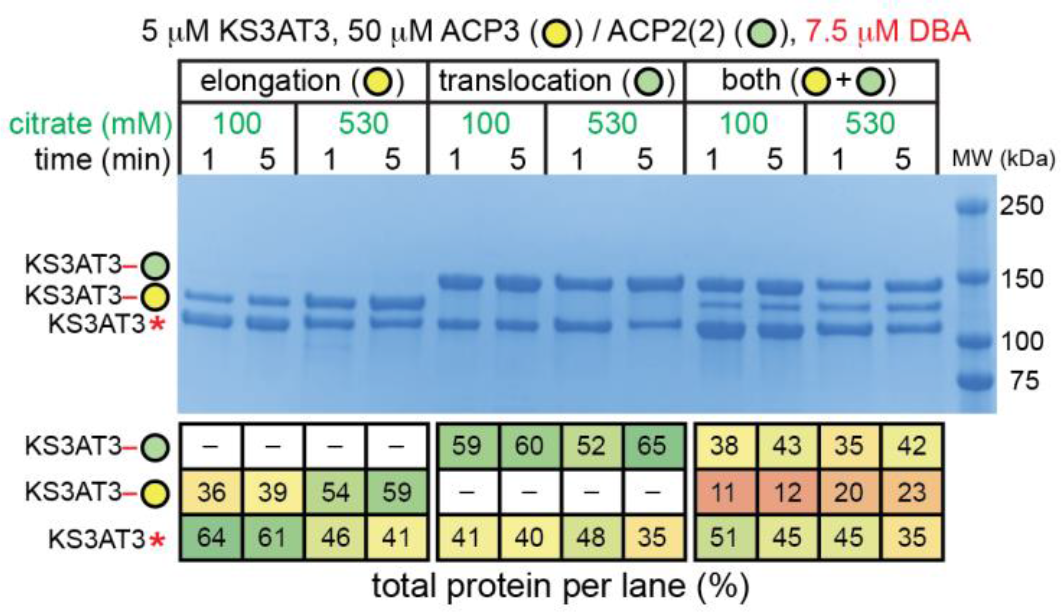
SDS-PAGE analysis of DBA crosslinking between the homodimeric KS-AT core of Module 3 and its *holo-*form elongation (ACP3) and translocation (ACP2(2)) ACP partners. Total protein per lane was quantified and expressed as an average percentage of two technical replicates (Supplementary Raw Data 2; standard deviation ≤2%). Red asterisks denote a protein whose KS active site harbors a quenched DBA moiety (as explained in Fig. S5A). For additional controls associated with both crosslinking reactions, see Figs. S10 and S11. For gel source data, see Supplementary Raw Data.

Three-dimensional classification and refinement of cryo-EM data from 7,500 dose-fractioned movies produced a noteworthy map at 3.05 Å gold-standard Fourier shell correlation (GSFSC) resolution^22^ from ∼25% (77,996) of the total particles (Figs. S13– S14). This map, which featured unambiguous density for one copy of ACP2(2), represents the first near-atomic resolution structure of a catalytically competent module-module interface from an assembly-line PKS. In this structure, ACP2(2) is bound to the KS active site cleft in an orientation that is entirely supportive of previous predictions based on mutational analysis, solution NMR, and *in silico* docking^23–25^ (Fig. 3A). Specifically, helix 1 of ACP2(2) forms multiple electrostatic contacts with the proximal AT domain (Fig. 3B). Although the region of the DBA-derived crosslink was not visible, the Ppant cofactor could be partially traced in the KS active site (Figs. 3B and S15). Notably, the map resolved the helical portion of the C-terminal docking domain of ACP2(2) engaged in hydrophobic and electrostatic interactions with the N-terminal docking domain of KS3AT3 (Fig. 3C). A short α-helical segment (11 residues) of ACP2(2) lies parallel in the groove of the α-coiled-coil docking domain of KS3AT3 to form a three-helix bundle, consistent with the solution NMR structure of a similar docking domain complex^20^. Unlike the NMR structure, however, which captured a symmetric four-helix bundle through covalent fusion of the interacting polypeptides^26^, our cryo-EM structure revealed an asymmetric complex in which only a single donor ACP interacts with its homodimeric acceptor KS partner (Fig. 3A, C). While statistical image analysis (discussed below) revealed the presence of homodimeric KS3AT3 particles in which both KS-AT clefts were occupied by ACP partners, none of these higher-order complexes could be refined to yield near-atomic resolution structures.

**Fig. 3.**
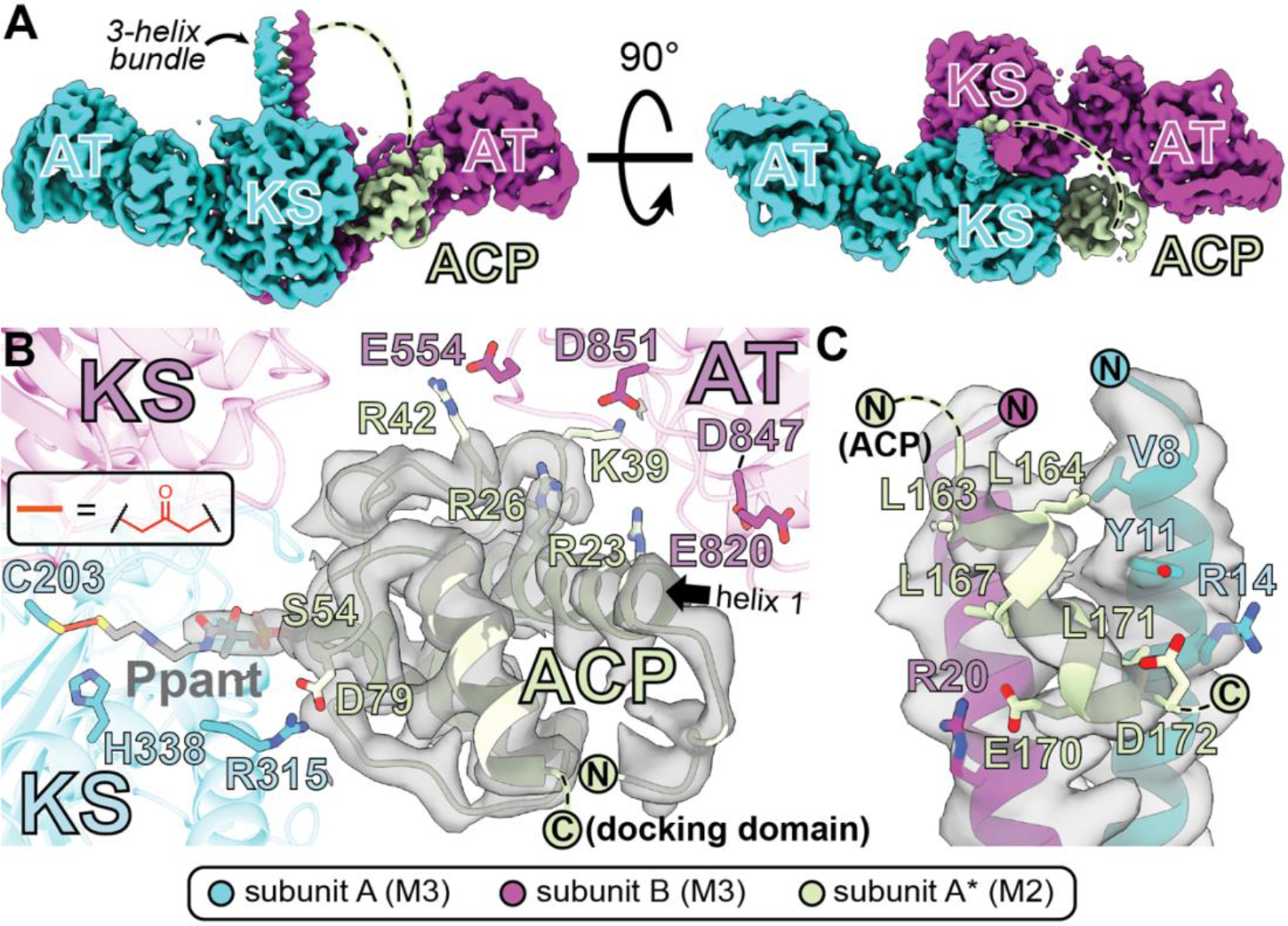
(**A**) Cryo-EM structure of an intermodular docking interaction during polyketide translocation at 3.05 Å GSFSC resolution represented by two 90°-related orientations of the homodimeric KS3AT3 fragment of DEBS crosslinked to its upstream ACP partner, ACP2(2) (*Translocation*, Figs. S13–S14). Interactions involving ACP2(2) at the (**B**) KS-AT cleft and (**C**) N-terminal docking domain of KS3AT3 are explicitly shown (67 residues separating the ACP and its C-terminal docking domain were unresolved; depicted as a dashed line). In panel **B**, the cryo-EM map surface is displayed within a 2 Å radius relative to ACP2(2) atoms. Prior mutagenesis analysis established that residues on helix 1 (black arrow) were critical for polyketide translocation^24^. This structure reveals that Arg23 and Arg26 on helix 1 are appropriately situated to form salt bridges with Glu820, Asp847, and Asp851 in the AT domain of KS3AT3. Based on proximity to the observed backbone atoms, additional interactions are likely to occur between Lys39 and Arg42 of ACP2(2) and Glu554 (from the KS-AT linker) and Asp851 (from the AT domain) of KS3AT3.

Our description of an asymmetric state for intermodular translocation of the growing polyketide chain contrasts with a previously reported symmetric state for the same reaction in a different assembly-line PKS^8^. Further studies will be required to establish whether these differences stem from technical considerations or if they reflect mechanistic divergence between related assembly lines.

### Cryo-EM analysis of crosslinked DEBS Modules 1 and 3

In addition to enabling structural characterization of the ACP2(2)–KS3AT3 complex, DBA crosslinking also yielded preparations of modules M1TE and M3TE bound to a monoclonal antibody fragment (F_ab_) 1B2^12^ that were suitable for cryo-EM analysis (Fig. S12). 1B2 did not affect the formation of bands **1** or **2** when included in DBA crosslinking reactions (Fig. S16). A total of 9,523 and 10,480 dose-fractionated movies were collected for crosslinked M1TE and M3TE, respectively. Following particle picking and 2D classification, *ab initio* reconstruction without imposed symmetry yielded 3D volumes resembling the previous cryo-EM structures of DEBS M1TE and a hybrid module derived by fusing fragments of M3, M1, and the TE domain (M3/1TE)^12,27^. Crosslinked M1TE contained two bound copies of 1B2 oriented symmetrically about the pseudo-C2 axis of symmetry, whereas crosslinked M3TE contained only a single 1B2 — akin to the previous structure of M3/1TE bound to 1B2 (Figs. S17–S22). (We previously ascribed this difference to a 12° bending of the N-terminal coiled-coil docking domain relative to the pseudo-C2 axis of symmetry that precluded binding of the second F_ab_ ^12^.) *Ab initio* classes were refined to yield consensus cryo-EM maps of crosslinked M1TE-1B2 and M3TE-1B2 with GSFSC resolutions of 3.18 Å each (Figs. S17–S22). Overall, significant conformational heterogeneity was observed in the C-terminal fragment of each module (harboring the KR, ACP, and TE domains) (Figs. S17–S22).

Cryo-EM maps of crosslinked M1TE revealed ACP density in the cleft between the KS and AT domains (Fig. S17), reminiscent of the *State 1* pre-elongation mode^12^. Also consistent with our earlier characterization of *State 1* of this module, no more than one bound ACP was observed in any of the 3D classes. From these data it also follows that, if a subunit harboring an intra-polypeptide crosslink (i.e., band **2**; Fig. 1B) exists in a dimeric state, then it must partner with an un-crosslinked subunit (i.e., band **3**; Fig. 1B).

To identify which cryo-EM maps of crosslinked M1TE corresponded to bands **1**–**3** (Fig. 1B), we inspected maps with observable ACPs to assign *inter*- or *intra*-molecular KS-ACP interactions. One map (3.73 Å) revealed an intermolecular KS-ACP linkage and presumably corresponded to band **1** (Fig. S23B). It also showed strong similarity to *State 1* described earlier and was therefore assigned as *Crosslinked State 1*. Another map (3.61 Å) supported an intramolecular KS-ACP linkage whose subunit was partnered with an un-crosslinked subunit (Fig. S23B), suggesting that band **2** and band **3** were in fact present in the same dimer (Fig. 1B).

Significant disorder in the regions connecting the KS-AT core of crosslinked M3TE to its ACP domain precluded assignment of its cryo-EM maps to bands **1**–**3**. Nonetheless, a single ACP clearly docking onto the active site of the KS could also be observed in crosslinked M3TE (Fig. 4A–C). Importantly, Arg1439, Arg1440, and Glu1444 from ACP3 interacted with reciprocally charged residues Glu554, Glu518, and Arg480, respectively. The latter set of residues map to the KS-AT linker subdomain (Fig. 4C–D). Prior mutagenesis studies have established that these ACP residues play pivotal roles in controlling KS-ACP specificity during chain elongation (Fig. 4E)^24^. (See Table S1 for a summary of structures and their compositions from this study and our related previous study^12^.)

**Fig. 4.**
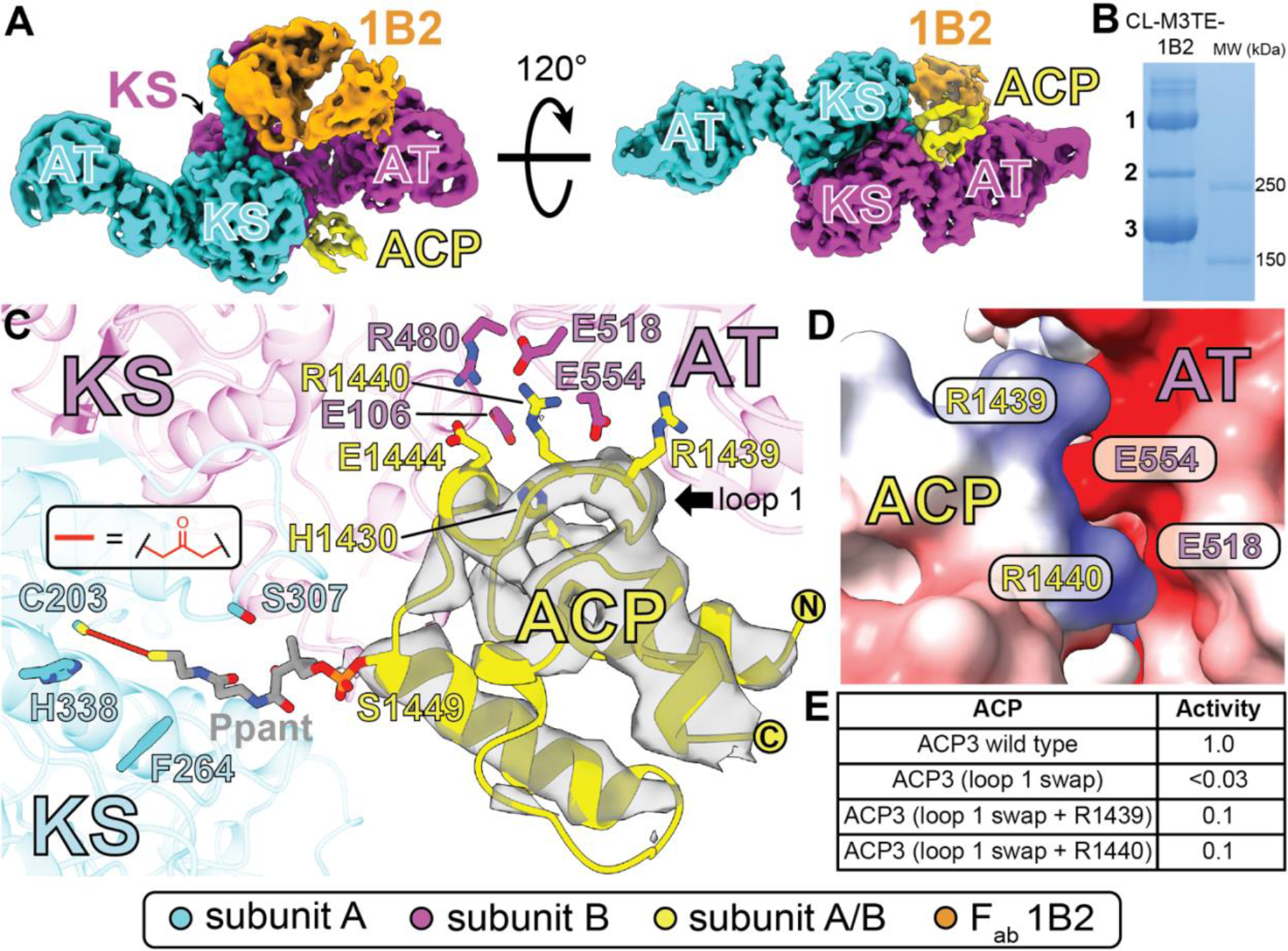
Structural analysis of crosslinked DEBS M3TE in complex with F_ab_ 1B2 (CL-M3TE-1B2). (**A**) The structure deduced from one map (designated *Cis F*_*ab*_*/ACP*; Figs. S20–S21) at 3.46 Å GSFSC resolution is shown in two 120°-related orientations. A similar cryo-EM map (designated *Trans F*_*ab*_*/ACP*; Figs. S20 and S22) was obtained at 3.40 Å GSFSC resolution; it contains an ACP bound to the opposite KS, indicating that F_ab_ 1B2 does not affect ACP binding. (**B**) SDS-PAGE analysis of the CL-M3TE-1B2 sample containing bands **1**–**3** (Fig. 1B) that was used for structural analysis (Fig. S12). (**C**) Close-up view of the interactions between ACP3 and the KS-AT cleft of M3TE. Based on proximity to the observed backbone atoms, electrostatic bonds are likely to occur between the ACP and the KS-AT interdomain linker from the opposite subunit to which the ACP3 is covalently attached (magenta versus cyan). The cryo-EM map surface is displayed within a 3 Å radius relative to the ACP atoms, and density for the Ppant group is not observed at the set threshold value. (**D**) Complementary electrostatic surfaces on the ACP (yellow) and KS-AT (magenta) are shown. (**E**) Variants of ACP3 were previously tested for their ability to facilitate polyketide chain elongation in partnership with the KS3AT3 didomain^24^. Almost all detectable activity was lost when the loop 1 of ACP3 (black arrow) was replaced with its counterpart from ACP6. However, when R1439 or R1440 were reintroduced into this construct individually, partial activity was restored. For gel source data, see Supplementary Raw Data.

### Probing the origin of partial occupancy during polyketide chain elongation

The repeated observation that only one KS-ACP interaction at a time is permitted in a homodimeric module led us to ask whether *in trans* crosslinking of the homodimeric KS-AT core of M3 (KS3AT3) with its elongation partner ACP3 would also be subject to this stoichiometric limit. As shown in Figure S11, DBA crosslinking of KS3AT3 and ACP3 under a range of ACP3 and citrate concentrations was unable to yield stoichiometric crosslinking of the two proteins. Cryo-EM analysis of the SEC-purified complex (Fig. S12) produced a consensus cryo-EM map at 3.71 Å GSFSC resolution (Figs. S24–S25). It revealed the expected structural features of an extended KS3AT3 homodimer along with weak density corresponding to the ACP bound at each of the KS active site clefts (not visual at the threshold values used in Figs. S24–S25).

To quantify the relative abundance of particles containing 0, 1, or 2 ACPs, the data were subjected to a recently described statistical image analysis methodology for measuring intersubunit variability at the level of individual particles^28^. Specifically, each pseudo-C2-symmetric particle was segmented into two asymmetric subunits by C1 to C2 symmetry expansion. Individual C2-segmented particles were then sorted into two 3D classes according to differences in ACP occupancy by employing a focused mask around their ACP-binding sites. As expected, one 3D class revealed clear and unambiguous density for ACP3 in the ACP-binding site, whereas the other was completely devoid of density in this site (Fig. 5A). Particles belonging to each class were then recombined with their symmetry mates to generate the intact C1-particles and then classified as KS3AT3 homodimers bound to 0, 1, or 2 ACP subunits. Strikingly, a near 1:2:1 particle ratio was observed between 3D classes with 0 (25.0%), 1 (50.3%), or 2 (24.7%) ACPs (Fig. 5A), indicating that docking of the elongation ACP partner occurred stochastically in the dissociated (i.e., KS-AT + ACP) state of a PKS module. In contrast, when an identical image analysis protocol was employed to quantify ACP occupancy of the crosslinked form of intact M1TE, only particles with 1 ACP were observed (Figs. 5B and S26). Thus, covalent tethering of an ACP to the catalytic KS-AT core of a homodimeric PKS module precludes both KS active sites from catalyzing chain elongation at the same time.

**Fig. 5.**
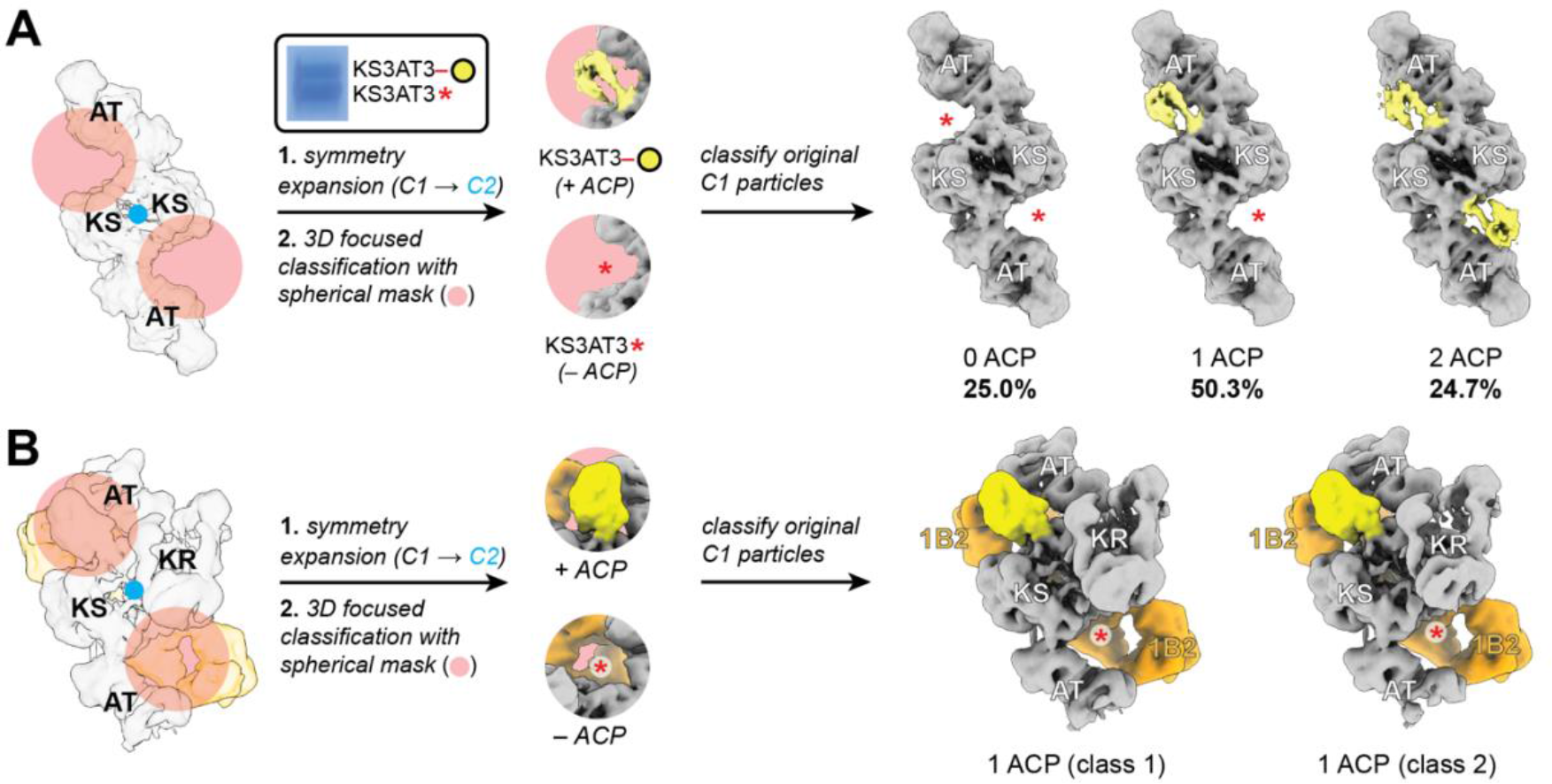
The ACP occupancy of KS-AT homodimers was measured by a recently developed method (statistical per-particle image analysis^28^) following crosslinking of (**A**) KS3AT3 + ACP3 (yellow circle) or (**B**) intact Module 1. The boxed inset in panel **A** shows SDS-PAGE analysis of the crosslinked material isolated as a single peak by SEC for cryo-EM analysis (Fig. S12). (**A, B**) Consensus particles were (**1**) symmetry-expanded about the pseudo-C2 axis of symmetry (blue dot), and (**2**) sorted according to differences in density at the ACP binding sites by 3D focused classification. Particles from each class were correlated to their C2 symmetry mates to categorize intact particles as bound to 0, 1, or 2 ACPs (see Methods). As elaborated in Fig. S26, slight differences in the ‘1 ACP’ classes in panel **B** resulted in their segregation during 3D focused classification. Red asterisks denote KS active sites harboring a quenched DBA moiety (Fig. S5A). For gel source data, see Supplementary Raw Data.

Our observation that no more than one KS domain of a homodimeric module can crosslink with its elongation ACP partner lends credence to a previous proposal that both ACPs co-migrate between two equivalent “reaction chambers” during the catalytic cycle^13^. To explain this phenomenon, Bagde et al. pointed out that C-terminal dimeric motifs — which are frequently encountered in assembly-like PKS modules — might hold the ACPs in proximity such that only one of them can partake in elongation at a given time. It should be noted that our crosslinking analysis was performed on PKS modules bearing C-terminal TE domains that are homodimeric^29^.

### Statistical image analysis of a module-module interface during polyketide translocation

Compelled by successful application of statistical per-particle image analysis to cryo-EM data for the crosslinked product of KS3AT3 and ACP3, we also sought to establish whether two upstream (translocation) ACP partners or a translocation and an elongation ACP partner could simultaneously dock onto the homodimeric KS-AT core of a module. To do so, the cryo-EM data from Fig. S13 were subjected to the same computational protocol outlined above, modified slightly to allow for the simultaneous docking of two identical or different ACPs onto the KS-AT homodimer. Overall, no homo- or hetero-dimeric states were forbidden in our analysis (Fig. S27). Thus, in contrast to the inability of the KS-AT core of a homodimeric PKS module to accommodate both elongation ACP domains simultaneously, there does not appear to be a barrier to concomitant chain translocation into both KS active sites or to synchronous translocation and elongation reactions. Our results, however, do not distinguish how a PKS module interacts with its ACPs in the presence of protein tethered intermediates during the catalytic cycle.

## DISCUSSION

Since the first report of a selective protein-protein interaction at the interface between two successive modules of an assembly-line PKS^19^, there has been a steady growth of evidence that protein-protein interactions play dominant roles in channeling reactive intermediates along the enzymatic assembly line^30^. Our cryo-EM structure of ACP2(2) docked to the KS-AT core of Module 3 of DEBS (Fig. 3) represents the first structure of an inter-modular interface in an assembly-line PKS at near-atomic resolution, thereby establishing a foundation for the structure-based engineering of hybrid assembly lines. Encouragingly, the observed KS-AT/ACP interfaces involving both ACP2(2), which participates in chain translocation to Module 3, and ACP3, which participates in chain elongation catalyzed by Module 3, are well supported by empirical data from mutagenesis and other approaches^23–25^. While these earlier studies flagged the ACP residues interacting with the KS-AT core, the corresponding residues on the KS-AT side of the interface remained unknown until now.

The distinct binding modes through which ACPs are recognized during polyketide chain translocation and elongation can be readily visualized by comparing our structures of DEBS Module 3 (Fig. S28A–B). Whereas ACP3 binds a relatively small portion of the KS-AT cleft during intra-modular elongation, ACP2(2) forms extensive interactions in the cleft while forming secondary contacts with the docking domain during inter-modular translocation. In this case, tighter recognition of the upstream ACP across a module-module junction is probably a reflection of the increased demand to overcome the entropic burden of a bimolecular substrate transfer. In fact, a recent study found that KS domains from PKS assembly lines genetically co-migrate with their upstream, not downstream, ACP partners^31,32^. While these evolutionary insights reinforce the importance of the ACP-KS partnership during translocation — and can be harnessed to pinpoint appropriate cut-sites for PKS module exchange^18,33,34^ — our 3.05 Å structure of a module-module interface provides the molecular blueprints from which to rationally build and test hybrid assembly lines capable of generating new-to-nature polyketides. (For a summary of the structures of DEBS reported here and previously^12^, see Table S1 and Fig. S28.)

Our cryo-EM analysis of DEBS has also underscored the importance of structural asymmetry in a homodimeric PKS module as it progresses through its catalytic cycle, while shining new light on its mechanistic origins. Notably, the putatively symmetric module state, where both elongation ACP partners are simultaneously docked to the KS-AT core of the module, is strongly disfavored. In contrast, when the KS-AT core of a homodimeric module is genetically dissociated from its elongation ACP partner, the dual-occupancy state is readily observed. Similarly, simultaneous crosslinking of one KS domain to its elongation ACP partner and the other KS to its translocation ACP partner is also feasible, as is the symmetric dual-occupancy state where both translocation ACP partners simultaneously dock onto their downstream module. It must however be recognized that DBA crosslinking is incapable of capturing module states whose formation depends on exergonic chain elongation. Such states have in fact been previously described^35^ and characterized by cryo-EM^12^.

Based on all available data thus far, we have therefore proposed the existence of a coupling mechanism between the exergonic C-C bond formation and its two associated thermoneutral (thiol-to-thioester exchange) translocation reactions that collectively define the catalytic cycle of each module of an assembly-line PKS (Fig. S29; Ref. 36). Not only can such coupling be expected to enforce directionality of polyketide biosynthesis along the assembly line, but it would also explain why disruptions to KS-AT/ACP interfaces can have profoundly negative kinetic consequences.

## Conclusion

The hallmark of an assembly-line PKS lies in the ability of each of its constituent enzymatic modules to toggle between *intra*modular elongation and *inter*modular translocation reactions involving the growing polyketide chain. Here we have described an asymmetric state associated with the latter reaction in which only one monomer of a homodimeric acceptor module engages with a single upstream ACP at a time. Taken together with our recent structural characterization of a module involved in the former reaction^12^, the catalytic relevance of asymmetry in KS-ACP interactions is clear. Our present report also includes two methodological advances — the use of DBA as a bifunctional crosslinker to stabilize transient PKS states for structural analysis; and the use of statistical per-particle image analysis of cryo-EM data to analyze asymmetric conformations of C_2_-symmetric PKS homodimers. We anticipate broad use of both methods in future studies of polyketide biosynthetic mechanisms.

## Supporting information

Supplementary Raw Data

## Author contributions

D.P.C., A.M.S., M.C., and C.K. conceived of the project aims. D.P.C. and A.M.S. collected non–cryo-EM experimental data. D.P.C. and Y.L. performed single-particle cryo-EM analysis. D.P.C. and K.L.B. refined the atomic models. M.C. performed the statistical per-particle image analysis. C.K. supervised all experiments. D.P.C. and C.K. wrote the initial manuscript which was revised and edited by all authors.

## Acknowledgements

The authors would like to thank Wah Chiu (Stanford University) for helpful discussions during the preparation of this manuscript.

## Data and materials availability

All atomic coordinates and cryo-EM maps have been deposited in the Protein Data Bank under accession codes 8TJN, 8TJO, 8TPW, 8TPX, 8TJP, and 8TKO and in the Electron Microscopy Data Bank under accession codes EMD-41305, EMD-41306, EMD-41495, EMD-41496, EMD-41307, and EMD-41355. All materials used in this study that are not commercially available can be made available by the authors upon request.

