## Supplementary Information for "Structural Basis for Intermodular Communication in Assembly-Line Polyketide Biosynthesis"

Materials and Methods ........ 2–9
Protein Sequences ........ 10–12
Supplementary Figures ........ 13–47
- Fig. S1: The 6-deoxyerythronolide B synthase (DEBS) ........ 13
- Fig. S2: The catalytic cycle of DEBS Module 1 ........ 14–15
- Fig. S3: SDS-PAGE analysis of 1,3-dibromoacetone (DBA)-mediated crosslinking of DEBS Modules 1 and 3 and time-course of the reaction ........ 16–17
- Fig. S4: DBA crosslinking of DEBS Module 1 and KS-AT fragment of Module 3 (KS3AT3) depends on both KS and ACP thiols ........ 18–19
- Fig. S5: Putative structures of DBA-crosslinked DEBS modules ........ 20
- Fig. S6: DBA crosslinking of a fluorescent probe, ACP2(2)-**FL**, with DEBS KS3AT3 ........ 21
- Fig. S7: DBA crosslinking of ACP2(2)-**FL** with DEBS Module 3 ........ 22
- Fig. S8: *A priori* trypsinolysis of putative DBA-crosslinked adducts ........ 23
- Fig. S9: Limited trypsinolysis of DBA-crosslinked DEBS Module 3 ........ 24
- Fig. S10: DBA-mediated crosslinking between DEBS KS3AT3 didomain and *holo*- versus *apo*-form ACP2(2) ........ 25-26
- Fig. S11: DBA-mediated crosslinking between DEBS KS3AT3 didomain and *holo*- versus *apo*-form ACP3 ........ 27–28
- Fig. S12: Size-exclusion chromatography (SEC) and SDS-PAGE analysis of samples used for single-particle cryo-EM ........ 29–30
Supplementary References.........................................................................52–53

### Materials and Methods

##### Materials

Bacterial growth media, purification resins, Amicon Ultra Centrifugal Filters, and other chemicals were purchased from Thermo Fisher Scientific or MilliPore Sigma. Antibiotics and isopropyl β-D-1-thiogalactopyranoside (IPTG) were purchased from Gold Biotechnology. 1,3-dibromoacetone (DBA) was purchased from Santa Cruz Biotechnology (sc-208768) or Alfa Aesar (H37325). Cryogenic electron microscopy (cryo-EM) Quantifoil R 2/1 300 mesh Cu grids were purchased from Electron Microscopy Sciences (Q3100CR1). NuPAGE 3–8% Tris-Acetate (3–8% TA) polyacrylamide gels (Invitrogen) were purchased from Thermo Fisher Scientific (EA0375PK2) and 4–20% Mini-PROTEAN TGX (4–20% TGX) polyacrylamide gels were purchased from Bio-Rad (#4561096).

##### General Methods

Protein affinity and size-exclusion chromatographies were carried out using an ÄKTA Pure chromatography system (Cytiva Life Sciences). The following proteins were expressed and purified for use in bimodule enzyme assays, as previously[1,2]: LDD(4), (5)M1(2), and (3)M2TE from the 6-deoxyerythronolide B synthase (DEBS; parenthetical numbers refer to native docking domains and the modules from which they derive);

auxiliary proteins MatB, PrpE, and methylmalonyl-CoA epimerase (SCME). To generate the ACP2(2)-**FL** probe, *apo*-ACP2(2) and Sfp were prepared as previously[3–5]. Protein concentrations were determined using the Bradford assay with bovine serum albumin standards by measuring absorbance at 595 nm in 1 cm cuvettes using a NanoDrop 2000c (Thermo Fisher Scientific)[6]. Plasmids used for protein expression are listed in Table S1. Unless otherwise stated, n=1 for all crosslinking and proteolysis experiments.

**Production of plasmid pDC9 for overexpression of DEBS ACP3**

DNA oligos 5'-TGGTGCCGCGCGGCAGCCATGGGCTTTCCCCGGACGAGCAGCAG-3' and 5'-CCGCAAGCTTGTCGACGGAGTTATTTGACGAGCCGGGCGCGCAGGT-3' were used to PCR amplify the ACP3 encoding region of *eryAII* with template pFW98[1]. A pET28 backbone was meanwhile amplified using DNA oligos 5'-CTCCGTCGACAAGCTTGCGGCCG3' and 5'ATGGCTGCCGCGCGGCACCAG-3' for two-component Gibson Assembly[7]. See **Protein Sequences** for the translated ACP3 product of this plasmid.

**Protein Expression and Purification**

<u>DEBS proteins</u>: Expression plasmids (see Table S1) were used to transform *Escherichia coli* BAP1 or BL21(DE3) competent cells[8] to generate proteins in their *holo* or *apo* states, respectively. Single colony transformants were selected to inoculate 20 mL Luria-Bertani (LB) media supplemented with appropriate antibiotic (i.e., 100 µg/mL carbenicillin or 50 µg/mL kanamycin; see Table S1) and grown overnight at 37 °C (220 rpm). Two mL of overnight seed culture were used to inoculate 1 L antibiotic-containing LB medium and grown at 37 °C (220 rpm) until an optical density at 600 nm ($OD_{600}$) of ~0.5 was achieved (4–8 L total). Cultures were cooled for 20 min in an ice bath before adding 0.25 mM IPTG and continuing growth at 18 °C (220 rpm) for 18 h. Cell harvesting, lysis, Ni-NTA affinity purification, and anion exchange chromatography was performed as previously[9,10]. Eluted fractions from anion exchange chromatography were evaluated for purity by SDS-PAGE before pooling and concentrating the protein to ≤1 mL using Amicon Ultra Centrifugal Filters (50 kDa MWCO) and flash-freezing in liquid $N_2$ for storage at -80 °C. DEBS modules were purified by size-exclusion chromatography (SEC) by injecting ≤2 mL of anion-exchange purified material onto a 120 mL Superdex 200 pg 16/600 column (Cytiva Life Sciences) and eluting isocratically with 0.1 M citric acid, 0.1 M NaCl, 10 mM HEPES, pH 7.2 [NaOH] (SEC buffer) at a flow rate of 1.5 mL/min. Fractions containing pure protein as assessed by SDS-PAGE were pooled and concentrated using Amicon Ultra Centrifugal Filters (Millipore Sigma).

<u>$F_{ab}$ 1B2</u>: Expression, periplasmic extraction, and Ni-NTA affinity purification of $F_{ab}$ 1B2 was performed as previously[9]. Purified protein was concentrated to ≤1 mL using Amicon Ultra Centrifugal Filters (10 kDa MWCO), supplemented with 5% glycerol, and flash-frozen in liquid $N_2$ for storage at -80 °C. 1B2 was further purified by SEC in a similar manner as above for DEBS modules, and fractions corresponding to the heterodimeric species (~83 mL retention volume; Fig. S12F) were pooled and concentrated using Amicon Ultra Centrifugal Filters (10 kDa MWCO; Millipore Sigma).

**DBA Crosslinking Reactions**

General DBA crosslinking (Figs. 1B and S3A): Prior to use in crosslinking reactions, DBA was filtered through a small plug (~1 mL) of anhydrous aluminum oxide under argon gas and diluted to 0.5 M in anhydrous DMF for storage at -80 °C. DBA was then diluted to 0.25 mM in 50% aqueous DMF for initiating crosslinking reactions no more than 15 min prior. In three identical replicates, SEC-purified DEBS M1TE (see **Protein Sequences**) (10 μM) was incubated in 10 mM 4-(2-hydroxyethyl)-1-piperazineethanesulfonic acid (HEPES), 100 μM tris(2-carboxyethyl)phosphine (TCEP), and 200 mM citric acid, pH 7.2 [NaOH] at 20 °C for 15 min before adding 12.5 μM DBA (reaction volume = 25 μL; reaction concentrations reflect their final concentrations after all components were added). Reactions were quenched after 30 s by the addition of an equal volume of 2x Laemmli buffer (Bio-Rad #1610737) supplemented with 5% (*v/v*) β-mercaptoethanol (BME) and heated at 95 °C for 2 min prior to SDS-PAGE analysis, using either 3–8% TA or 4–20% TGX polyacrylamide gels (3–8% TA gels provided better separation between bands **1** and **2** than 4–20% TGX gels — see Figs. S4A,B and S3 for comparison).

Kinetics of DBA crosslinking and citrate dependence thereof (Fig. S3B): SEC-purified DEBS M3TE (5 μM) was incubated in 10 mM HEPES, 100 μM TCEP, and 50 mM, 200 mM, or 500 mM citric acid, pH 7.2 [NaOH] at 20 °C for 15 min before adding 7.5 μM DBA (reaction volume = 25 μL; reaction concentrations reflect their final concentrations after all components were added). Reactions were quenched at three different time points (5 s, 1 min, or 25 min) by the addition of an equal volume of 2x Laemmli buffer (Bio-Rad #1610737) supplemented with 5% (*v/v*) BME and heated at 95 °C for 2 min prior to SDS-PAGE analysis.

Requirement of an ACP domain for DBA crosslinking of M1TE (Fig. S4A): DEBS M1TE prepared in its *apo* and *holo* states (see **Protein Expression and Purification**) was used in side-by-side DBA crosslinking reactions to measure the dependence of bands **1** and **2** formation on the presence of the 4′-phosphopantetheine (Ppant) cofactor of the ACP. M1TE (10 μM) was incubated in 10 mM HEPES, 100 μM TCEP, and 200 mM citric acid, pH 7.2 [NaOH] at 20 °C for 15 min before adding 6.3 μM, 12.5 μM, or 25 μM DBA (reaction volume = 20 μL; reaction concentrations reflect their final concentrations after all components were added). Reactions were quenched after 0.5 min, 5 min, or 15 min by the addition of an equal volume of 2x Laemmli buffer (Bio-Rad #1610737) supplemented with 5% (*v/v*) BME and heated at 95 °C for 2 min prior to SDS-PAGE analysis.

Requirement of a KS domain for DBA crosslinking of M1TE (Fig. S4B): DEBS M1TE (31 μM) was incubated with 3.1 mM cerulenin or DMSO (control) in 0.1 M $NaH_2PO_4$, pH 8 [NaOH] at 20 °C for 1 hour. Residual cerulenin and DMSO were removed using 7 kDa MWCO Zeba spin desalting columns (Thermo Fisher: 89882) equilibrated with 10 mM HEPES, 200 mM citric acid, pH 7.2 [NaOH]. Cerulenin treated and untreated M1TE (10 μM) were incubated in 10 mM HEPES, 100 μM TCEP, and 200 mM citric acid, pH 7.2 [NaOH] at 20 °C for 15 min before adding 6.3 μM, 12.5 μM, or 25 μM DBA (reaction volume = 10 μL; reaction concentrations reflect their final concentrations after all components were added). Reactions were quenched after 1.2 min or 8 min by the addition of an equal volume of 2x Laemmli buffer (Bio-Rad #1610737) supplemented with 5% (v/v) BME and heated at 95 °C for 2 min prior to SDS-PAGE analysis. To ensure that cerulenin inhibits M1, DEBS (5)M1(2) was treated with cerulenin or DMSO (control) and desalted in the same manner as for M1TE.

Requirement of both KS and ACP domains for DBA crosslinking of KS3AT3 (Fig. S4C): Wild-type or C202A variant KS3AT3 (5 μM) and *apo*- or *holo*-form ACP2(2) (25 μM) were incubated in 10 mM HEPES, 100 μM TCEP, and 200 mM citric acid, pH 7.2 [NaOH] at 20 °C for 15 min before adding 10 μM DBA (reaction volume = 20 μL; reaction concentrations reflect their final concentrations after all components were added). Reactions were quenched after 1 min by the addition of an equal volume of 2x Laemmli buffer (Bio-Rad #1610737) supplemented with 5% (v/v) BME and heated at 95 °C for 2 min prior to SDS-PAGE analysis.

DBA crosslinking of M3TE with fluorescent ACP2(2) (Fig. S7): See below for the preparation of the fluorescein labeled ACP2(2) probe, ACP2(2)-**FL**. SEC-purified DEBS M3TE (5 μM) was incubated in 10 mM HEPES, 100 μM TCEP, and 150 mM, 260 mM, or 510 mM citric acid, pH 7.2 [NaOH] at 20 °C for 15 min before adding 10 μM DBA (reaction volume = 20 μL; reaction concentrations reflect their final concentrations after all components were added). Reactions were quenched after 4.5 min by the addition of an equal volume of 2x Laemmli buffer (Bio-Rad #1610737) supplemented with 5% (v/v) BME and heated at 95 °C for 2 min prior to SDS-PAGE analysis. In-gel fluorescence measurements were made using a ChemiDoc MP Imaging System (Bio-Rad) set to the fluorescein excitation and emission settings.

Preparation of crosslinked modules and KSAT-ACP complexes for single-particle cryo-EM analysis: To scale-up the crosslinked M1TE and M3TE products for single-particle cryo-EM analysis, we prepared 5–16 replicate DBA crosslinking reactions and pooled them after quenching to avoid potential mixing effects on crosslinking efficiency due to increased reaction volume. Each reaction was carried out as above (see "General DBA crosslinking") with slight adaptations. That is, M1TE or M3TE (40 μM) were incubated with 10 mM HEPES, 200 μM TCEP, and 200 mM citric acid, pH 7.2 [NaOH] for 15 min before adding 50 μM DBA (reaction volume = 60 μL x 5, reaction scale = 12 nmol). KS3AT3 (20 μM) and ACP3 (235 μM) were incubated with 10 mM HEPES, 100 μM TCEP, and 450 mM citric acid, pH 7.2 [NaOH] for 15 min before adding 30 μM DBA (reaction volume = 60 μL x 16, reaction scale = 19.2 nmol). Finally, KS3AT3 (20 μM), ACP2(2) (100 μM), and ACP3 (200 μM) were incubated with 10 mM HEPES, 100 μM TCEP, and 450 mM citric acid, pH 7.2 [NaOH] for 15 min before adding 30 μM DBA (reaction volume = 60 μL x 16, reaction scale = 19.2 nmol). Reactions were quenched after 30 s by the addition of an equal volume of 2x Laemmli buffer (Bio-Rad #1610737) supplemented with 5% (v/v) BME and heated at 95 °C for 2 min prior to SDS-PAGE analysis. Reaction concentrations reflect their final concentrations after all components were added. Whereas crosslinked M1TE and M3TE were SEC-purified in 0.1 M citric acid, 0.1 M NaCl, 10 mM HEPES, pH 7.2 [NaOH] (Fig. S12A, B), crosslinked KS3AT3-ACP3 and KS3AT3-ACP2(2)/ACP3 were SEC-purified in 0.3 M citric acid, 0.1 M NaCl, 10 mM HEPES, pH 7.2 [NaOH] (Fig. S12C–E).

Crosslinking of KS3AT3 with *holo*- and *apo*-form ACP3 (Fig. S11): SEC-purified DEBS KS3AT3 (5 μM) was incubated with 5 μM, 50 μM, or 250 μM *holo*- or *apo*-form ACP3 in 10 mM HEPES, 100 μM TCEP, and 50 mM, 200 mM, or 530 mM citric acid, pH 7.2 [NaOH] at 20 °C for 15 min before adding 7.5 μM DBA (reaction volume = 20 μL; reaction concentrations reflect their final concentrations after all components were added). Reactions were quenched at two different time points (2 and 10 min) by the addition of an equal volume of 2x Laemmli buffer (Bio-Rad #1610737) supplemented with 5% (*v/v*) BME and heated at 95 °C for 2 min prior to SDS-PAGE analysis. A control reaction was also included that was prepared in the same manner as above; however, *apo*-ACP3 (0.5 mM) was first treated with 1 mM bismaleimidoethane (BMOE) for 15 min in 200 mM citric acid, pH 7.2 [NaOH] followed by addition of 3 mM BME and buffer exchanged using a 7 kDa MWCO Zeba spin desalting column (Thermo Fisher: 89882) equilibrated with a similar buffer ("Control" in Fig. S11).

Crosslinking of KS3AT3 with *holo*- and *apo*-form ACP2(2) (Fig. S10): ACP2(2) was prepared in three different ways prior to crosslinking with KS3AT3: (1) *apo*-form ACP2(2) (500 μM) was incubated with 4 mM CoA, 10 mM $MgCl_2$, and 50 μM Sfp in 0.1 M $NaH_2PO_4$ pH 7.2 [NaOH] for 1 hour (reaction volume = 25 μL; reaction concentrations reflect their final concentrations after all components were added) before buffer exchanging using a 7 kDa MWCO Zeba spin desalting column (Thermo Fisher: 89882) equilibrated with 0.1 M $NaH_2PO_4$ pH 7.2 [NaOH] to afford the *holo*-form ACP2(2) "ACP2(2) / Sfp + CoA;" (2) *apo*-form ACP2(2) (500 μM) was incubated with 10 mM $MgCl_2$, 50 μM Sfp, and without exogenous CoA in 0.1 M $NaH_2PO_4$ pH 7.2 [NaOH] for 1 hour (reaction volume = 25 μL; reaction concentrations reflect their final concentrations after all components were added) before buffer exchanging using a 7 kDa MWCO Zeba spin desalting column (Thermo Fisher: 89882) equilibrated with 0.1 M $NaH_2PO_4$ pH 7.2 [NaOH] to afford the *apo*-form ACP2(2) "ACP2(2) / Sfp  CoA;" (however, this sample was later determined to be not entirely *apo* due to co-purification of Sfp with CoA from *E. coli* BL21(DE3); Fig. S10); (3) a final form of *apo*-ACP2(2) was not treated with any of the above reaction components as in (1) and (2) and used directly in DBA crosslinking reactions with KS3AT3 following SEC purification. SEC-purified DEBS KS3AT3 (5 μM) was incubated with 5 μM, 50 μM, or 250 μM of the above ACP2(2) forms (1–3) in 10 mM HEPES, 100 μM TCEP, and 50 mM, 200 mM, or 530 mM citric acid, pH 7.2 [NaOH] at 20 °C for 15 min before adding 7.5 μM DBA (reaction volume = 20 μL; reaction concentrations reflect their final concentrations after all components were added). Reactions were quenched at various time points (1.7 min, 9 min, or 10 min) by the addition of an equal volume of 2x Laemmli buffer (Bio-Rad #1610737) supplemented with 5% (*v/v*) BME and heated at 95 °C for 2 min prior to SDS-PAGE analysis.

**Inhibition of DEBS M1 by Cerulenin**

To a 15.3 μL mixture of 457 mM $NaH_2PO_4$, 6.5 mM TCEP, 13 mM $MgCl_2$, 7.8 mM ATP, 3 μM PrpE, 3 μM MatB, 5 μM SCME, and 5 μM LDD(4) was added 4.1 μL of 19 μM cerulenin treated or untreated (5)M1(2) (prepared in the same manner as above for M1TE), 15 μM (3)M2TE, and 7.3 μM coenzyme A (CoA). The reactions were initiated by the addition of 0.6 μL of a mixture containing 35.7 mM sodium propionate, 35.7 mM methylmalonic acid, and 28.6 mM NADPH to arrive at a final reaction volume of 20 μL and the following final reaction concentrations: 350 mM $NaH_2PO_4$, 5 mM TCEP, 10 mM $MgCl_2$, 6 mM ATP, 2 μM PrpE, 2 μM MatB, 4 μM SCME, 4 μM LDD(4), 4 μM cerulenin treated or untreated (5)M1(2), 3 μM (3)M2TE, 1.5 mM CoA, 1 mM sodium propionate, 1 mM methylmalonic acid, and 0.8 mM NADPH. The 20 μL reactions were transferred to 384-well clear-bottom plates (Corning, product # 3765) to measure depletion of NADPH absorbance at 340 nm every 10 s using a BioTek Synergy HT plate reader at 20 °C (Fig. S4D).

**Preparation of Fluorescein-labeled ACP2(2) (ACP2(2)-FL)**

A 50 mM stock solution of fluorescein isothiocyanate (FITC) was freshly prepared in anhydrous DMSO. To 250 μM *apo*-ACP2(2) in 0.3 M $NaH_2PO_4$ pH 8 [NaOH] was added 1, 2.5, 5, or 10 equivalents of FITC (i.e., 4 different 50 μL reactions). The reactions were terminated by removing unreacted FITC using 7 kDa MWCO Zeba spin desalting columns (Thermo Fisher: 89882) equilibrated with 0.1 M $NaH_2PO_4$ pH 7.2 [NaOH]. The eluent containing fluorescein-labeled *apo*-ACP2(2) (*apo*-ACP2(2)-**FL**) was 4′-phosphopantetheinylated by incubating 209 μM *apo*-ACP2(2)-**FL** with 20.9 μM Sfp, 3 mM CoA, and 10 mM $MgCl_2$ in 0.1 M $NaH_2PO_4$ pH 7.2 [NaOH] for 1 h at 20 °C to generate *holo*-form ACP2(2)-**FL** (referred to throughout as "ACP2(2)-**FL**"). To measure the extent of fluorescein labeling, the absorbance at 500 nm, corresponding to the experimental maximum absorbance of ACP2(2)-**FL**, was measured using a NanoDrop 2000c (Thermo Fisher Scientific) to calculate the dye-to-protein ratios for each ACP2(2)-**FL** sample. A maximum dye-to-protein ratio of 0.25 was measured (i.e., corresponding to conjugation reactions in which *apo*-ACP2(2) was incubated with 10 equivalents of FITC).

**Limited Trypsinolysis of DEBS M3TE**

We utilized a previously identified[11] trypsin-labile site at the KR-ACP junction of DEBS M3 to characterize the crosslinked species (bands **1** and **2**) by limited trypsinolysis. Trypsin was prepared at a concentration of 0.5 mg/mL in 50 mM sodium acetate pH 5 [NaOH] for long term storage at -80 °C and subsequently diluted to 50 μg/mL in 0.1 M $NaH_2PO_4$ pH 7.2 [NaOH] before usage in trypsinolysis experiments. M3TE (5 μM) and CL-M3TE (5 μM) were trypsinized at 20 °C in the presence of 5 μg/mL trypsin in 100 mM citric acid, 50 mM BME, 5 mM HEPES, 50 mM $NaH_2PO_4$, pH 7.2 [NaOH]. Trypsin was inactivated after 1, 4, 10, and 16 min by the addition of an equal volume of 2x Laemmli buffer (Bio-Rad #1610737) supplemented with 5% (v/v) BME and heated at 95 °C for 2 min prior to SDS-PAGE analysis. Taking the panel of putative crosslinked structures, we made *a priori* predications of the expected tryptic products, assuming a single cleavage event at the equivalent trypsin-labile sites in each subunit (Fig. S8). Comparison of the tryptic products in the DBA-crosslinked versus un-crosslinked M3TE revealed transient species consistent with band **1** corresponding to an asymmetric, singly crosslinked dimer (i.e., *State 1*) (Fig. S9). We considered band **2** to be a self-crosslinked monomer, in accordance with its expected mobility relative to a singly crosslinked dimer (Figs. 1B and S5). This assignment was further supported by fluorescent probe crosslinking (Figs. S6–S7) and cryo-EM analysis (Figs. S17–S19; S23).

**Isolation of Crosslinked and Un-crosslinked Module + $F_{ab}$ 1B2 Complexes**

All DEBS modules (crosslinked or un-crosslinked) and $F_{ab}$ 1B2 used in cryo-EM experiments were individually purified via SEC prior to preparation and re-purification of the module-$F_{ab}$ complexes. Module-1B2 complexes were prepared by adding 1.5 equivalents of 1B2 heterodimer per equivalent of PKS monomer, in accordance with the binding stoichiometry[12], and incubated on ice for 30 min before isolating the complex by SEC. Protein complex samples (≤ 2 mL) were injected onto a 120 mL Superdex 200 pg 16/600 column (Cytiva) at a flow rate of 1.5 mL/min and fractionated into 3 mL isocratically with SEC buffer using an ÄKTA Pure protein purification FPLC system (Cytiva) (Fig. S12A, B).

**Isolation of Crosslinked KSAT-ACP Complexes**
Protein complex samples (≤ 2 mL) were injected onto a 120 mL Superdex 200 pg 16/600 column (Cytiva) at a flow rate of 1.5 mL/min and fractionated into 3 mL isocratically with SEC buffer containing 0.3 M citric acid using an ÄKTA Pure protein purification FPLC system (Cytiva) (Fig. S12C–E).

**Cryo-EM Sample Preparation and Data Collection**
Crosslinked and un-crosslinked DEBS module-$F_{ab}$ complexes were concentrated to 5–10 mg/mL using Amicon Ultra Centrifugal Filters (50 kDa MWCO) before adding 0.03% nonyl phenoxypolyethoxylethanol (NP-40) and applying 3 μL onto glow-discharged 300-mesh R 2/1 Quantifoil copper grids. The grids were blotted for 4 s at 4 °C and 100% relative humidity and vitrified in liquid ethane using a Vitrobot Mark IV (Thermo Fisher Scientific). Grids of crosslinked KSAT-ACP complexes were prepared in the same way; except they were diluted from 0.3 M to 0.1 M citric acid prior to vitrification. Samples were imaged at 300 kV accelerating voltage with a Titan Krios G3i cryo-electron microscope (Thermo Fisher Scientific) equipped with a K3 (Gatan) or Falcon4 (Thermo Fisher Scientific) direct-electron detector (DED) and BioQuantum or Selectris energy filters at nominal magnifications of 81,000× or 130,000×, corresponding to a calibrated sampling of 1.1 Å/pixel or 0.946 Å/pixel, respectively. EPU software (Thermo Fisher Scientific) was used to record dose-fractionated movies composed of 40 individual frames in Lzw non-gain-normalized .tiff format (with Gatan K3 detector) or gain-normalized .mrc format (with Falcon 4 detector) and with a total dose of 50 electrons and dose rates of 6.9, 16.1, and 21.7 $e^{-}\cdot pixel^{-1}\cdot s^{-1}$ (see Table S2 for details).

**Single-particle Cryo-EM Image Processing and 3D Reconstruction**
Single-particle cryo-EM image analysis was carried out in Relion[13] and cryoSPARC[14], and statistical per-particle image analysis was implemented in EMAN2[15] (see Figs. S13–S14, S17–S22, S24–S25, and S26–S27 for details). In all cases, dose-fractionated image stacks were applied to motion correction, dose weighting, and contrast transfer function (CTF) estimation before reference-free automated particle picking. In some cases, initial particles were used to generate templates by 2D classification followed by template-based particle picking. The full particle-batches were applied directly to ab-initio reconstruction in cryoSPARC for CL-M1TE-1B2 and CL-M3TE-1B2 datasets, whereas iterative rounds of 2D classification were implemented prior to ab-initio reconstruction for CL-KS3AT3-ACP3 and CL-KS3AT3-ACP2(2)/ACP3 datasets. C1 symmetry was specified for all ab-initio reconstructions and in subsequent homogenous/heterogeneous

refinements. Particles were transported from Relion to cryoSPARC via “import particle stack” in cryoSPARC; whereas, particles in cryoSPARC format (.cs) were converted to .star format for transport into Relion via the csparc2star.py script in pyem[16].

**Statistical Per-Particle Image Analysis**

Crosslinked KS3AT3 + ACP3 / Crosslinked M1TE (Fig. 5): All particles were first aligned to the pseudo-C2 symmetry axis and then segmented into two, symmetrically related asymmetric units. A soft spherical mask centered at the ACP binding site was used for focused 3D classification into the two classes shown in Figure 5. C2-particles were regrouped into their original C1-particles by symmetry-mate pairing and classified as bound to 0, 1, or 2 ACPs based on the class assignments of their constituent asymmetric units.

Crosslinked KS3AT3 + ACP2(2) + ACP3 (Fig. S27): All particles were first aligned to the pseudo-C2 symmetry axis and then segmented into two, symmetrically related asymmetric units. A soft spherical mask centered at the ACP binding site was used for focused 3D classification into the four classes shown in Figure S27A. C2-particles were regrouped into their original C1-particles by symmetry-mate pairing and classified as 1 of 16 possible classes based on the class assignments of their constituent asymmetric units. Due to different particle counts in each of the four classes (Fig. S27B), we were unable to assess intersubunit coupling in the same way as with KS3AT3 + ACP3 (Fig. 5A).

#### Protein Sequences

| *Key:* |
| --- |
| KS = ketosynthase, AT = acyltransferase, KR = ketoreductase, ACP = acyl carrier protein; TE = thioesterase |
| DEBS M1 |
| DEBS M2; **DEBS M2 Docking Domain** |
| DEBS M3; **DEBS M3 Docking Domain** |
| DEBS TE (M6) |
| $F_{ab}$ 1B2 |
| Linkers/Tags |

*DEBS M3 with its native N-terminal docking domain and a C-terminal TE domain from M6 (M3TE; pRSG34)* **|** **(3)**-KS3-AT3-KR3-ACP3-TE

**MASTDSEKVAEYLRRATLDLRAARQRIRELE**SDPIAIVSMACRLPGGVNTPQRLWELLREGGET
LSGFPTDRGWDLARLHHPDPDNPGTSYVDKGGFLDDAAGFDAEFFGVSPREAAAMDPQQRLLLE
TSWELVENAGIDPHSLRGTATGVFLGVAKFGYGEDTAAAEDVEGYSVTGVAPAVASGRISYTMG
LEGPSISVDTACSSSLVALHLAVESLRKGESSMAVVGGAAVMATPGVFVDFSRQRALAADGRSK
AFGAGADGFGFSEGVTLVLLERLSEARRNGHEVLAVVRGSALNQDGASNGLSAPSGPAQRRVIR
QALESCGLEPGDVDAVEAHGTGTALGDPIEANALLDTYGRDRDADRPLWLGSVKSNIGHTQAAA
GVTGLLKVVLALRNGELPATLHVEEPTPHVDWSSGGVALLAGNQPWRRGERTRRARVSAFGISG
TNAHVIVEEAPEREHRETTAHDGRPVPLVVSARTTAALRAQAAQIAELLERPDADLAGVGLGLA
TTRARHEHRAAVVASTREEAVRGLREIAAGAATADAVVEGVTEVDGRNVVFLFPGQGSQWAGMG
AELLSSSPVFAGKIRACDESMAPMQDWKVSDVLRQAPGAPGLDRVDVVQPVLFAVMVSLAELWR
SYGVEPAAVVGHSQGEIAAAHVAGALTLEDAAKLVVGRSRLMRSLSGEGGMAAVALGEAAVRER
LRPWQDRLSVAAVNGPRSVVVSGEPGALRAFSEDCAAEGIRVRDIDVDYASHSPQIERVREELL
ETTGDIAPRPARVTFHSTVESRSMDGTELDARYWYRNLRETVRFADAVTRLAESGYDAFIEVSP
HPVVVQAVEEAVEEADGAEDAVVVGSLHRDGGDLSAFLRSMATAHVSGVDIRWDVALPGAAPFA
LPTYPFQRKRYWLQPAAPAAASDELAYRVSWTPIEKPESGNLDGDWLVVTPLISPEWTEMLCEA
INANGGRALRCEVDTSASRTEMAQAVAQAGTGFRGVLSLLSSDESACRPGVPAGAVGLLTLVQA
LGDAGVDAPVWCLTQGAVRTPADDDLARPAQTTAHGFAQVAGLELPGRWGGVVDLPESVDDAAL
RLLVAVLRGGGRAEDHLAVRDGRLHGRRVVRASLPQSGSRSWTPHGTVLVTGAASPVGDQLVRW
LADRGAERLVLAGACPGDDLLAAVEEAGASAVVCAQDAAALREALGDEPVTALVHAGTLTNFGS
ISEVAPEEFAETIAAKTALLAVLDEVLGDRAVEREVYCSSVAGIWGGAGMAAYAAGSAYLDALA
EHHRARGRSCTSVAWTPWALPGGAVDDGYLRERGLRSLSADRAMRTWERVLAAGPVSVAVADVD
WPVLSEGFAATRPTALFAELAGRGGQAEAEPDSGPTGEPAQRLAGLSPDEQQENLLELVANAVA
EVLGHESAAEINVRRAFSELGLDSLNAMALRKRLSASTGLRLPASLVFDHPTVTALAQHTSQLD
SGTPAREASSALRDGYRQAGVSGRVRSYLDLLAGLSDFREHFDGSDGFSLDLVDMADGPGEVTV
ICCAGTAAISGPHEFTRLAGALRGIAPVRAVPQPGYEEGEPLPSSMAAVAAVQADAVIRTQGDK
PFVVAGHSAGALMAYALATELLDRGHPPRGVVLIDVYPPGHQDAMNAWLEELTATLFDRETVRM
DDTRLTALGAYDRLTGQWRPRETGLPTLLVSAGEPMGPWPDDSWKPTWPFEHDTVAVPGDHFTM
VQEHADAIARHIDAWLGGGNSSSVDKLAAALEHHHHHH

*DEBS M1 with an N-terminal docking domain from M3 and a C-terminal TE domain from M6 (M1TE; pDC1)*[10] **|** **(3)**-KS1-AT1-KR1-ACP1-TE

**MASTDSEKVAEYLRRATLDLRAARQRIRELE**GEPVAVVAMACRLPGGVSTPEEFWELLSEGRDA
VAGLPTDRGWDLDSLFHPDPTRSGTAHQRGGGFLTEATAFDPAFFGMSPREALAVDPQQRLMLE
LSWEVLERAGIPPTSLQASPTGVFVGLIPQEYGPRLAEGGEGVEGYLMTGTTTSVASGRIAYTL
GLEGPAISVDTACSSSLVAVHLACQSLRRGESSLAMAGGVTVMPTPGMLVDFSRMNSLAPDGRC
KAFSAGANGFGMAEGAGMLLLERLSDARRNGHPVLAVLRGTAVNSDGASNGLSAPNGRAQVRVI
QQALAESGLGPADIDAVEAHGTGTRLGDPIEARALFEAYGRDREQPLHLGSVKSNLGHTQAAAG
VAGVIKMVLAMRAGTLPRTLHASERSKEIDWSSGAISLLDEPEPWPAGARPRRAGVSSFGISGT
NAHAIIEEAPQVVEGERVEAGDVVAPWVLSASSAEGLRAQAARLAAHLREHPGQDPRDIAYSLA
TGRAALPHRAAFAPVDESAALRVLDGLATGNADGAAVGTSRAQQRAVFVFPGQGWQWAGMAVDL
LDTSPVFAAALRECADALEPHLDFEVIPFLRAEAARREQDAALSTERVDVVQPVMFAVMVSLAS
MWRAHGVEPAAVIGHSQGEIAAACVAGALSLDDAARVVALRSRVIATMPGNKGMASIAAPAGEV
RARIGDRVEIAAVNGPRSVVVAGDSDELDRLVASCTTECIRAKRLAVDYASHSSHVETIRDALH
AELGEDFHPLPGFVPFFSTVTGRWTQPDELDAGYWYRNLRRTVRFADAVRALAEQGYRTFLEVS
AHPILTAAIEEIGDGSGADLSAIHSLRRGDGSLADFGEALSRAFAAGVAVDWESVHLGTGARRV
PLPTYPFQRERVWLEPKPVARRSTEVDEVSALRYRIEWRPTGAGEPARLDGTWLVAKYAGTADE
TSTAAREALESAGARVRELVVDARCGRDELAERLRSVGEVAGVLSLLAVDEAEPEEAPLALASL
ADTLSLVQAMVSAELGCPLWTVTESAVATGPFERVRNAAHGALWGVGRVIALENPAVWGGLVDV
PAGSVAELARHLAAVVSGGAGEDQLALRADGVYGRRWVRAAAPATDDEWKPTGTVLVTGGTGGV
GGQIARWLARRGAPHLLLVSRSGPDADGAGELVAELEALGARTTVAACDVTDRESVRELLGGIG
DDVPLSAVFHAAATLDDGTVDTLTGERIERASRAKVLGARNLHELTRELDLTAFVLFSSFASAF
GAPGLGGYAPGNAYLDGLAQQRRSDGLPATAVAWGTWAGSGMAEGPVADRFRRHGVIEMPPETA
CRALQNALDRAEVCPIVIDVRWDRFLLAYTAQRPTRLFDEIDDARRAAPQAAAEPRVGALASLP
APEREKALFELVRSHAAAVLGHASAERVPADQAFAELGVDSLSALELRNRLGAATGVRLPTTTV
FDHPDVRTLAAHLTSELGSGTPAREASSALRDGYRQAGVSGRVRSYLDLLAGLSDFREHFDGSD
GFSLDLVDMADGPGEVTVICCAGTAAISGPHEFTRLAGALRGIAPVRAVPQPGYEEGEPLPSSM
AAVAAVQADAVIRTQGDKPFVVAGHSAGALMAYALATELLDRGHPPRGVVLIDVYPPGHQDAMN
AWLEELTATLFDRETVRMDDTRLTALGAYDRLTGQWRPRETGLPTLLVSAGEPMGPWPDDSWKP
TWPFEHDTVAVPGDHFTMVQEHADAIARHIDAWLGGGNSSSVDKLAAALEHHHHHH

*KS3AT3 (pAYC02)*

**MVTDSEKVAEYLRRATLDLRAARQRIRELE**SDPIAIVSMACRLPGGVNTPQRLWELLREGGETL
SGFPTDRGWDLARLHHPDPDNPGTSYVDKGGFLDDAAGFDAEFFGVSPREAAAMDPQQRLLLET
SWELVENAGIDPHSLRGTATGVFLGVAKFGYGEDTAAAEDVEGYSVTGVAPAVASGRISYTMGL
EGPSISVDTACSSSLVALHLAVESLRKGESSMAVVGGAAVMATPGVFVDFSRQRALAADGRSKA
FGAGADGFGFSEGVTLVLLERLSEARRNGHEVLAVVRGSALNQDGASNGLSAPSGPAQRRVIRQ
ALESCGLEPGDVDAVEAHGTGTALGDPIEANALLDTYGRDRDADRPLWLGSVKSNIGHTQAAAG
VTGLLKVVLALRNGELPATLHVEEPTPHVDWSSGGVALLAGNQPWRRGERTRRAAVSAFGISGT
NAHVIVEEAPEREHRETTAHDGRPVPLVVSARSTAALRAQAAQIAELLERPDADLAGVGLGLAT
TRARHEHRAAVVASTREEAVRGLREIAAGAATADAVVEGVTEVDGRNVVFLFPGQGSQWAGMGA
ELLSSSPVFAGKIRACDESMAPMQDWKVSDVLRQAPGAPGLDRVDVVQPVLFAVMVSLAELWRS
YGVEPAAVVGHSQGEIAAAHVAGALTLEDAAKLVVGRSRLMRSLSGEGGMAAVALGEAAVRERL
RPWQDRLSVAAVNGPRSVVVSGEPGALRAFSEDCAAEGIRVRDIDVDYASHSPQIERVREELLE
TTGDIAPRPARVTFHSTVESRSMDGTELDARYWYRNLRETVRFADAVTRLAESGYDAFIEVSPH
PVVVQAVEEAVEEADGAEDAVVVGSLHRDGGDLSAFLRSMATAHVSGVDIRWDVALPGAAPFAL
PTYPFQRKRYWLQPAAPAAASDELAYRSSSVDKLAAALEHHHHHH

*ACP2(2) (pNW6)*

MGSSHHHHHHSSGLVPRGSHMLRDRLAGLPRAERTAELVRLVRTSTATVLGHDDPKAVRATTPF
KELGFDSLAAVRLRNLLNAATGLRLPSTLVFDHPNASAVAGFLDAELG**TEVRGEAPSALAGLDA
LEAALPEVPATEREELVQRLERMLAALRPVAQAADASGTGANPSGDDLGEAGVDELLEALGREL
DGD**

*ACP2 (pNW7)*

MGSSHHHHHHSSGLVPRGSHMLRDRLAGLPRAERTAELVRLVRTSTATVLGHDDPKAVRATTPF
KELGFDSLAAVRLRNLLNAATGLRLPSTLVFDHPNASAVAGFLDAELG

*ACP3 (pDC9)*

MGSSHHHHHHSSGLVPRGSHGLSPDEQQENLLELVANAVAEVLGHESAAEINVRRAFSELGLDS
LNAMALRKRLSASTGLRLPASLVFDHPTVTALAQHLRARLVK

*$F_{ab}$ 1B2 (heavy chain)*

MAEVQLVQSGGGLVQPGRSLRLSCTASGFTFGDYAMSWVRQAPGKGLEWVGFIRSKAYGGTTEY
AASVKGRFTISRDDSKSIAYLQMNSLKTEDTAVYYCTRGGTLFDYWGQGTLVTVSSASTKGPSV
FPLAPSSKSTSGGTAALGCLVKDYFPEPVTVSWNSGALTSGVHTFPAVLQSSGLYSLSSVVTVP
SSSLGTQTYICNVNHKPSNTKVDKKVEPKSCAALVPRGSAHHHHHHAADYKDDDDKA

*$F_{ab}$ 1B2 (light chain)*

LFAIPLVVPFYSHSALDVVMTQSPLSLPVTPGEPASISCRSSQSLLHSNGYNYLDWYLQKPGQS
PQLLIYLGSNRASGVPDRFSGSGSGTDFTLKISRVEAEDVGVYYCMQSLQTPRLTFGPGTKVDI
KRTVAAPSVFIFPPSDEQLKSGTASVVCLLNNFYPRGAKVQWKVDNALQSGNSQESVTEQDSKD
STYSLSSTLTLSKADYEKHKVYACEVTHQGLSSPVTKSFNRGEC

#### Supplementary Figures

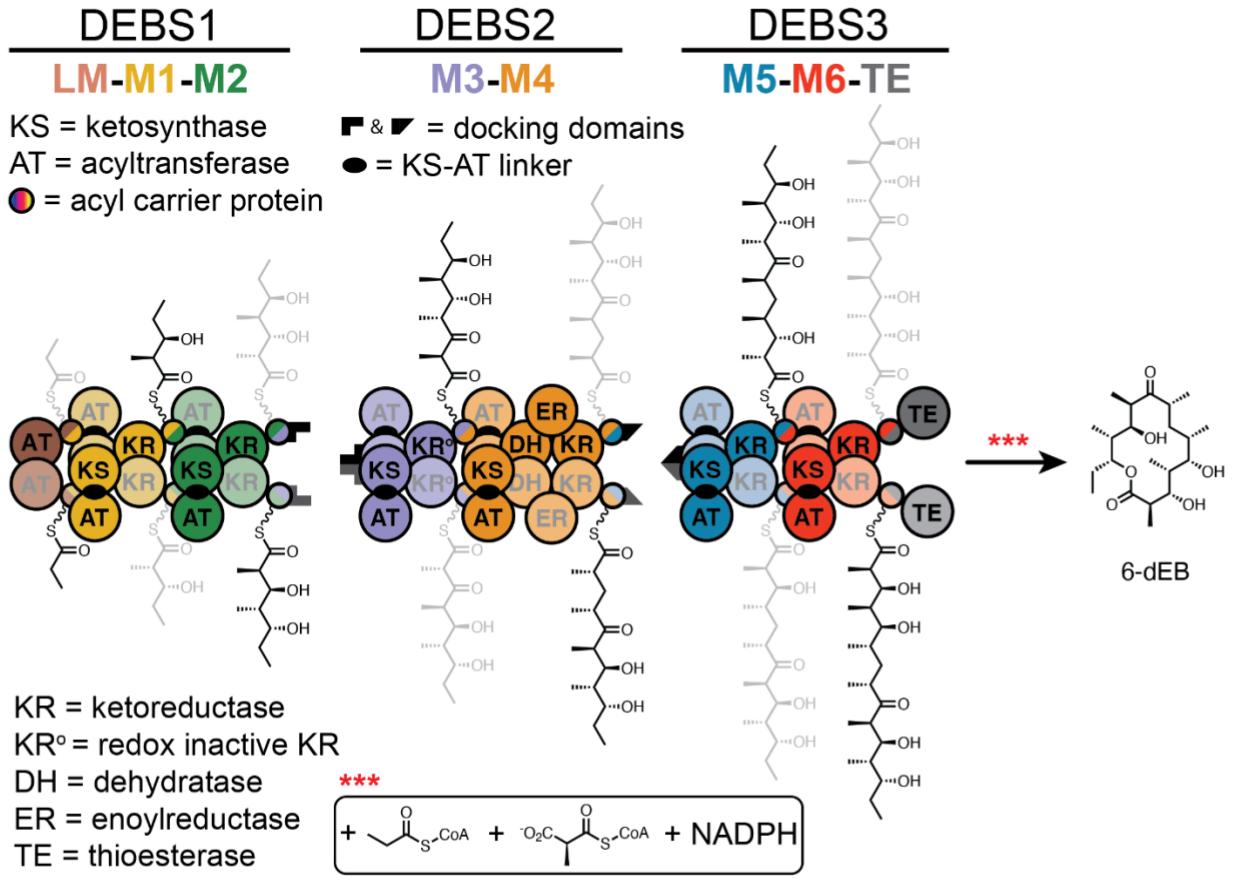

**Fig. S1.** The ~2 MDa 6-deoxyerythronolide B (6-dEB) synthase (DEBS). DEBS consists of three homodimeric polypeptides, DEBS 1–3. The six elongating modules (M1–M6) are flanked by an N-terminal loading module (LM) and a C-terminal thioesterase (TE) domain and collectively assemble 6-dEB from propionyl-CoA, (2*S*)-methylmalonyl-CoA, and NADPH precursors (red asterisks). Each subunit of homodimeric DEBS 1–3 is distinguished by heavy and light shading. The growing polyketide intermediate is similarly distinguished by heavy and light shading to illustrate its directional channeling via alternating subunit attachment points during biosynthesis. This is because polyketide elongation occurs via an *inter*-subunit KS-ACP interaction, whereas translocation occurs via an *intra*-subunit KS-ACP interaction (Fig. S2)[10,17]. The C-terminal TE domain catalyzes macrocyclization of the fully elongated intermediate to offload 6-dEB from the assembly line.

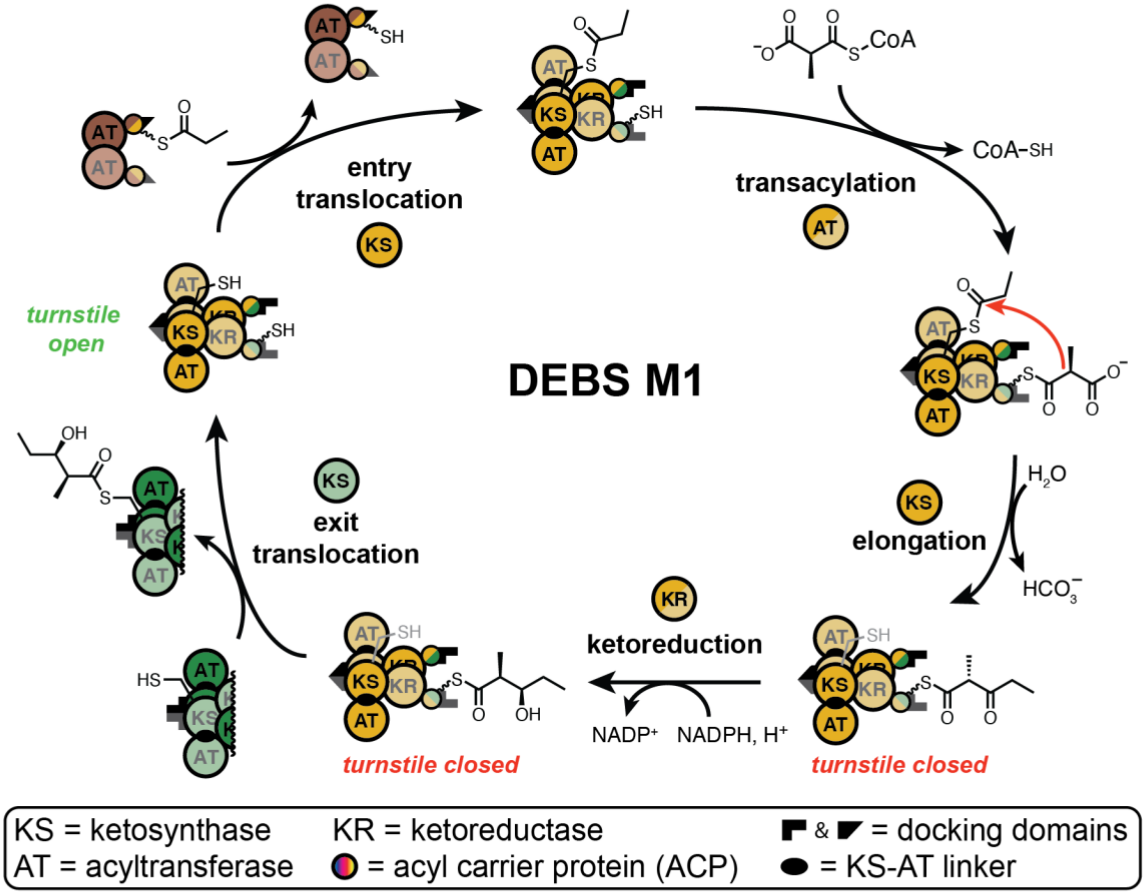

**Fig. S2.** The catalytic cycle of DEBS Module 1 (M1), a representative assembly-line PKS module (Fig. S1). Both subunits are shown, distinguished by heavy and light shading, to depict both inter- and intra-molecular events during the catalytic cycle (i.e., inter-molecular elongation and intra-molecular translocation)[10,17]. Only one polyketide intermediate is depicted as being processed, consistent with our previous finding that a homodimeric module is ~50% occupied under steady-state turnover conditions[2]. For clarity, the modules in this scheme are represented as stand-alone entities that interact *in trans* via shape complementary N- and C-terminal docking domains[4,18]. Although natively LM, M1, and M2 interact as part of a single, homodimeric polypeptide (Fig. S1), they can be reconstituted as stand-alone proteins in the way that is depicted (Fig. S4E; see also Ref. 1). The catalytic cycle of M1 begins with KS-catalyzed transfer of a propionyl group onto the KS active site Cys residue (entry translocation). Meanwhile, the AT domain selects an appropriate acyl-CoA extender unit for transfer of a (2*S*)-methylmalonyl group onto the 4′-phosphopantetheine (Ppant) thiol cofactor of an ACP (transacylation). The KS then catalyzes decarboxylative Claisen condensation between the KS-bound electrophile and ACP-bound nucleophile to form an elongated (2*R*)-methyl-3-ketopentanoyl-ACP thioester product (elongation). There is now evidence that elongation is energetically coupled to a "turnstile" mechanism for unidirectional biosynthesis along the assembly line. Biochemical[2] and structural[10] data support a mechanism in which elongation induces a conformational change in the module's AT domains such that acyl-ACP substrates or products can no longer enter their KS active sites (denoted as *turnstile closed*). Such a mechanism ensures that assembly-line PKS modules do not behave iteratively and that only one intermediate is processed by the module at a given time, for increased fidelity. Other domains beyond the core KS, AT, and ACP domains may then modify the elongated product by setting its redox state, substitution pattern, and/or stereochemistry at the newly formed α- and β-positions[19]. In the case of DEBS M1, a KR domain stereospecifically reduces the β-ketone while epimerizing the α-stereocenter to form a (2*S*,3*R*)-3-hydroxy-2-methylpentanoyl-ACP thioester product (ketoreduction). The modified and elongated product is then shuttled to the KS domain of DEBS Module 2 (exit translocation) which is catalyzed by the recipient KS domain through a process analogous to entry translocation. This reaction is concomitant with turnstile opening, thereby reactivating the KS Cys residue to begin the next catalytic cycle.

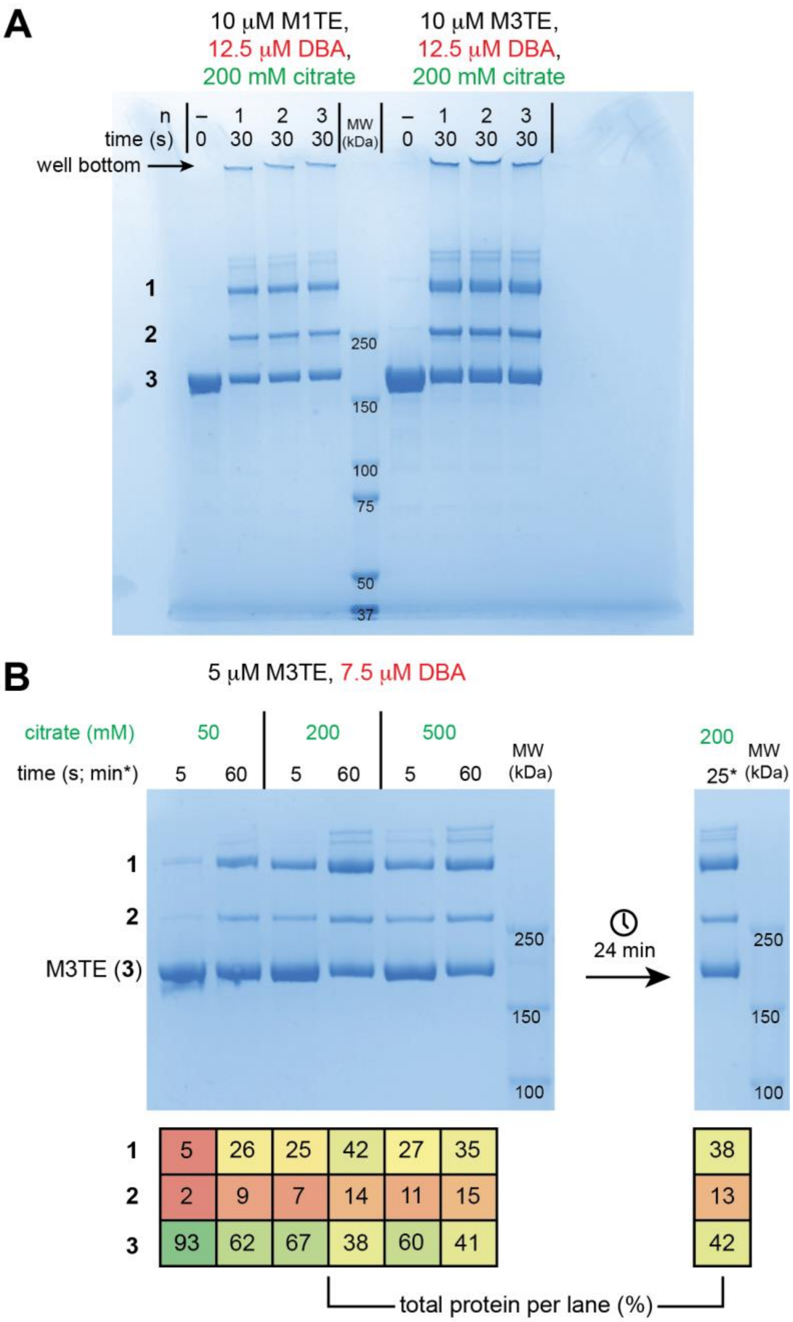

**Fig. S3.** (**A**) Uncropped SDS-PAGE analysis of DBA-mediated crosslinking of DEBS M1TE and M3TE (cropped for use in Fig. 1B). Three replicate 30 s reactions (n) were analyzed alongside the unreacted M1TE or M3TE (–). Bands **1** and **2** correspond to DBA-dependent crosslinked products, and band **3** corresponds to the un-crosslinked M1TE/M3TE monomers. (**B**) Time- and citrate-dependent DBA-mediated crosslinking of M3TE. Three different reactions were quenched at 2 or 3 different time points (i.e., 5 s, 60 s, and 25 min) by the addition of 2.5% (v/v) BME and carried out under three different citrate concentrations (i.e., 50 mM, 200 mM, and 500 mM). The reactions reached ~50% completion by 5 s. Each band was quantified in GelAnalyzer 19.1 and tabulated as its percentage of the total protein per lane. Residual protein amounts (<10%) per lane are accounted for by the high molecular weight products (MW > band **1**). For gel source data, see Supplementary Raw Data.

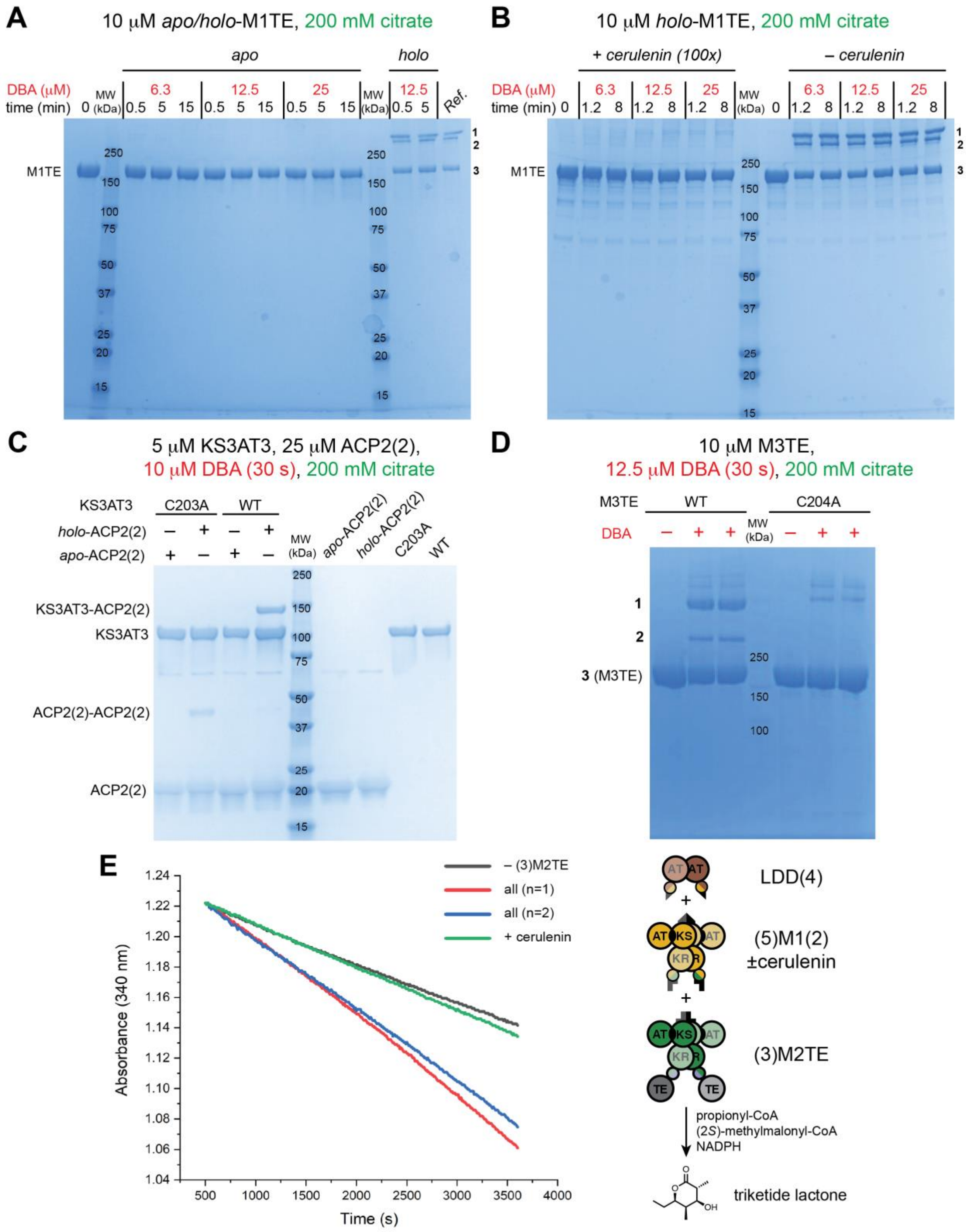

**Fig. S4.** DBA crosslinking of DEBS M1TE and the KS-AT didomain fragment of Module 3 (KS3AT3) depends on both KS and ACP thiols. (**A**) Bands **1** and **2** do not form when the Ppant thiol is removed from M1TE (i.e., when prepared in the *apo*-form) or (**B**) when the KS active site Cys is inactivated by the epoxide inhibitor, cerulenin. (**C**) Crosslinking between KS3AT3 and ACP2(2) also does not occur when the ACP Ppant thiol is removed or the KS active Cys is mutated to an Ala residue (i.e., variant C203A). Complete loss of crosslinking observed in the KS3AT3-C203A variant suggested that other nucleophilic groups in the KS active site (i.e., His, Ser, or Lys side chains) do not participate in the DBA crosslinking reaction with ACP2(2). (**D**) However, when the same Cys to Ala mutation is introduced in M3TE (i.e., M3TE-C204A), a small degree of crosslinked product was observed that appeared coincident with band **1**, indicating that another nucleophile in the KS active site may participate in crosslinking with the intramodular ACP. This nucleophile is likely to be His338 or His378 based on their proximity to Cys204 and the ability of His to react with DBA under similar conditions[20]. Note, in panels **A** and **B**, the protein samples were analyzed by SDS-PAGE using a 4–20% TGX polyacrylamide gel (Bio-Rad #4561096) which resolved bands **1** and **2** more weakly relative to the 3–8% TA polyacrylamide gels used elsewhere (e.g., panel **D**; Figs. 1B and S3A). For gel source data, see Supplementary Raw Data. (**E**) Biosynthesis of a triketide lactone by DEBS LDD(4) + (5)M1(2) + (3)M2TE was reconstituted and monitored indirectly by NADPH depletion. When the terminal (3)M2TE module was omitted ("–(3)M2TE"; black trace) or (5)M1(2) was preincubated with cerulenin ("+cerulenin"; green trace), the catalytic turnover of the bimodular assembly line was compromised, presumably due to prevention of offloading by the TE domain in (3)M2TE or the covalently inactivated (5)M1(2) by cerulenin, respectively. Traces in red or blue ("all") indicate two replicate reactions featuring every assembly line component and substrate minus cerulenin.

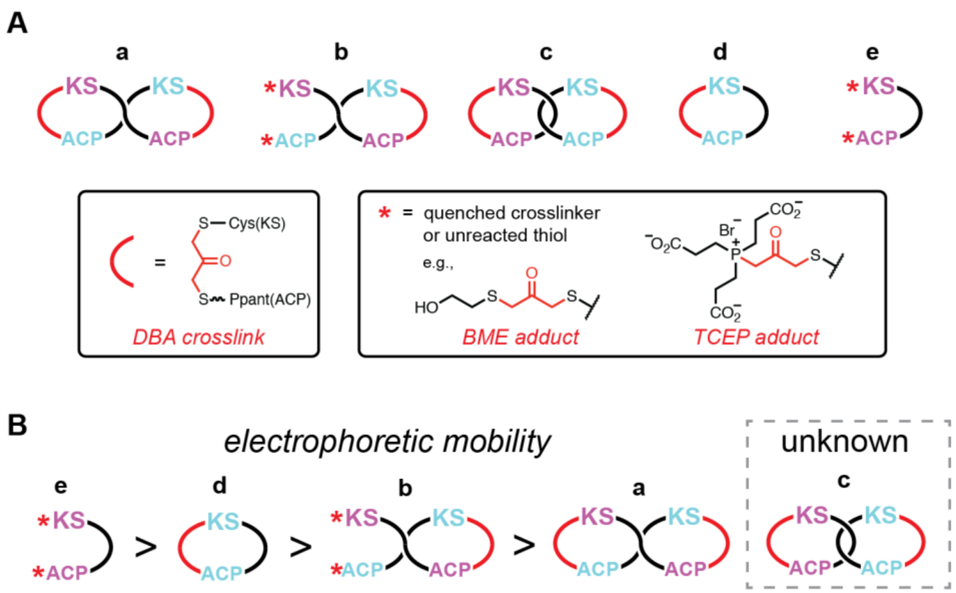

**Fig. S5.** (**A**) Putative structures of DBA-crosslinked DEBS modules. The list of possible structures was guided by a previous analysis of DBA-mediated site-specific (KS-ACP) crosslinking of the mammalian fatty acid synthase (mFAS)[21]. (**B**) Therein, Witkowski et al. discovered that self-crosslinked mFAS monomers (**d**) traveled with a decreased electrophoretic mobility relative to un-crosslinked monomers (**e**) and that inter-molecularly crosslinked mFAS dimers (**a** & **b**) traveled with reduced mobility relative to self-crosslinked monomers (**d**). Furthermore, inter-molecularly crosslinked mFAS dimers in which both KS-ACP pairs were linked (**a**) traveled with reduced mobility relative to those in which only a single KS-ACP pair was linked (**b**). Because species **c** was not detected, its relative mobility to species **a**, **b**, **d**, and **e** is unknown. For gel source data, see Supplementary Raw Data.

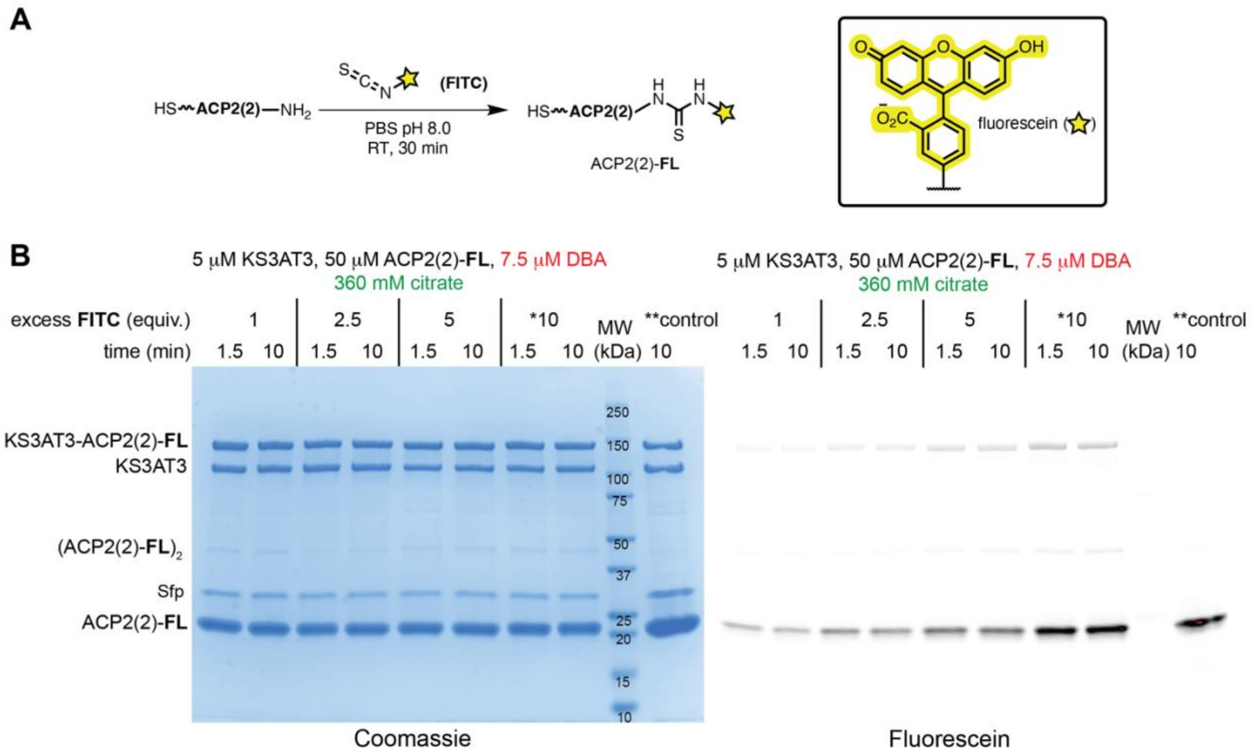

| Lane | 1 | 2 | 3 | 4 | 5 | 6 | 7 | 8 | 9 | 10 |
|---|---|---|---|---|---|---|---|---|---|---|
| KS3AT3-ACP2(2)-**FL** | 56 | 55 | 54 | 54 | 59 | 59 | 58 | 56 | – | 50 |
| KS3AT3 | 44 | 45 | 46 | 46 | 41 | 41 | 42 | 44 | – | 50 |

total protein per lane (%)

**Fig. S6.** DBA crosslinking of a fluorescent probe, ACP2(2)-**FL**, with the KS-AT fragment of DEBS Module 3 (KS3AT3). (**A**) Preparation of the fluorescent probe, ACP2(2)-**FL**. (**B**) Four different preparations of ACP2(2)-**FL** were added to KS3AT3 in 10-fold excess (see **Preparation of Fluorescein-labeled ACP2(2)**). *Prior to this experiment, ACP2(2) was incubated with 1–10 equivalents of FITC before isolating the conjugated ACP2(2)-**FL** product with a calculated maximum labeling efficiency of ~25%. **A control crosslinking reaction was carried out in which ACP2(2) lacking the fluorescein label was crosslinked with KS3AT3 and spiked with ACP2(2)-**FL** after the reaction was quenched with BME. Proteins were analyzed by SDS-PAGE using a 3–8% TA polyacrylamide gel (Invitrogen #EA0375PK2). The gel was imaged by Coomassie blue stain or by fluorescein fluorescence using a ChemiDoc MP gel imager (Bio-Rad #12003154). Adducts of KS3AT3 with ACP2(2)-FL (KS3AT3-ACP2(2)-FL) compared to un-crosslinked KS3AT3 were quantified by GelAnalyzer 19.1 and their amounts tabulated as the percentage of total protein per lane. For gel source data, see Supplementary Raw Data.

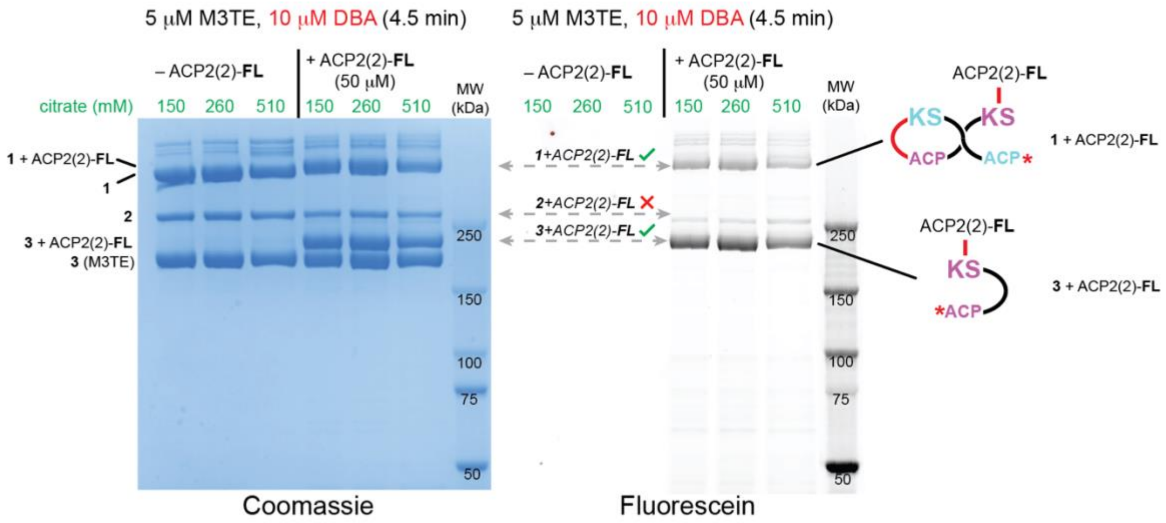

**Fig. S7.** M3TE was treated with (+ACP2(2)-**FL**; 10 equivalents) or without (–ACP2(2)-**FL**; buffer control) the fluorescent probe ACP2(2)-**FL** prior to the addition of DBA (2 equivalents) under three different citrate concentrations (150 mM, 260 mM, and 510 mM) to identify vacant KS active sites in bands **1**–**3**. Coomassie (left) and in-gel fluorescence (right) revealed the formation of two new adducts consistent with band **1** + ACP2(2)-**FL** and band **3** + ACP2(2)-**FL**. However, no major species was detected that would indicate ACP2(2)-**FL** labeling of band **2**, consistent with its assignment as a self-linked monomer (Fig. 1B). Several ACP2(2)-**FL** labeled minor products were also observed, possibly through non-specific crosslinking reactions between the probe and other Cys or His residues of M3TE (Fig. S4D). The red arcs and lines denote DBA crosslinks, and the red asterisks denote an unreacted thiol or quenched DBA group (Fig. S5A). For gel source data, see Supplementary Raw Data.

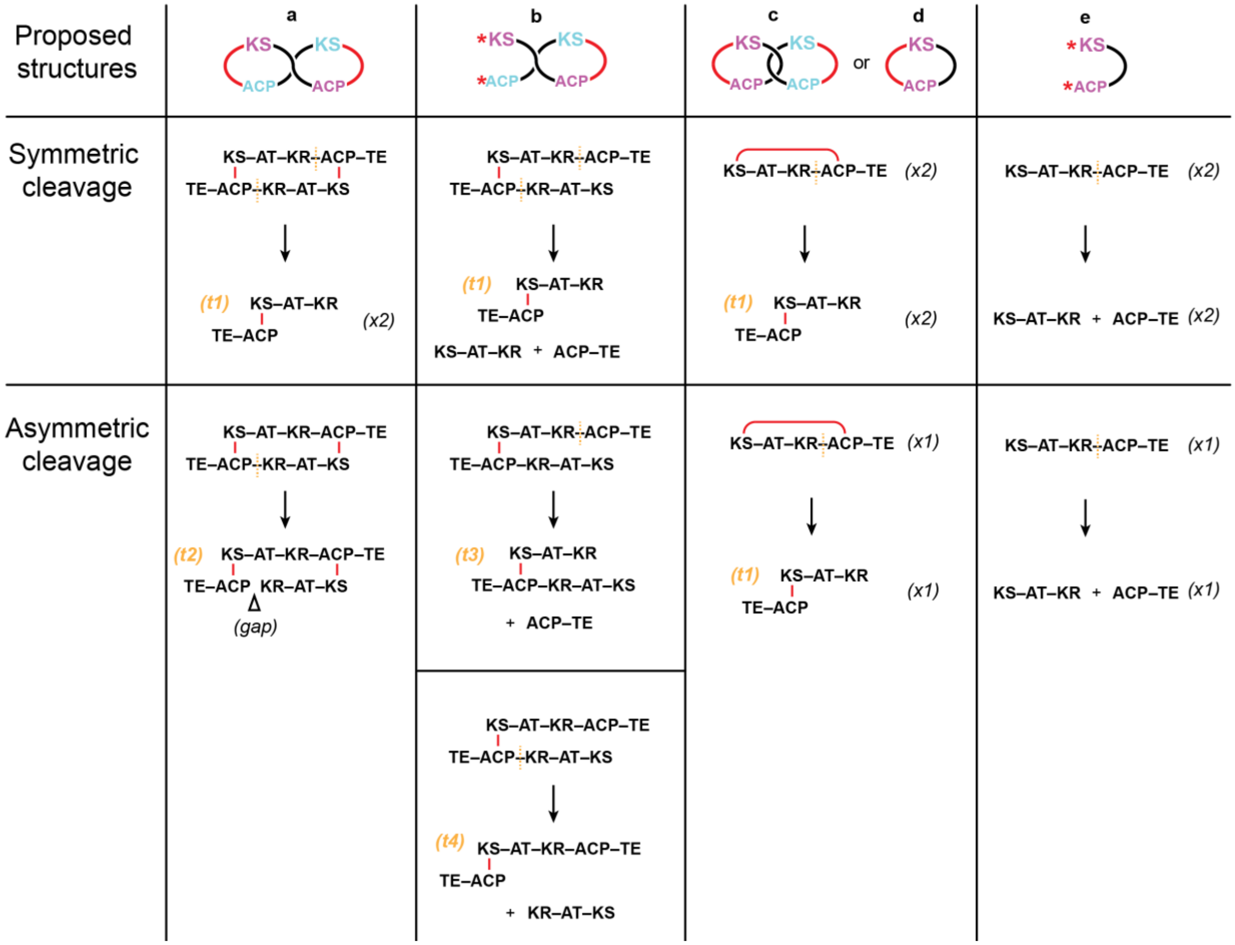

**Fig. S8.** *A priori* predictions of the tryptic products (Fig. S5A) of M3TE based on the previously identified trypsin-labile site between the KR and ACP[11]. Yellow dashed lines depict the cleavage sites, and two different scenarios were considered to account for relative differences in tryptic susceptibility of the homodimer (i.e., symmetric cleavage that implicates both tryptic sites and asymmetric cleavage that implicates only one of two sites). We envisioned that if the trypsinolysis reactions were quenched rapidly enough, transient tryptic products (***t1–t4***) might be detected via SDS-PAGE and differentiated according to their unique masses and crosslinking patterns (Fig. S9). The red arcs and lines denote DBA crosslinks, and the red asterisks denote an unreacted thiol or quenched DBA group (Fig. S5A).

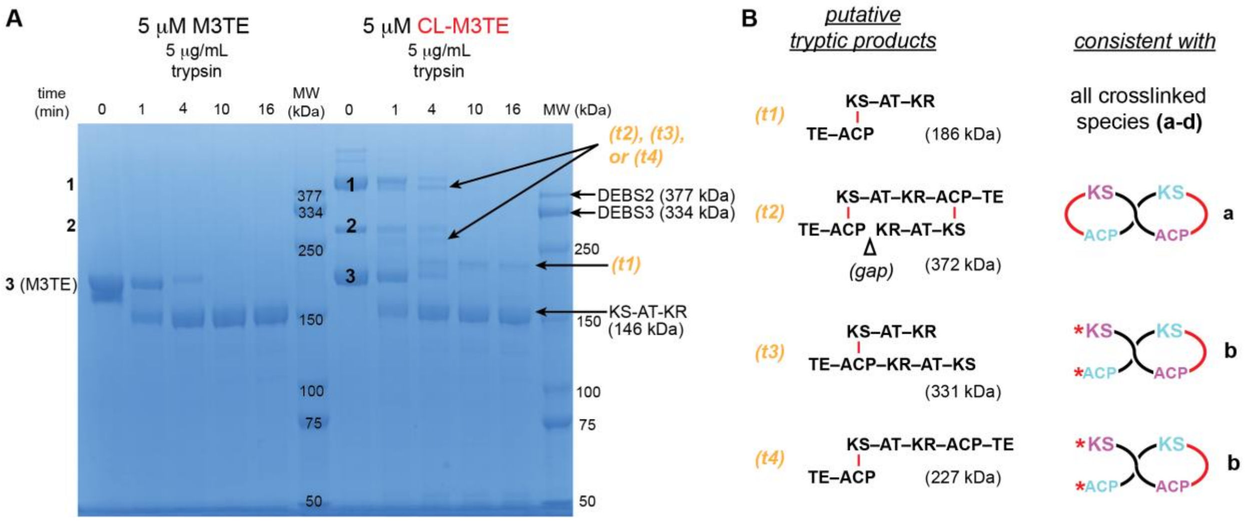

**Fig. S9.** Limited trypsinolysis of DEBS M3TE and DBA-crosslinked M3TE. Transient species observed in CL-M3TE, but not M3TE, were putatively assigned as ***t1***–***t4*** (Fig. S8). Species ***t1*** is consistent with parent species **a**-**d**, species ***t2*** is consistent with parent species **a**, and species ***t3*** and ***t4*** are consistent with parent species **b** (Fig. S5A). The fact that two transient species were observed that precede maximum accumulation of putative ***t1*** argues for band **1** corresponding to species **b** based on its unique degradation pathway relative to species **a** (Fig. S8). Recombinant, purified DEBS2 (377 kDa) and DEBS3 (334 kDa) bimodules were added to the molecular weight markers for additional mass references[1]. However, the crosslinked proteins are not expected to migrate as a function of mass in the same way as un-crosslinked protein standards[21]. For example, the M3TE dimer is ~372 kDa, and crosslinking alters its migration relative to a denatured, linear polypeptide of approximately the same mass (i.e., DEBS2). Proteins were analyzed by SDS-PAGE using a 3–8% TA polyacrylamide gel (Invitrogen #EA0375PK2). The red arcs and lines denote DBA crosslinks, and the red asterisks denote an unreacted thiol or quenched DBA group (Fig. S5A). For gel source data, see Supplementary Raw Data.

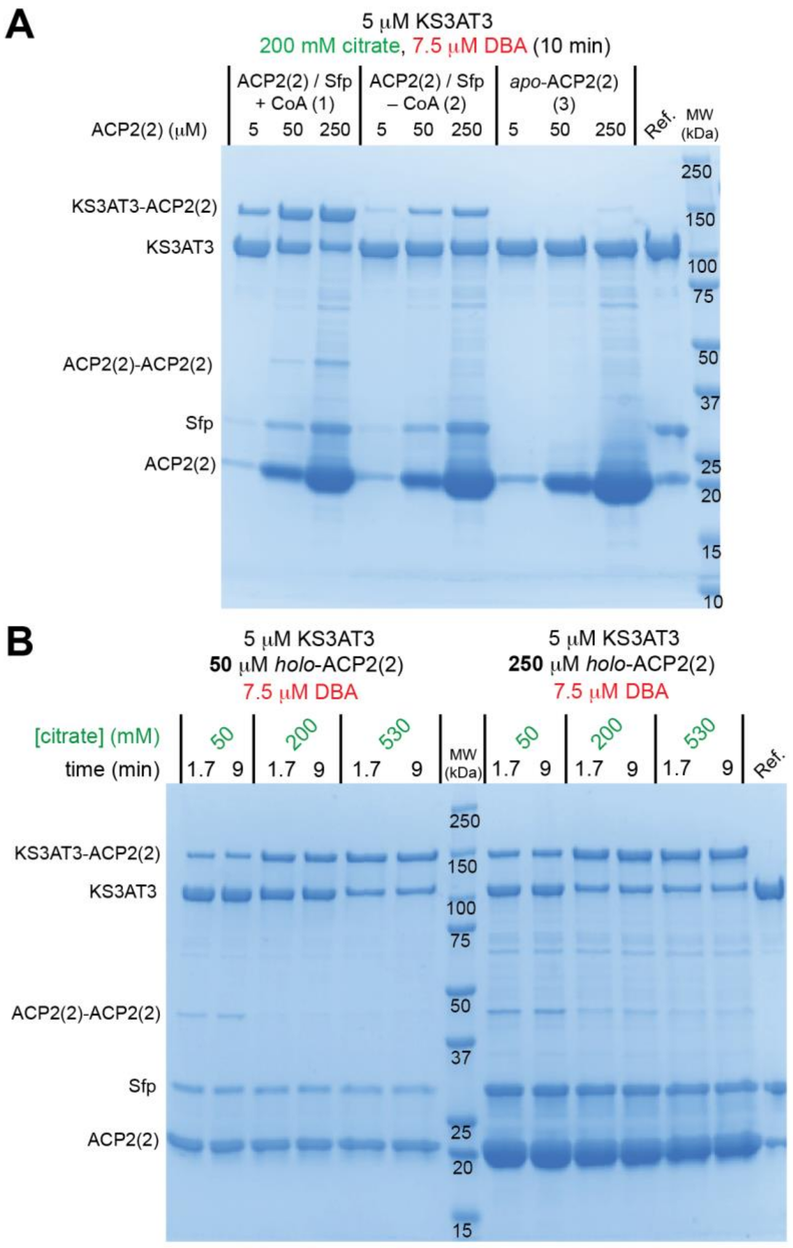

| Lane | 1 | 2 | 3 | 4 | 5 | 6 | 7 | 8 | 9 | 10 | 11 | 12 | 13 |
|---|---|---|---|---|---|---|---|---|---|---|---|---|---|
| KS3AT3-ACP2(2) | 23 | 23 | 44 | 43 | 60 | 59 | – | 38 | 37 | 62 | 62 | 65 | 67 |
| KS3AT3 | 77 | 77 | 56 | 57 | 40 | 41 | – | 62 | 63 | 38 | 38 | 35 | 33 |

total protein per lane (%)

**Fig. S10.** DBA-mediated crosslinking between the KS-AT core of DEBS Module 3 (KS3AT3) and its translocation ACP partner (ACP2(2)). (**A**) KS3AT3 was crosslinked with 1–50 equivalents of three different preparations of ACP2(2): (1) "ACP2(2) / Sfp + CoA" corresponded to *holo*-ACP2(2) derived by incubating the *apo*-protein with Sfp PPTase[5] and CoA; (2) "ACP2(2) / Sfp – CoA" corresponded to ACP2(2) incubated with Sfp only (without CoA); and (3) "*apo*-ACP2(2)" was treated with neither Sfp nor CoA (see **DBA Crosslinking Reactions**). Crosslinking was initiated by the addition of 1.5 equivalents of

DBA in the presence of 200 mM citrate. The presence of small quantities of the crosslinked adduct in preparation (2) compared to trace amounts in preparation (3) implied that recombinant Sfp co-purified with CoA in *E. coli* BL21(DE3). Endogenous PPTases in *E. coli*, AcpS[22] and AcpT[23], that converted apo-ACP2(2) into its *holo*-form might explain the trace quantities of crosslinked adduct observed in preparation (3). (**B**) DBA crosslinking of KS3AT3 with 1–50 equivalents of *holo*-ACP2(2) (preparation (1)) and 50–530 mM citrate. “Ref.” lanes include all proteins –DBA. Adducts of KS3AT3 with ACP2(2) (KS3AT3-ACP2(2)) compared to un-crosslinked KS3AT3 were quantified by GelAnalyzer 19.1 and their amounts tabulated as the percentage of total protein per lane. For gel source data, see Supplementary Raw Data.

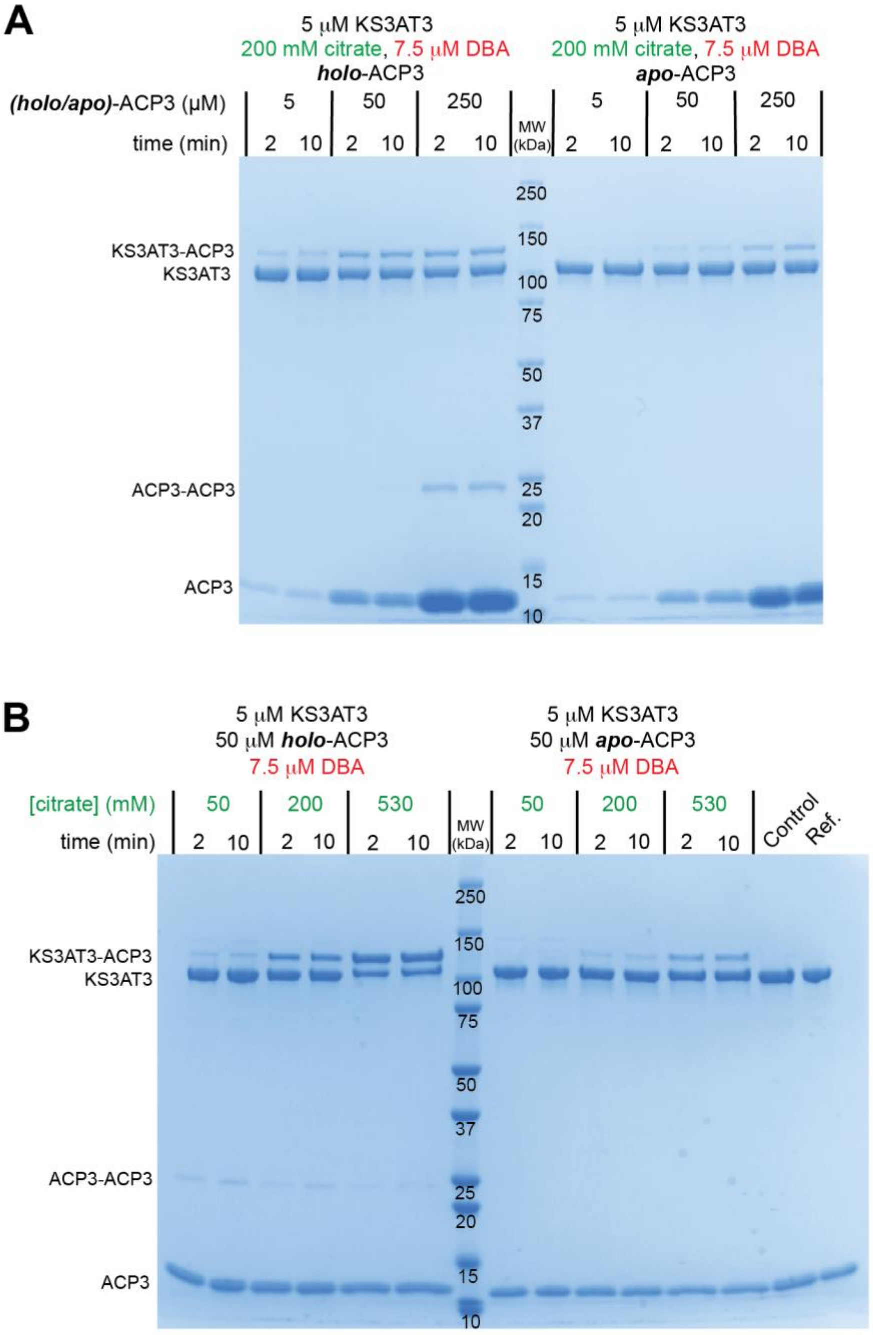

| Lane | 1 | 2 | 3 | 4 | 5 | 6 | 7 | 8 | 9 | 10 | 11 | 12 | 13 |
|---|---|---|---|---|---|---|---|---|---|---|---|---|---|
| KS3AT3-ACP3 | 4* | 5* | 31 | 32 | 60 | 58 | – | 0* | 0* | 5 | 6 | 22 | 22 |
| KS3AT3 | 95 | 93 | 69 | 68 | 40 | 42 | – | 98 | 98 | 95 | 94 | 78 | 78 |

total protein per lane (%)

**Fig. S11.** DBA crosslinking between the KS-AT core of DEBS Module 3 (KS3AT3) and ACP3. (**A**) KS3AT3 was crosslinked with 1–50 equivalents of (left) *holo*-ACP3 or (right) *apo*-ACP3 by adding 1.5 equivalents of DBA in the presence of 200 mM citrate. (**B**) Similar to panel **A**, KS3AT3 was crosslinked with 10 equivalents of (left) *holo*-ACP3 or (right) *apo*-ACP3 by adding 1.5 equivalents of DBA under varying citrate concentration. Enhancement of crosslinking efficiency by citrate is consistent with its ability to enhance the rate of chain elongation by DEBS and other PKSs[10,24]. As explained in Fig. S10, *apo*-ACP3 contains small amounts of its *holo*-form counterpart, leading to the appearance of crosslinked products under conditions of vast ACP excess. A “Control” reaction was tested in which the *apo*-ACP3 thiol groups were protected with a maleimide group prior to the addition of DBA (see the Supplementary Methods for details). “Ref.” lanes include all proteins –DBA. Adducts of KS3AT3 with ACP3 (KS3AT3-ACP3) compared to un-crosslinked KS3AT3 were quantified by GelAnalyzer 19.1 and their amounts tabulated as the percentage of total protein per lane. *Residual protein amounts (≤2%) are accounted for by undefined high molecular weight products (MW > KS3AT3-ACP3). For gel source data, see Supplementary Raw Data.

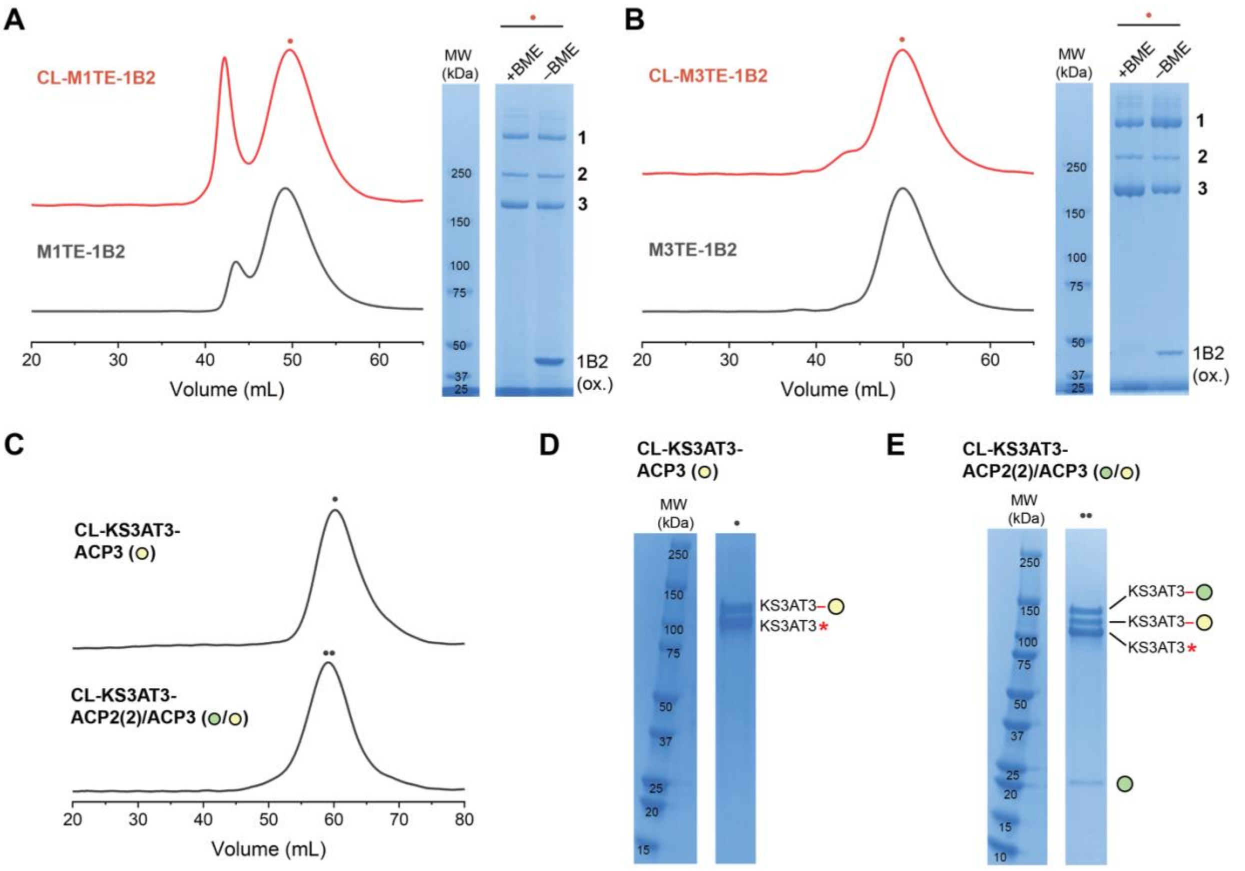

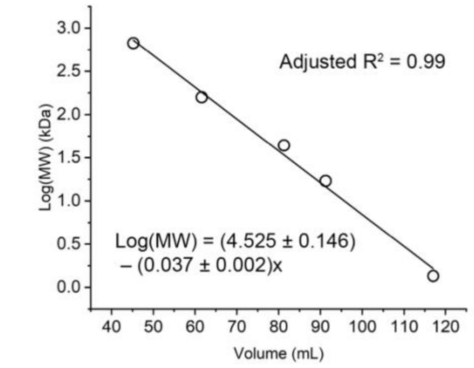

**G**

| Sample | Volume (mL) | MW (kDa) | Calc. MW (kDa) |
|---|---|---|---|
| M1TE-1B2 (dimer) | 49.6 | 480.0 | 498.3 |
| M3TE-1B2 (dimer) | 49.9 | 476.0 | 485.8 |
| KS3AT3–● /KS3AT3*(dimer) | 59.0 | 219.7 | 226.1 |
| KS3AT3–○ /KS3AT3*(dimer) | 60.3 | 210.7 | 202.3 |

**Fig. S12.** Size-exclusion chromatography (SEC) analysis and preparation of each cryo-EM sample used in this study. (**A**, **B**) Chromatograms of native (black) and crosslinked (CL; red) M1TE and M3TE. (Both modules are bound to $F_{ab}$ 1B2.) Reducing and non-reducing SDS-PAGE analysis (± BME) of peaks marked with red dots are also shown. In panel **A**, slight differences in the peak elution volumes probably reflect experimental errors during sample injection that resulted in different starting elution times. (**C**) Analogous SEC chromatograms of homodimeric KS3AT3 crosslinked to ACP3 (top) or to ACP2(2) + ACP3 (bottom). (**D**, **E**) SDS-PAGE analysis for peaks marked with black dots in panel **C**. (**F**) Protein standards (Bio-Rad #1511901) were analyzed by SEC to generate a standard curve. (**G**) Observed and calculated molecular weights and retention volumes of each cryo-EM sample used in this study is tabulated. Red asterisks denote KS3AT3 didomains with a quenched DBA moiety (as explained in Fig. S5A). For gel source data, see Supplementary Raw Data.

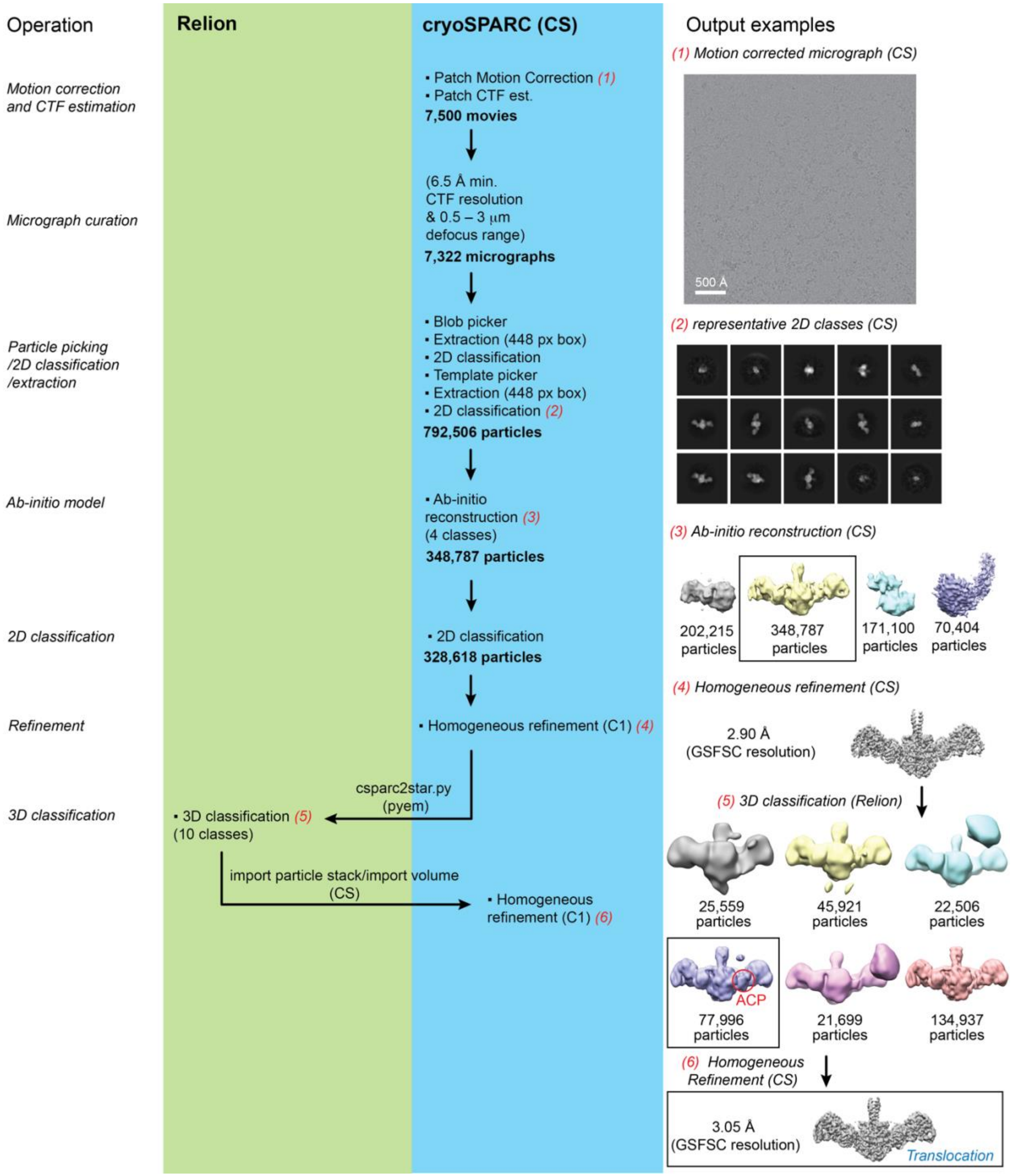

**Fig. S13.** Single-particle cryo-EM analysis of a DBA-crosslinked sample containing homodimeric KS3AT3, ACP2(2), and ACP3 (as prepared in Fig. S12). A combination of processing tools in Relion[13] and cryoSPARC (CS)[14] were implemented to obtain the final cryo-EM map (bottom right).

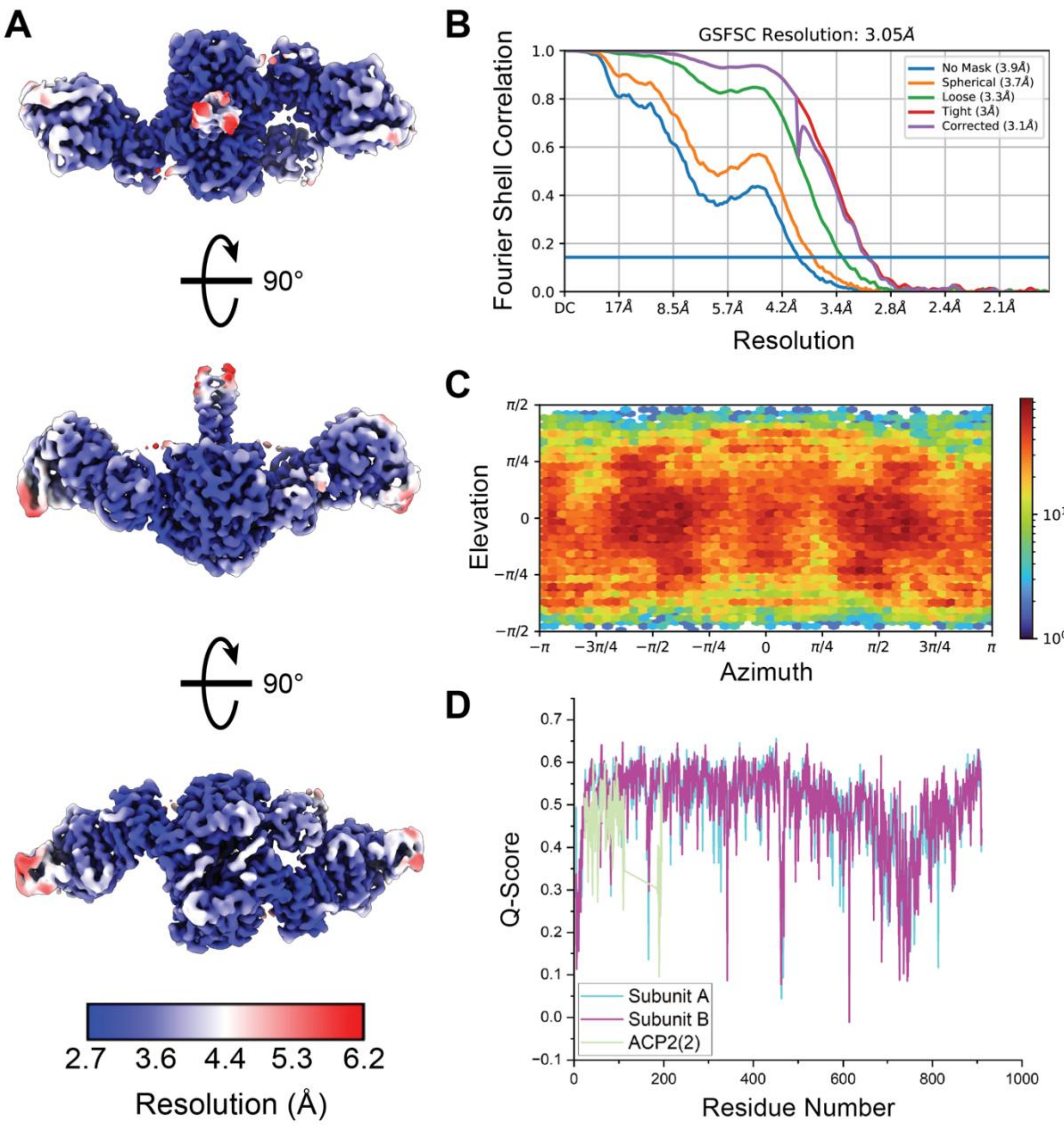

**Fig. S14.** Cryo-EM map and model validation of the *Translocation* structure (Fig. S13). (**A**) Local resolution map of *Translocation* generated in cryoSPARC[14]. (**B**) Curves display the Fourier shell correlation (FSC) between two independently refined half maps at different spatial frequencies (resolution) generated in cryoSPARC[14]. The gold-standard FSC (GSFSC) resolution is defined as the spatial frequency at which the corrected FSC curve intersects with a threshold FSC value of 0.143[25]. (**C**) Euler angle distribution plot generated in cryoSPARC shows a heat map representation of the number of images for each particle orientation[14]. (**D**) The Q-score assessment of atom resolvability within the cryo-EM map is plotted for all residues of the *Translocation* model (PDB 8TKO)[26].

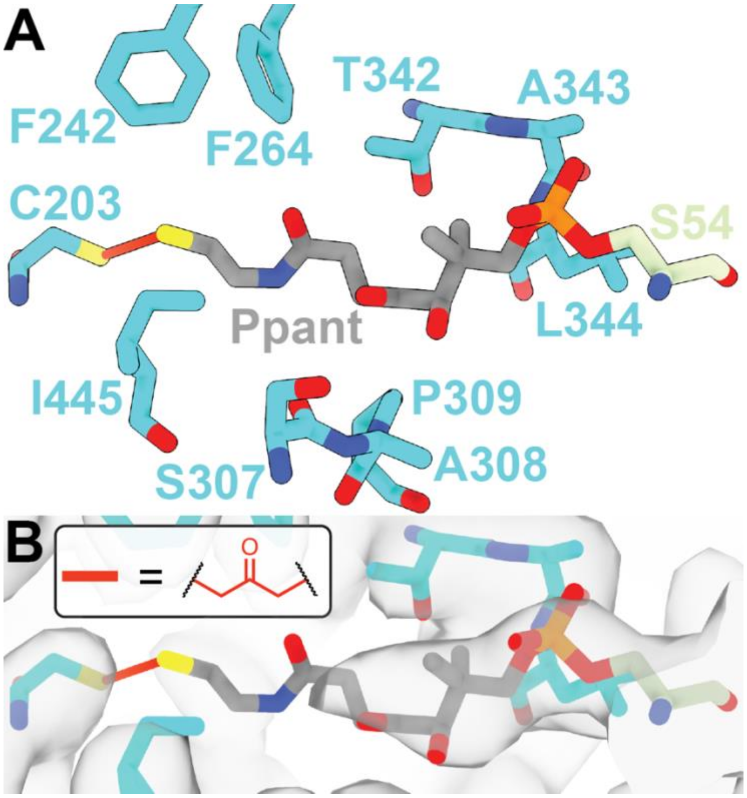

**Fig. S15.** Close-up analysis of the site of DBA-mediated crosslinking between the KS Cys203 and ACP 4′-phosphopantetheine (Ppant) thiols in the *Translocation* structure (PDB 8TKO). (**A**) Residues in the KS active site (cyan) that are ≤5 Å separated from the Ppant cofactor are shown as sticks. (**B**) The cryo-EM map is displayed in the KS active site to highlight partial Ppant cofactor density and lacking density for the DBA-derived crosslink (red line; inset). Crosslinking by DBA is expected to impart a 3 Å separation between the KS (C203) and ACP (Ppant) sulfur atoms, although disorder in this region prevented accurate modeling. The measured interatomic distance between the C203 and Ppant sulfur atoms is 3.6 Å.

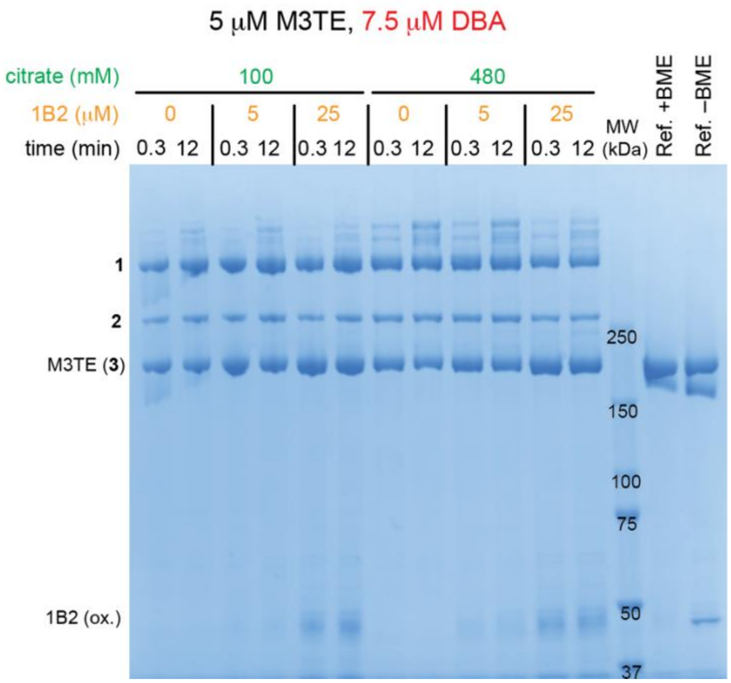

**Fig. S16.** Antibody fragment ($F_{ab}$) 1B2 has no effect on DBA-mediated crosslinking of M3TE. Six different reactions at varying 1B2 (0–25 μM) and citrate (100–480 mM) concentrations were sampled at two different time-points (15 s, 12 min) and analyzed by SDS-PAGE. Reference samples included M3TE + 1B2 ± 2.5% (v/v) BME. For gel source data, see Supplementary Raw Data.

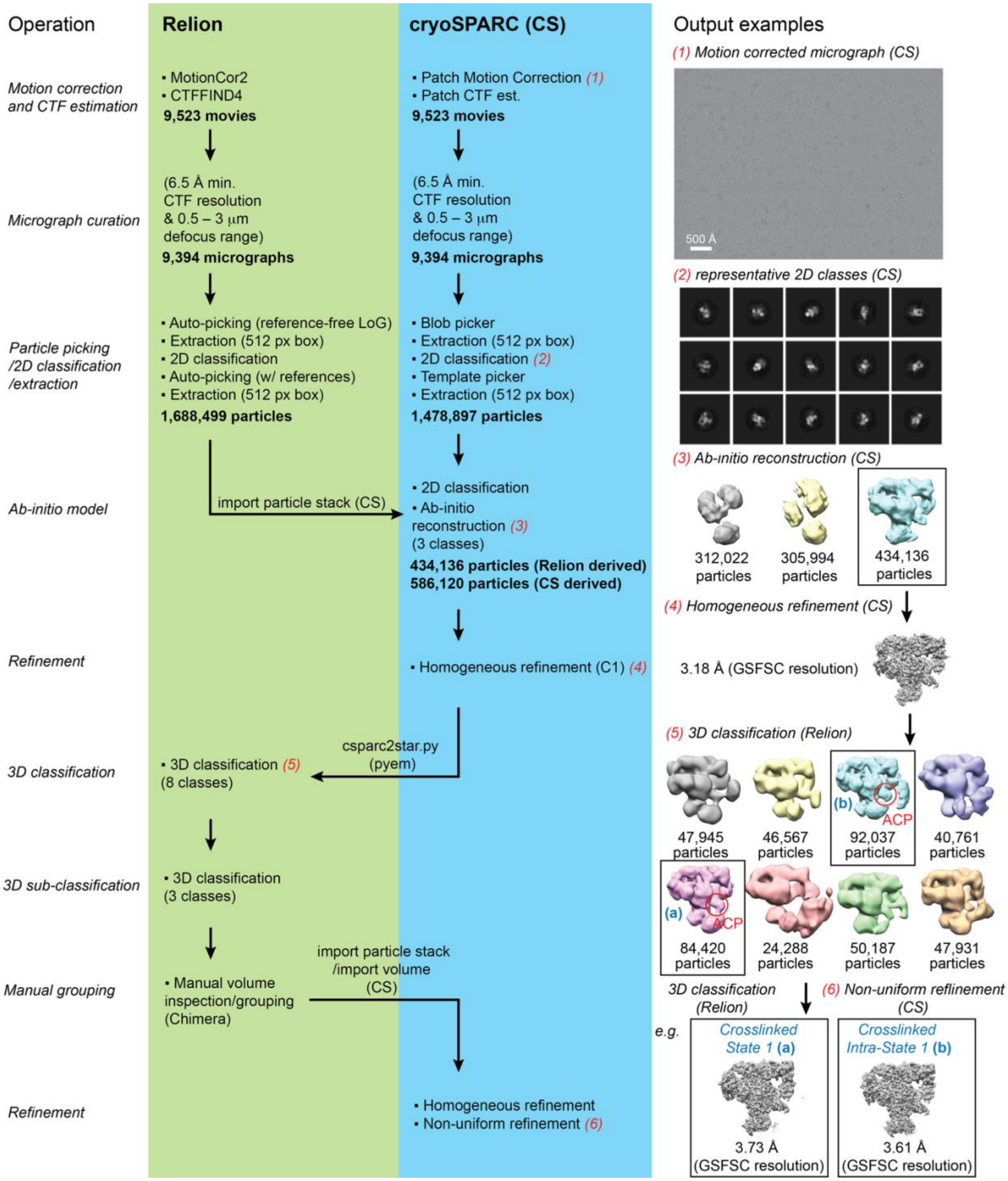

**Fig. S17.** Single-particle cryo-EM analysis of DBA-crosslinked DEBS M1TE in complex with $F_{ab}$ 1B2 (CL-M1TE-1B2). A combination of processing tools in Relion[13] and cryoSPARC (CS)[14] were implemented to obtain the final maps of *Crosslinked State 1* (**a**; see also Fig. S18)) and *Crosslinked Intra-State 1* (**b**; see also Fig. S19) (bottom right).

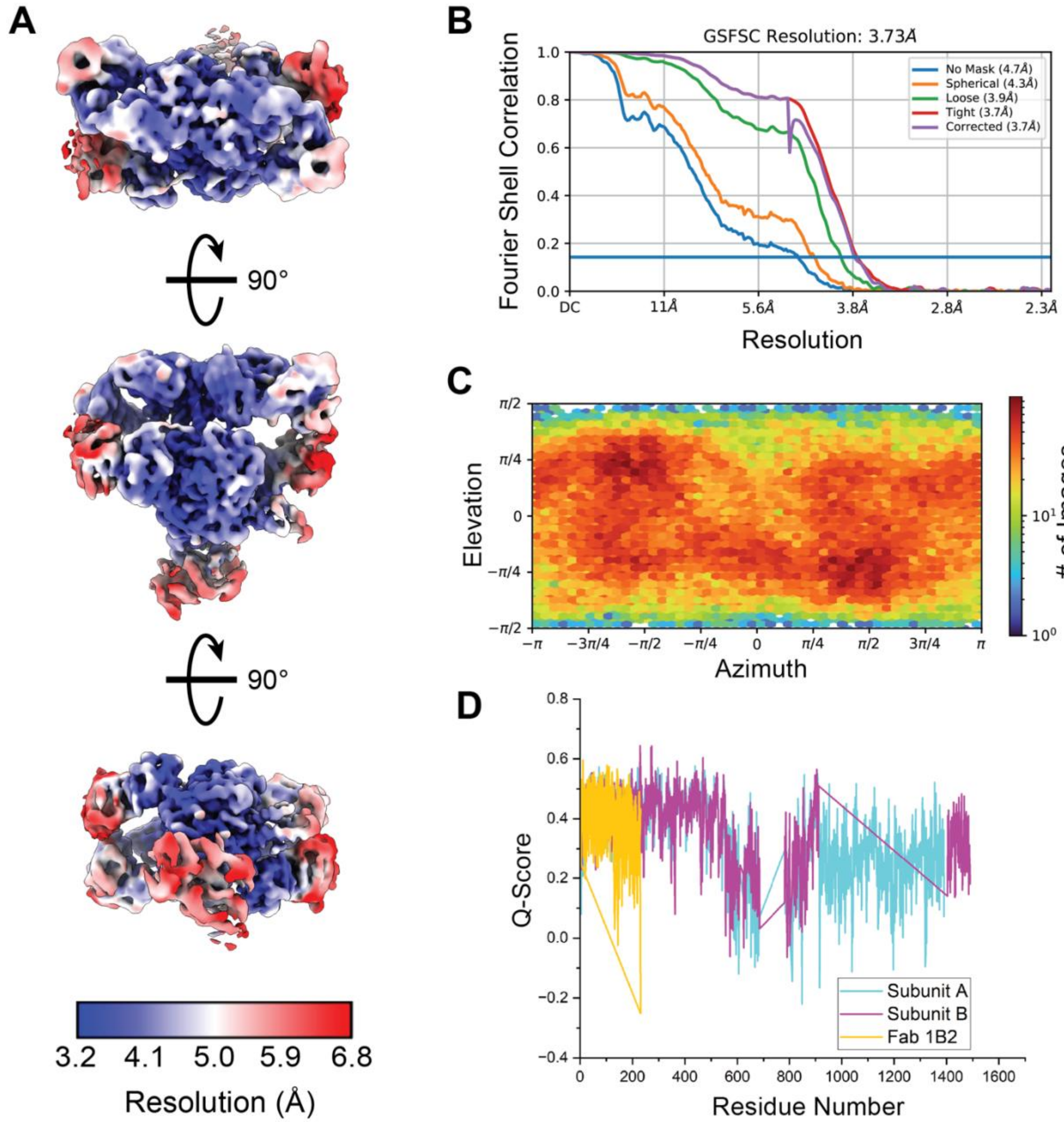

**Fig. S18.** Cryo-EM map and model validation of *Crosslinked State 1* (Fig. S17). (**A**) Local resolution map of *Crosslinked State 1* generated in cryoSPARC[14]. (**B**) Curves display the Fourier shell correlation (FSC) between two independently refined half maps at different spatial frequencies (resolution) generated in cryoSPARC[14]. The gold-standard FSC (GSFSC) resolution is defined as the spatial frequency at which the corrected FSC curve intersects with a threshold FSC value of 0.143[25]. (**C**) Euler angle distribution plot generated in cryoSPARC shows a heat map representation of the number of images for each particle orientation[14]. (**D**) The Q-score assessment of atom resolvability within the cryo-EM map is plotted for all residues of the *Crosslinked State 1* model (PDB 8TJN)[26].

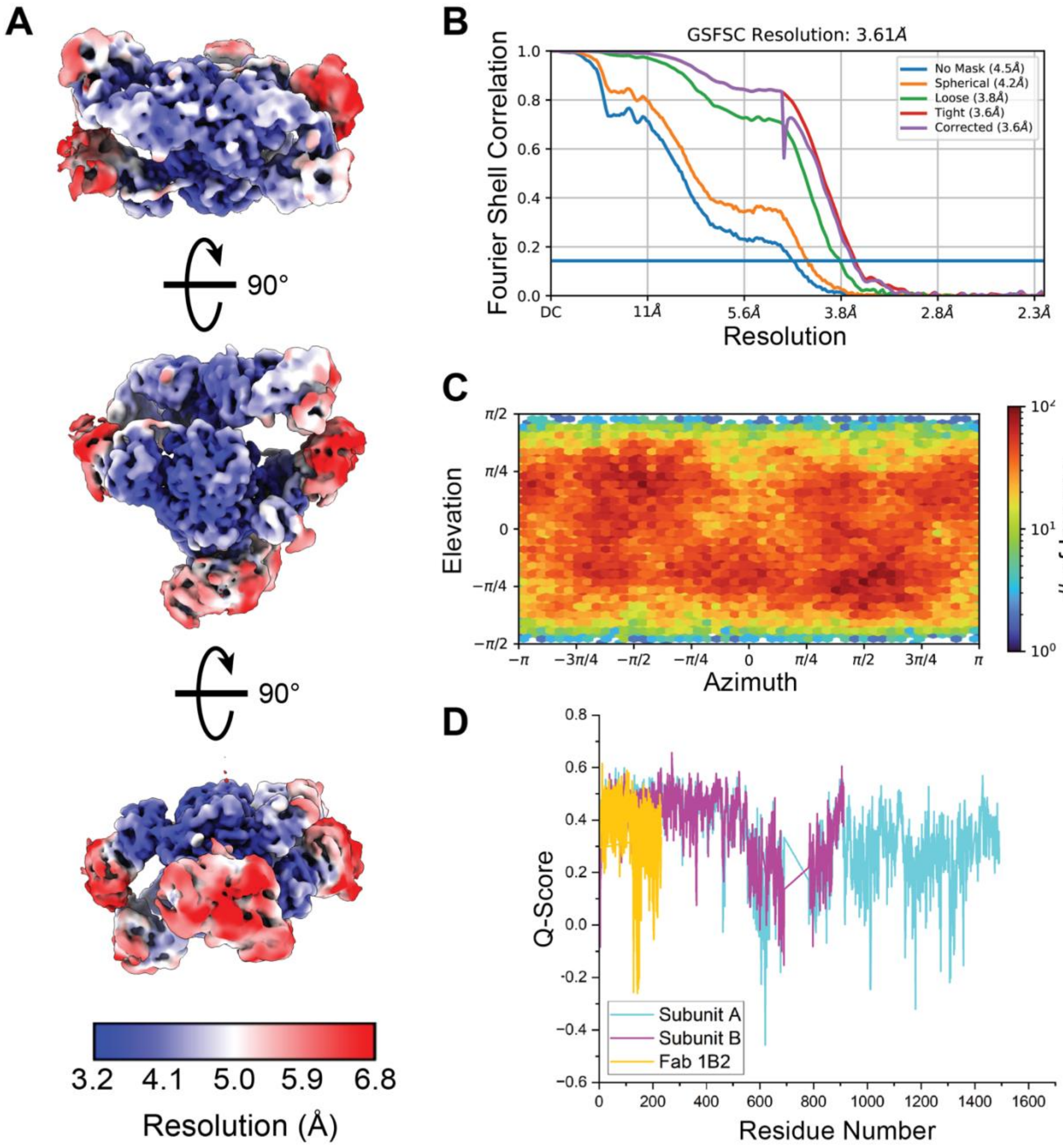

**Fig. S19.** Cryo-EM map and model validation of *Intra-Crosslinked State 1* (Fig. S17). (**A**) Local resolution map of *Intra-Crosslinked State 1* generated in cryoSPARC[14]. (**B**) Curves display the Fourier shell correlation (FSC) between two independently refined half maps at different spatial frequencies (resolution) generated in cryoSPARC[14]. The gold-standard FSC (GSFSC) resolution is defined as the spatial frequency at which the corrected FSC curve intersects with a threshold FSC value of 0.143[25]. (**C**) Euler angle distribution plot generated in cryoSPARC shows a heat map representation of the number of images for each particle orientation[14]. (**D**) The Q-score assessment of atom resolvability within the cryo-EM map is plotted for all residues of the *Intra-Crosslinked State 1* model (PDB 8TJO)[26].

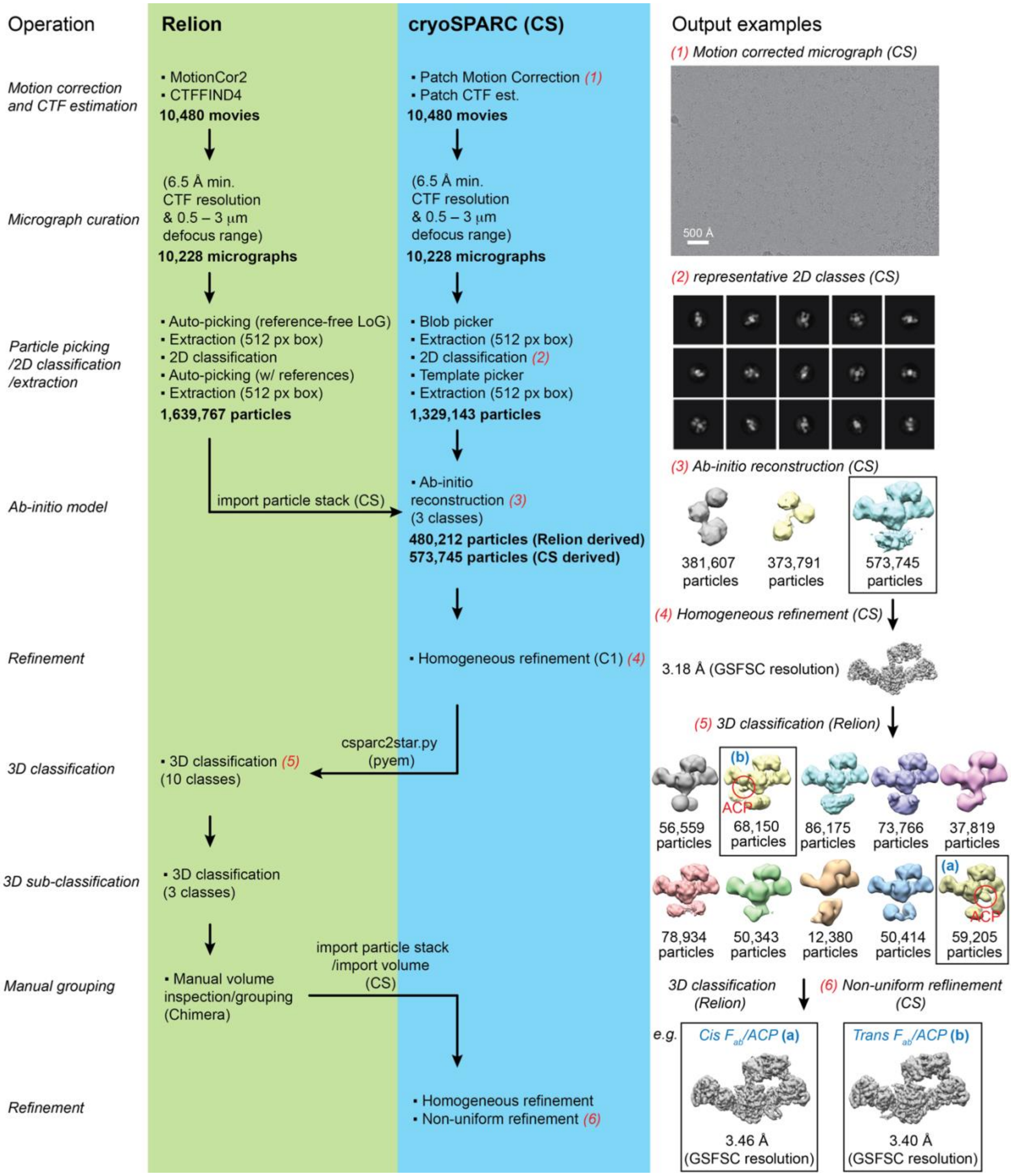

**Fig. S20.** Single-particle cryo-EM analysis of DBA-crosslinked DEBS Module 3 in complex with $F_{ab}$ 1B2 (CL-M3TE-1B2). A combination of processing tools in Relion[13] and cryoSPARC (CS)[14] were implemented to obtain the final maps of *Cis $F_{ab}$/ACP* (**a**; see also Fig. S21) and *Trans $F_{ab}$/ACP* (**b**; see also Fig. S22) (bottom right). Note that regions of density in the final maps cannot be visualized at the set threshold values.

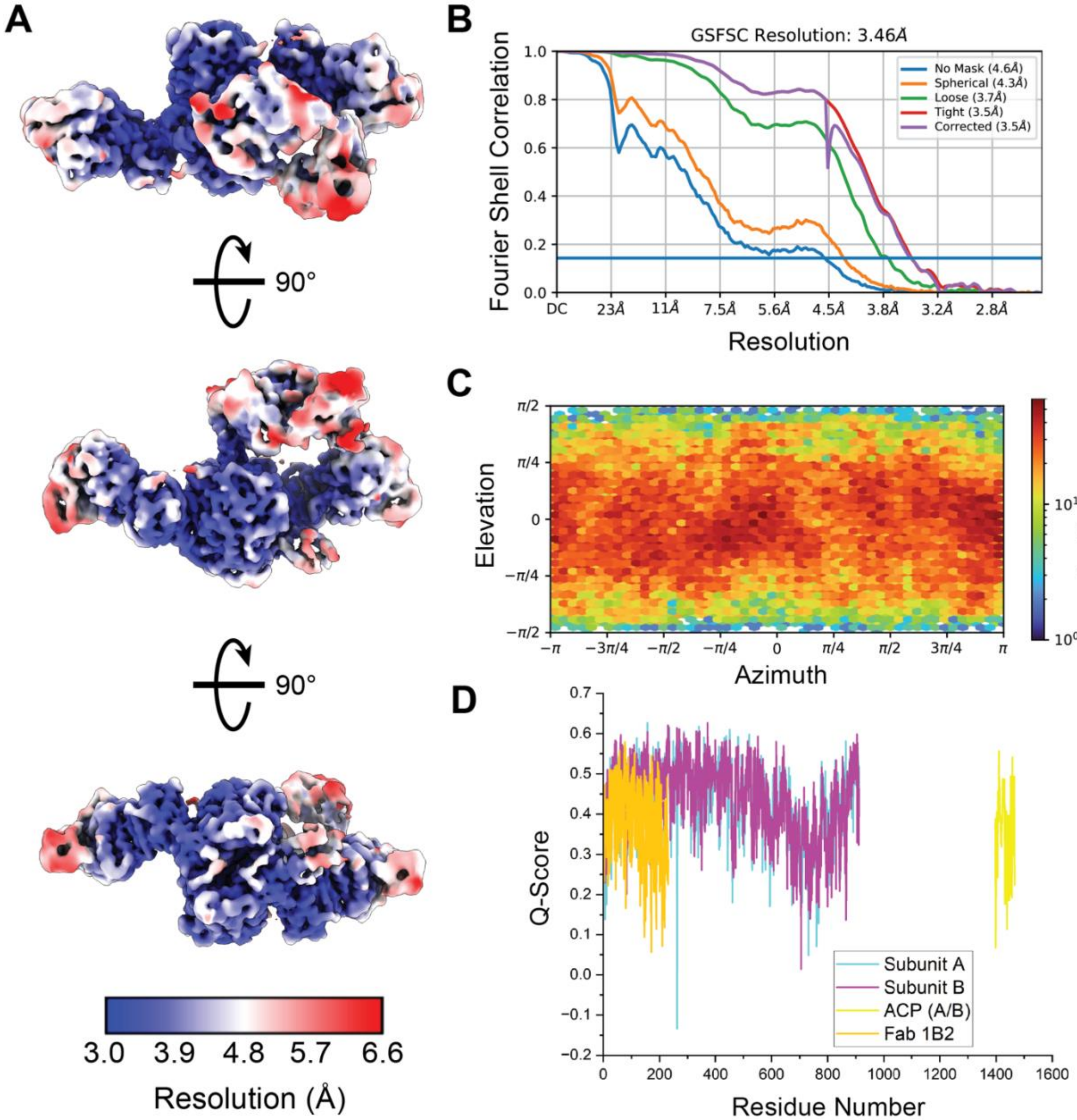

**Fig. S21.** Cryo-EM map and model validation of *Cis F<sub></sub>* *Cis $F_{ab}$/ACP* (Fig. S20). (**A**) Local resolution map of *Cis $F_{ab}$/ACP* generated in cryoSPARC[14]. (**B**) Curves display the Fourier shell correlation (FSC) between two independently refined half maps at different spatial frequencies (resolution) generated in cryoSPARC[14]. The gold-standard FSC (GSFSC) resolution is defined as the spatial frequency at which the corrected FSC curve intersects with a threshold FSC value of 0.143[25]. (**C**) Euler angle distribution plot generated in cryoSPARC shows a heat map representation of the number of images for each particle orientation[14]. (**D**) The Q-score assessment of atom resolvability within the cryo-EM map is plotted for all residues of the *Cis $F_{ab}$/ACP* model (PDB 8TPW)[26].

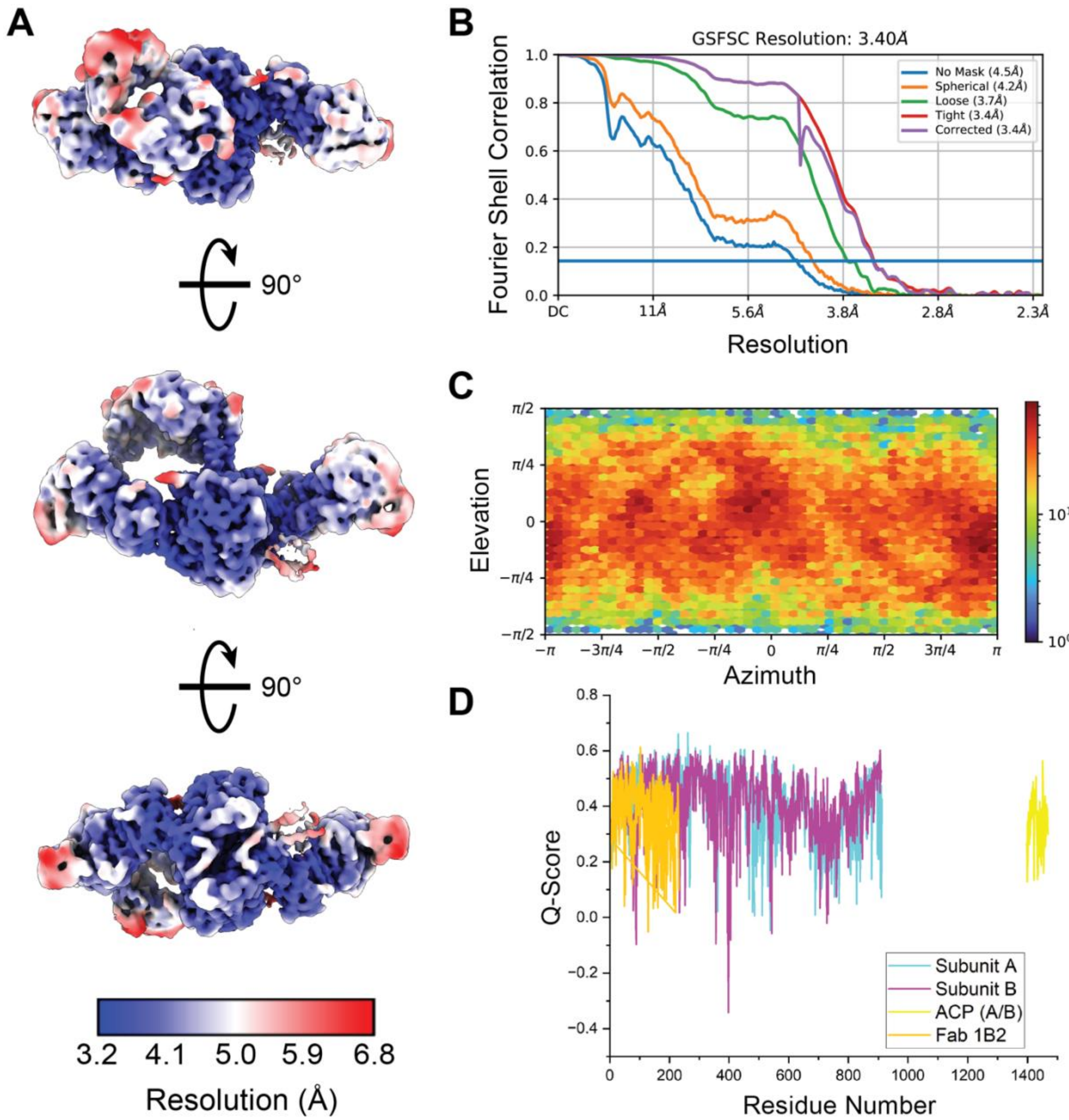

**Fig. S22.** Cryo-EM map and model validation of *Trans $F_{ab}$/ACP* (Fig. S20). (**A**) Local resolution map of *Trans $F_{ab}$/ACP* generated in cryoSPARC[14]. (**B**) Curves display the Fourier shell correlation (FSC) between two independently refined half maps at different spatial frequencies (resolution) generated in cryoSPARC[14]. The gold-standard FSC (GSFSC) resolution is defined as the spatial frequency at which the corrected FSC curve intersects with a threshold FSC value of 0.143[25]. (**C**) Euler angle distribution plot generated in cryoSPARC shows a heat map representation of the number of images for each particle orientation[14]. (**D**) The Q-score assessment of atom resolvability within the cryo-EM map is plotted for all residues of the *Trans $F_{ab}$/ACP* model (PDB 8TPX)[26]. The ACP and its 4′-phosphopantetheine (Ppant) cofactor were poorly resolved by this map.

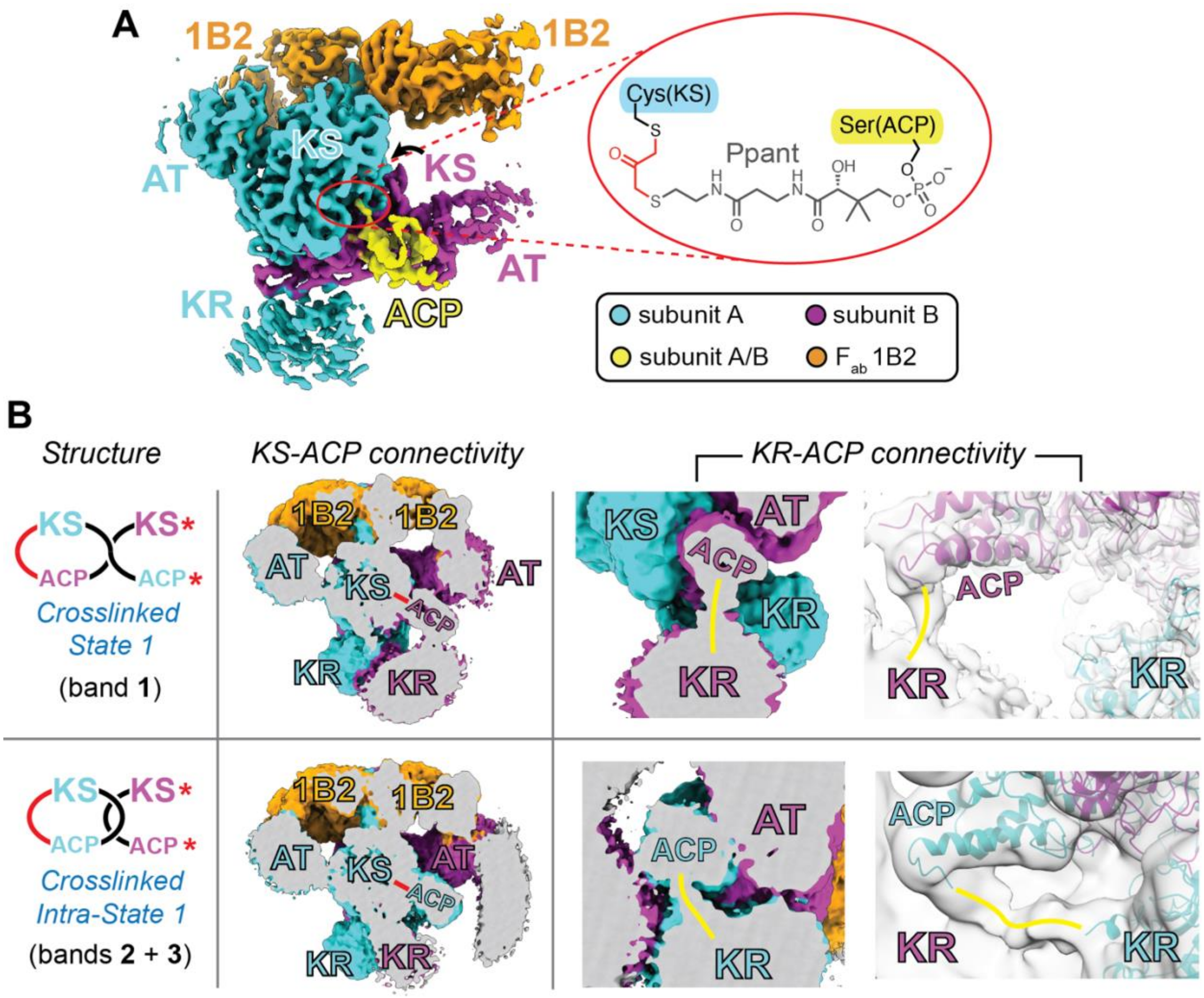

**Fig. S23** Identification of bands **1**–**3** (Fig. 1B) by cryo-EM map analysis. (**A**) Consensus cryo-EM map of the crosslinked DEBS M1TE in complex with $F_{ab}$ 1B2 at 3.18 Å GSFSC resolution (step *(4)* in Fig. S17). Ppant = 4′-phosphopantetheine. (**B**) 3D classification of the data in panel **A** produced two maps that supported the bands **1**–**3** assignments (step *(5)* in Fig. S17). *Crosslinked State 1* (3.73 Å; corresponding to band **1**) contains an intermolecular KS-ACP crosslink, whereas *Crosslinked Intra-State 1* (3.61 Å; corresponding to bands **2** + **3**) contains an intramolecular KS-ACP crosslink. Red arcs and lines denote DBA crosslinks, and red asterisks denote a quenched DBA group (Fig. S5A). Yellow lines denote regions of continuous density used to assign ACP domains to their corresponding subunits in cyan or magenta. To improve the resolvability of the KR-ACP linkers, we applied our recently reported map refinement procedure based on Gaussian mixture models[27]. Two snapshots of these maps are shown on the right-most column under *"KR-ACP connectivity"* in panel **B**.

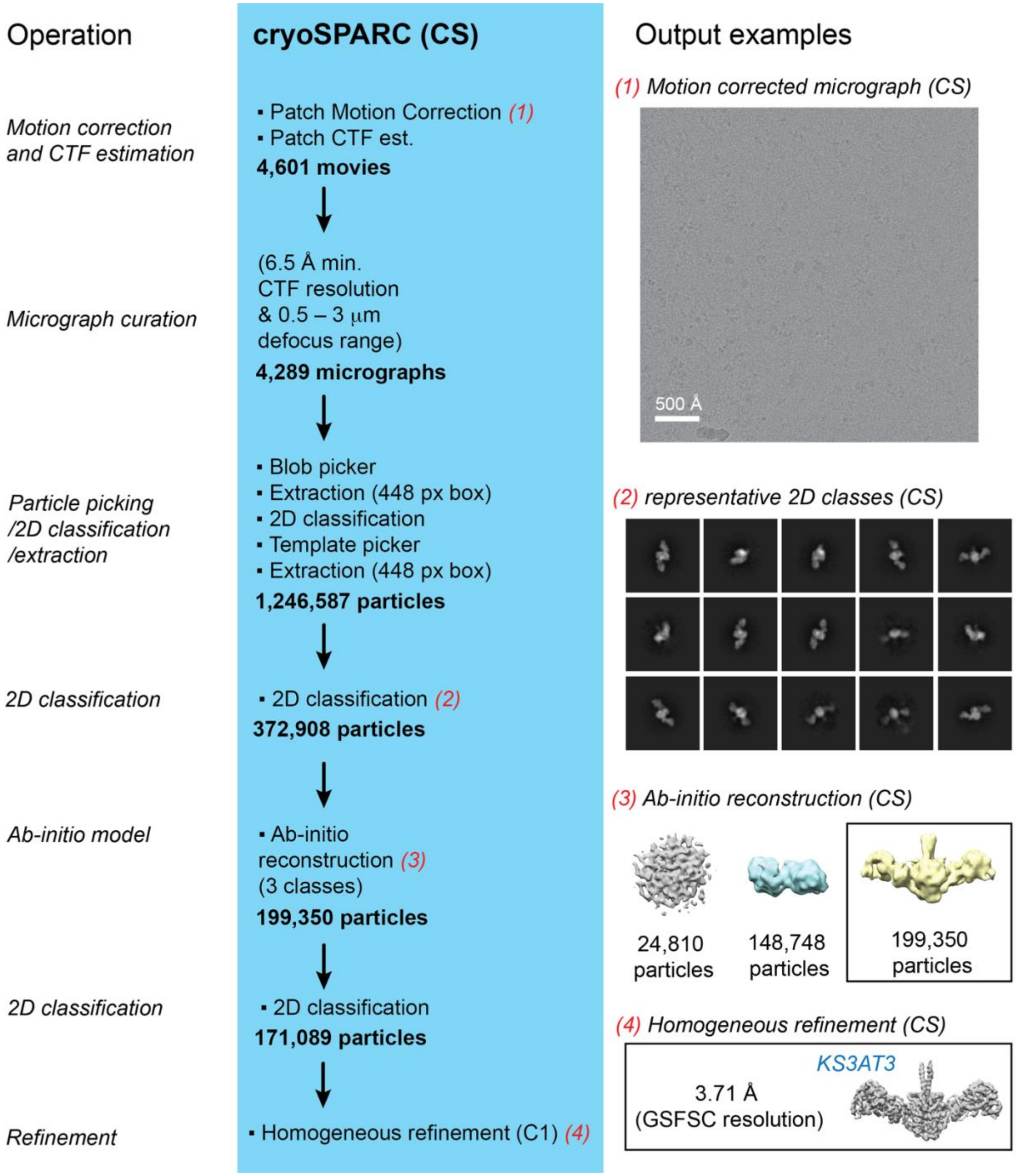

**Fig. S24.** Single-particle cryo-EM analysis of DBA-crosslinked DEBS KS3AT3-ACP3. A combination of processing tools in cryoSPARC (CS)[14] were implemented to obtain the final cryo-EM map (bottom right). The second *ab initio* volume from step *(3)* appears to be a KS-AT monomer that could not be refined to greater than 7 Å GSFSC resolution.

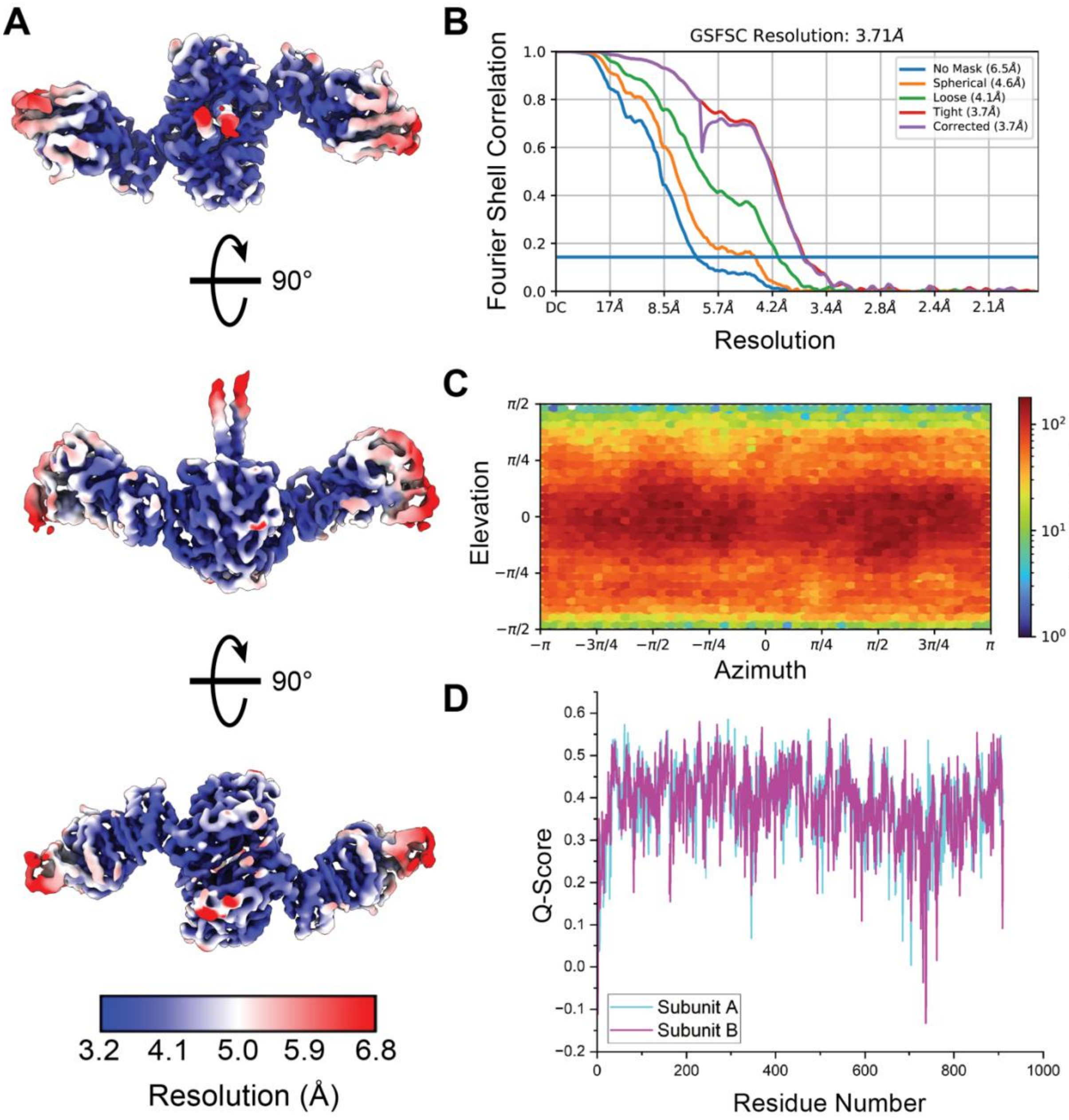

**Fig. S25.** Cryo-EM map and model validation of the *KS3AT3* structure (Fig. S24). (**A**) Local resolution map of *KS3AT3* generated in cryoSPARC[14]. (**B**) Curves display the Fourier shell correlation (FSC) between two independently refined half maps at different spatial frequencies (resolution) generated in cryoSPARC[14]. The gold-standard FSC (GSFSC) resolution is defined as the spatial frequency at which the corrected FSC curve intersects with a threshold FSC value of 0.143[25]. (**C**) Euler angle distribution plot generated in cryoSPARC shows a heat map representation of the number of images for each particle orientation[14]. (**D**) The Q-score assessment of atom resolvability within the cryo-EM map is plotted for all residues of the *KS3AT3* model (PDB 8TJP)[26].

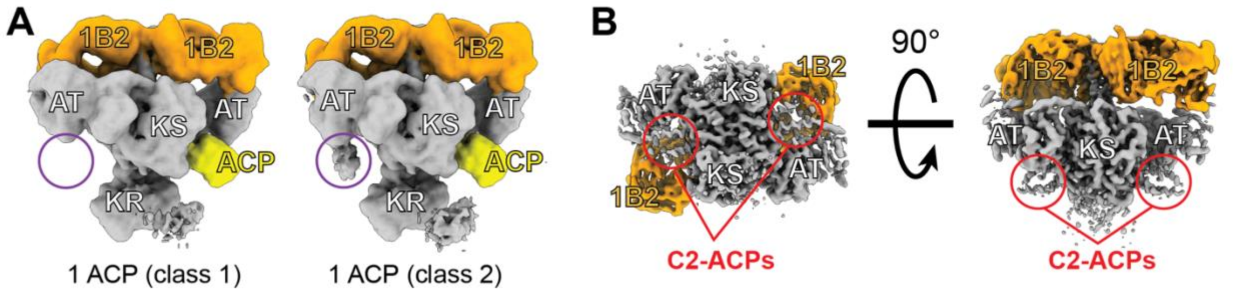

**Fig. S26.** Statistical per-particle image analysis of crosslinked Module 1 of DEBS (M1TE) bound to $F_{ab}$ 1B2 (CL-M1TE-1B2) (supplement to Fig. 5B). (**A**) Alternative views of the '1 ACP' class 1 and class 2 in Fig. 5B show an undefined region of density near the AT domain (circled in purple) that is unique to class 2 and resulted in their segregation during 3D focused classification. (**B**) To further analyze this observation, the particles and volume from example *"(3) Ab initio Reconstruction"* (Fig. S17) were subjected to homogeneous C2-refinement[14]. Visualization of the C2-refined map at two 90°-related orientations highlights double ACP occupancy resulting from imposed symmetry. Statistical per-particle image analysis, however, was not supportive of this state, as no more than one ACP bound to a KS active site was detected in any of the particles by this method (Fig. 5B and panel **A**, above).

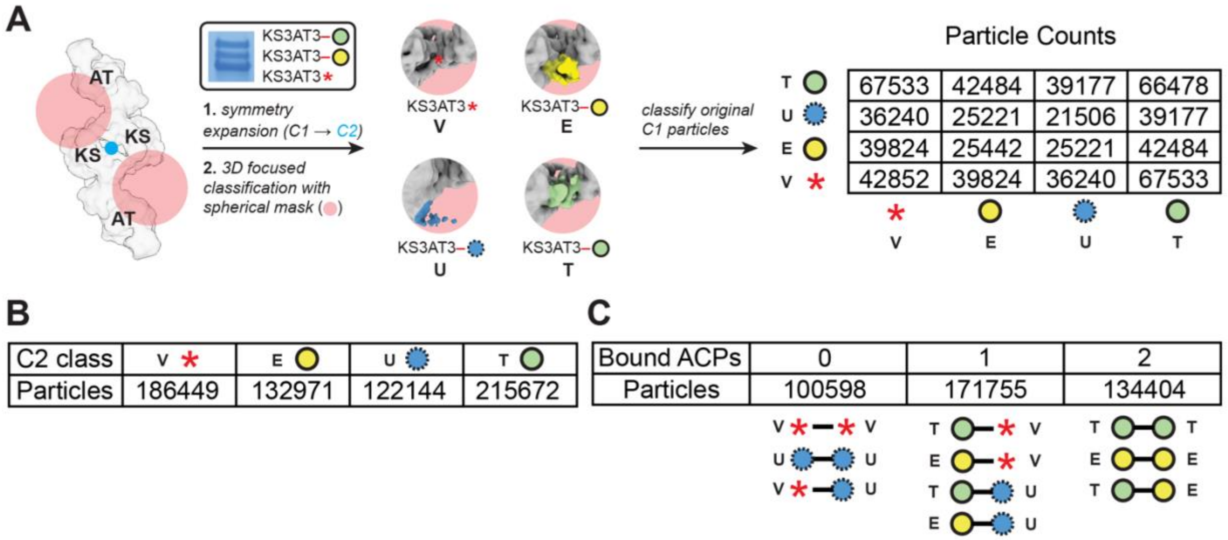

**Fig. S27.** Statistical per-particle image analysis of cryo-EM data following crosslinking of KS3AT3 with ACP2(2) (green circle) and ACP3 (yellow circle). The boxed inset in panel **A** shows SDS-PAGE analysis of the crosslinked material isolated as a single peak by SEC for cryo-EM analysis (Fig. S12). Red asterisks denote the KS3AT3 protein whose KS active site harbors a quenched DBA moiety (Fig. S5A). (**A**) Consensus particles were (**1**) symmetry-expanded about the pseudo-C2 axis of symmetry (blue dot) and (**2**) sorted according to differences in density at the ACP binding sites by 3D focused classification. The C2-refined particles were binned into four classes: two to account for ACP2(2) (designated **T** for translocation) and ACP3 (designated **E** for elongation), one to account for a vacant cleft (**V**), and a fourth to capture any unexpected structural features (**U**). (Class **U** may correspond to an average representation of crosslinked but dissociated ACPs.) We then correlated the C2-refined particles to their symmetry mates, as in Figure 5, to quantify each homo- or hetero-dimeric species. (**B**) The particle counts for each C2 class following step (**2**) in panel **A**. (**C**) Particle counts of the classified C1 particles in panel **A** were determined after sorting them into groups according to their number of bound ACPs, assuming classes **E** and **T** corresponded to ACP-bound states and classes **V** and **U** corresponded to unbound states. For gel source data, see Supplementary Raw Data.

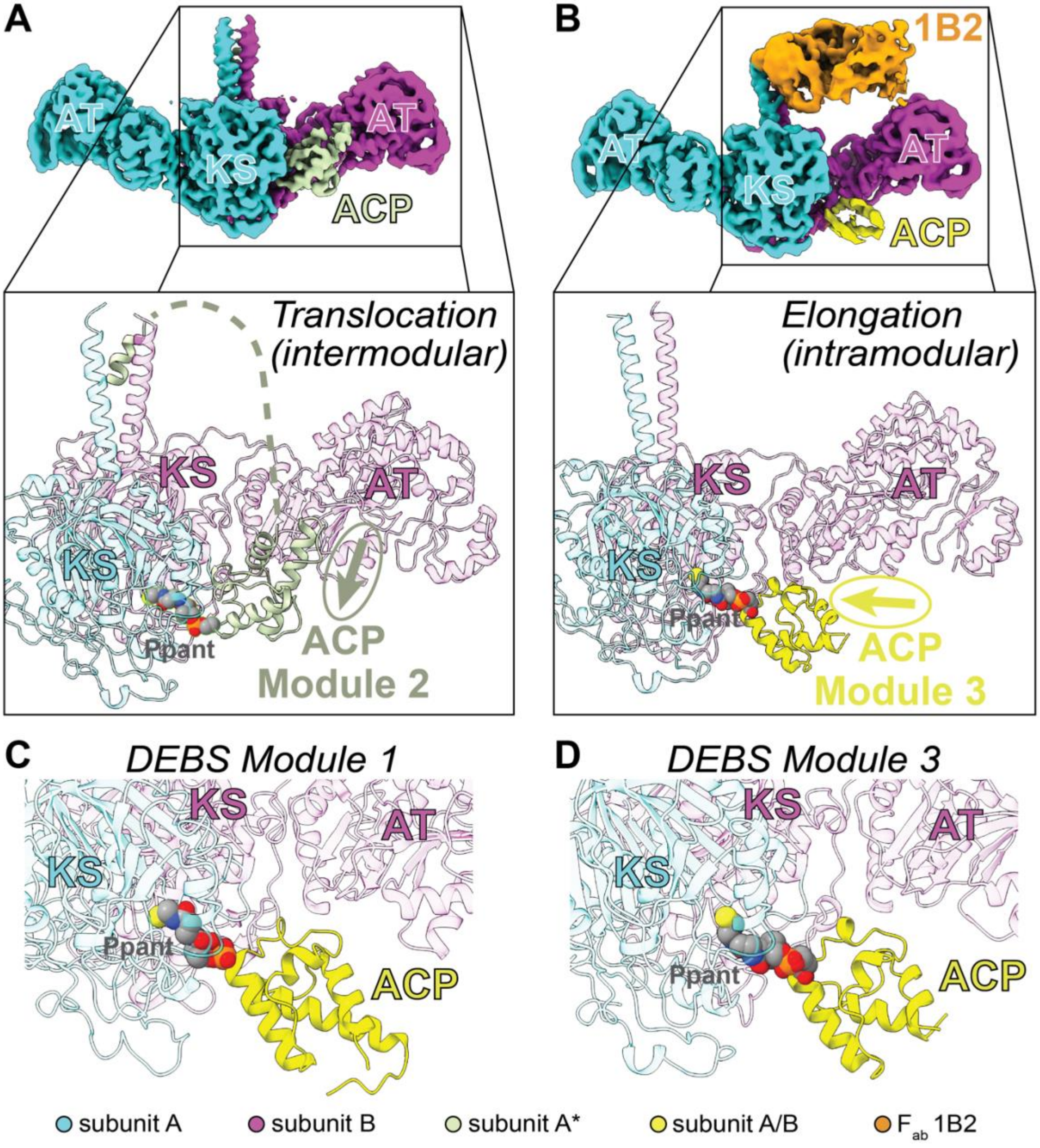

**Fig. S28.** (**A**, **B**) Comparison of the (**A**) translocation (Fig. 3) and (**B**) elongation (Fig. 4) ACP binding modes of DEBS Module 3. A circled arrow points N- to C-terminally along helix 1 of the ACPs in each structure to emphasize their unique orientations. The ACP from Module 2 in panel **A**, ACP2(2), also contains a C-terminal docking domain that forms a 3-helix bundle with the complementary coiled-coil docking domain of Module 3 (67 residues separating the ACP and its C-terminal docking domain were unresolved; depicted as a dashed line). (**C**, **D**) Intramodular ACP-KS interactions during elongation are similar in DEBS (**C**) Module 1 and (**D**) Module 3. Atoms of the 4′-phosphopantetheine (Ppant) cofactor are represented as spheres. Relative differences in the Ppant conformations in panels C and D reflect uncertainties in the atomic coordinates of the Ppant groups due to reduced local resolution in the KS active site (>3.5 Å; Fig. S21A).

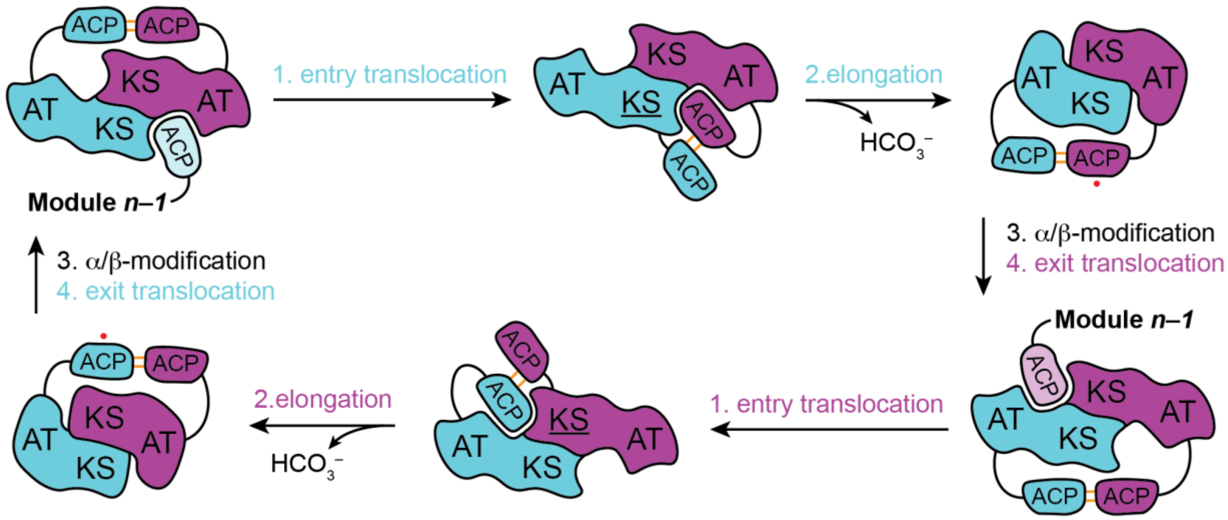

**Fig. S29.** Proposed model for coupling between polyketide translocation and elongation catalyzed by a homodimeric PKS module. The catalytic cycle consists of two equivalent half-cycles (steps 1–4) carried out asynchronously by opposite subunits. Coupling of thermoneutral translocation and exergonic elongation promotes a change in specificity of the KS domain for its upstream and internal ACPs, respectively. Pre-elongation, the ACPs are sequestered to one KS-AT cleft, consistent with the formation of bands **1** and **2** (Fig. 1B). Post-elongation, the KS-AT cleft volumes are constricted by the turnstile mechanism (Fig. S2) to prevent premature acyl-ACP substrate entry during $\alpha$/$\beta$-modification and exit translocation[2]. Although both KS-AT clefts are depicted as closed to ACP entry, we point out that possible module occupancy with an upstream donor ACP during or following elongation might physically occlude cleft constriction such that an asymmetric state is formed — akin to our previously observed structure (PDB 7M7G). We propose constriction of both KS-AT clefts in accordance with the previous “turnstile closed” structure (PDB 7M7J) which was obtained in the absence of an upstream donor ACP[10]. Underlining of KS domains indicates acylation of their active site Cys residues with electrophilic substrates. Red dots on ACP domains denote that the Ppant arm is bound to a post-elongation polyketide intermediate that is optionally modified at its $\alpha$- and $\beta$-positions by other catalytic domains of the module (not shown for clarity). Two orange lines connect the ACPs at each stage to depict non-covalent forces predicted to hold them together (i.e., dimeric elements of the C-terminal docking or TE domains)[28]. The fact that each state has no more than one underlined KS domain or red dot is consistent with our previous finding that a homodimeric module is occupied by no more than one polyketide intermediate under steady-state turnover conditions[2].

#### Supplementary Tables

| **Structure** | **PDB ID** | **Resolution (Å, GSFSC)** | **Sample** | **Expression Plasmid(s)** |
|---|---|---|---|---|
| *State 1* | 7M7F | 3.2 | DEBS M1TE + 1B2 | pDC1[10] |
| *State 1'* | 7M7H | 4.1 | DEBS M1TE + 1B2 | pDC1[10] |
| *State 2* | 7M7G | 4.1 | DEBS M1TE + 1B2 | pDC1[10] |
| *Turnstile Closed* | 7M7J | 4.3 | DEBS M1* + 1B2 | pDC7[10] |
| *Module 3/1 Hybrid* | 7M7E | 3.2 | DEBS M3/1TE + 1B2 | pTED23[9] |
| *Translocation* | 8TKO | 3.1 | DEBS KS3AT3 + ACP2(2) + ACP3 | pAYC02[29] + pNW6[3] + pDC9 |
| *Crosslinked State 1* | 8TJN | 3.7 | DEBS M1TE + 1B2 | pDC1[10] |
| *Crosslinked Intra-State 1* | 8TJO | 3.6 | DEBS M1TE + 1B2 | pDC1[10] |
| *Cis $F_{ab}$/ACP* | 8TPW | 3.5 | DEBS M3TE + 1B2 | pRSG34[4] |
| *Trans $F_{ab}$/ACP* | 8TPX | 3.4 | DEBS M3TE + 1B2 | pRSG34[4] |
| *KS3AT3* | 8TJP | 3.7 | DEBS KS3AT3 + ACP3 | pAYC02[29] + pDC9 |

**Table S1.** Summary of near-atomic resolution structures of intact DEBS modules solved to date and their composition. GSFSC = gold-standard Fourier shell correlation. *DEBS M1 was also fused to the N-terminal docking domain of Module 3 for complexation with $F_{ab}$ 1B2 and covalently bound to its native diketide product after incubation with substrates[10].

| Plasmid Name | Encoded Protein | Antibiotic Resistance | Reference |
|---|---|---|---|
| pBL12 | DEBS LDD(4) | Km | [1] |
| pBL13 | DEBS (5)M1(2) | Cb/Am | [1] |
| pDC1 | DEBS M1TE † | Cb/Am | [10] |
| pBL16 | DEBS (3)M2TE | Cb/Am | [1] |
| pRSG34 | DEBS M3TE † | Cb/Am | [4] |
| pNW6 | DEBS ACP2(2) | Km | [3] |
| pNW7 | DEBS ACP2 | Km | [3] |
| pDC9 | DEBS ACP3 | Cb/Am | *This study* |
| pAYC02 | DEBS KS3AT3 | Cb/Am | [29] |
| PrpE-pET28 | PrpE | Km | [2] |
| *SCME-pET28 | SCME | Km | [30] |
| *MatB-pET28 | MatB | Km | [30,31] |
| pRSG56 | Sfp | Km | [4] |
| n/a | $F_{ab}$ 1B2 | Cb/Am | [32] |

*A gift from Prof. Michelle Chang's lab (Princeton University)
†The encoded proteins contain a C-terminal TE domain and the N-terminal docking domain from DEBS M3 (see **Protein Sequences** above).

**Table S2.** Plasmids used in this study (Km = kanamycin; Cb = carbenicillin; Am = ampicillin).

| Sample | CL-M1TE-1B2 | CL-M3TE-1B2 | CL-KS3AT3-ACP3 | CL-KS3AT3-ACP2(2)/ACP3 |
|---|---|---|---|---|
| **Module expression plasmid** | pDC1 | pRSG34 | pAYC02 & pDC9 | pAYC02, pNW6, & pDC9 |
| **Microscope** | Titan Krios G3i | Titan Krios G3i | Titan Krios G3i | Titan Krios G3i |
| **Voltage (kV)** | 300 | 300 | 300 | 300 |
| **Camera** | K3 | K3 | Falcon4 | Falcon4 |
| **Magnification** | 81,000 | 81,000 | 130,000 | 130,000 |
| **Pixel size (Å)** | 1.100 | 1.257 | 0.946 | 0.946 |
| **Total Dose (e- / Å$^2$)** | 50 | 50 | 50 | 50 |
| **Exposure time (s)** | 3.76 | 2.79 | 6.45 | 6.45 |
| **Dose rate (e- pixel$^{-1}$ s$^{-1}$)** | 16.1 | 21.7 | 6.94 | 6.94 |
| **Defocus range during data collection (μm)** | -1.0 – -2.5 | -1.0 – -2.5 | -1.0 – -2.5 | -1.0 – -2.5 |
| **Number of micrographs** | 9,523 | 10,480 | 4,601 | 7,500 |
| **Symmetry** | C1 | C1 | C1 | C1 |
| **Number of pre-refinement particles** | 434,136 | 573,745 | 199,350 | 348,787 |

**Table S3.** Cryo-EM data collection parameters.

| Sample | CL-M1TE-1B2 | CL-M1TE-1B2 | CL-M3TE-1B2 | CL-M3TE-1B2 | CL-KS3AT3-ACP3 | CL-KS3AT3-ACP2(2)/ACP3 |
|---|---|---|---|---|---|---|
| **Structure** | *Crosslinked State 1* | *Crosslinked Intra-State 1* | *Cis $F_{ab}$/ACP* | *Trans $F_{ab}$/ACP* | *KS3AT3* | *Translocation* |
| **PDB ID** | 8TJN | 8TJO | 8TPW | 8TPX | 8TJP | 8TKO |
| **EMDB ID** | EMD-41305 | EMD-41306 | EMD-41495 | EMD-41496 | EMD-41307 | EMD-41355 |
| **Resolution (0.143 FSC, Å)** | 3.73 | 3.61 | 3.46 | 3.40 | 3.71 | 3.05 |
| **Clashscore (all atoms)** | 11.43 | 11.31 | 7.30 | 14.25 | 4.65 | 6.16 |
| **Poor rotamers (%)** | 1.55 | 0.43 | 1.46 | 0.73 | 0.73 | 1.25 |
| **Ramachandran outliers (%)** | 0.38 | 0.34 | 0.04 | 0.76 | 0.06 | 0.05 |
| **Ramachandran favored (%)** | 92.95 | 91.02 | 95.63 | 85.16 | 96.90 | 95.28 |
| **MolProbity score** | 2.17 | 2.09 | 1.83 | 2.32 | 1.42 | 1.74 |
| **Bond length (RMSD, Å)** | 0.003 | 0.003 | 0.003 | 0.003 | 0.003 | 0.002 |
| **Bond angles (RMSD, °)** | 0.599 | 0.698 | 0.495 | 0.692 | 0.604 | 0.485 |

**Table S4.** Single-particle cryo-EM model refinement parameters.
