## Supplementary Raw Data for "Structural Basis for Intermodular Communication in Assembly-Line Polyketide Biosynthesis"

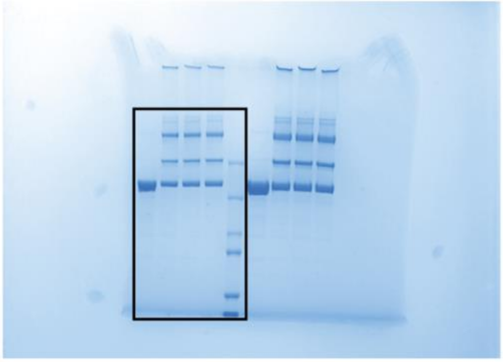

Fig. 1

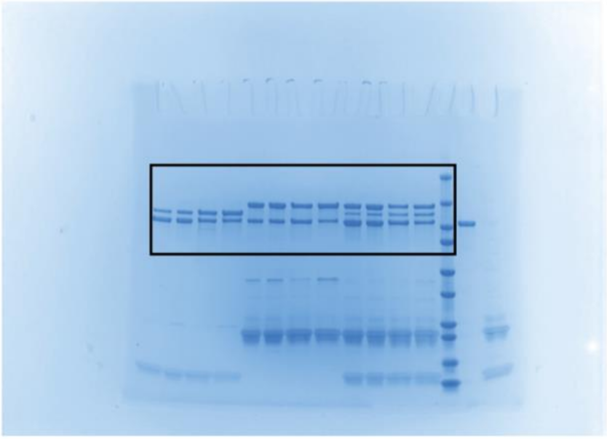

Fig. 2

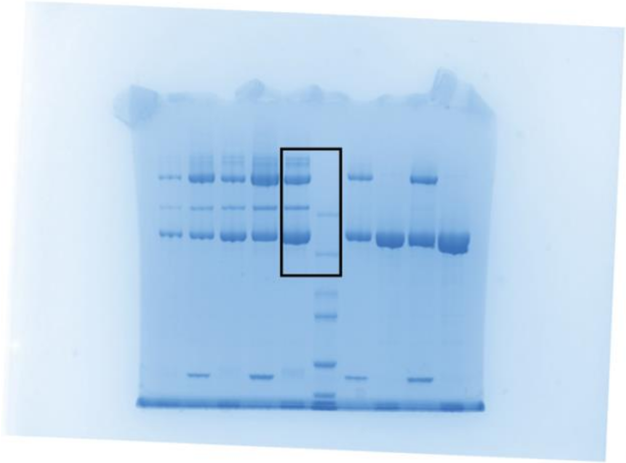

Fig. 3

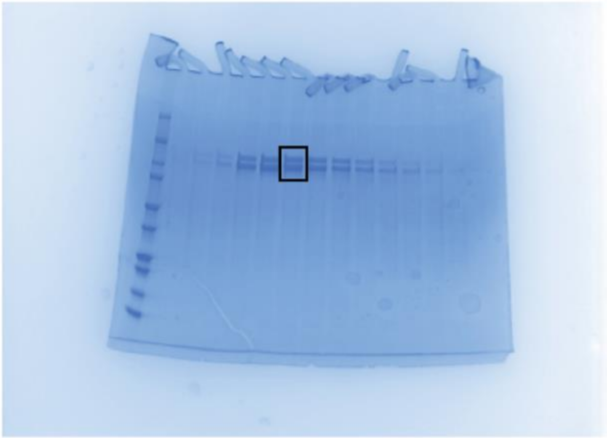

Fig. 4

**Raw Data 1:** Uncropped, raw gel images that were used to create Figs. 1–4. A black box indicates where the gel image was cropped for its presentation in the Figure.

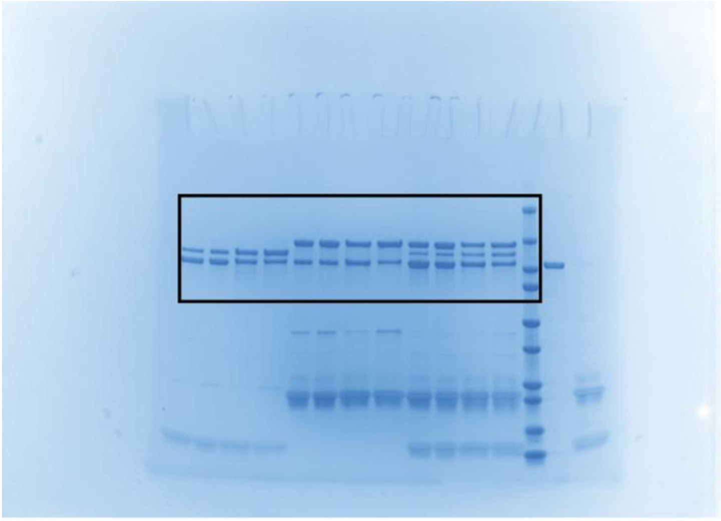

Fig. 2 (replicate 1, displayed)

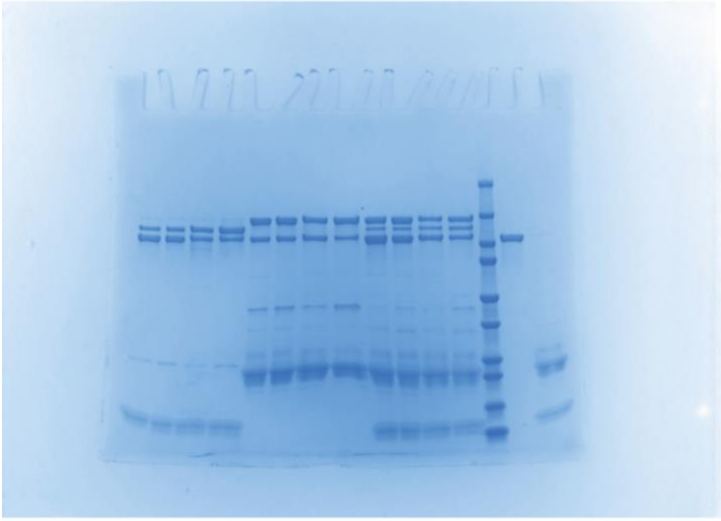

Fig. 2 (replicate 2, not displayed)

**Raw Data 2:** Uncropped, raw gel images that were used to create Fig. 2. A black box indicates where the gel image was cropped for its presentation in the Figure.

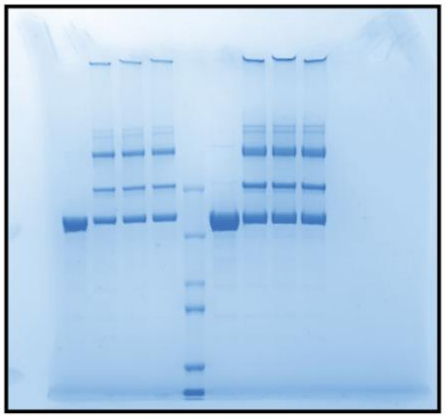

Panel A

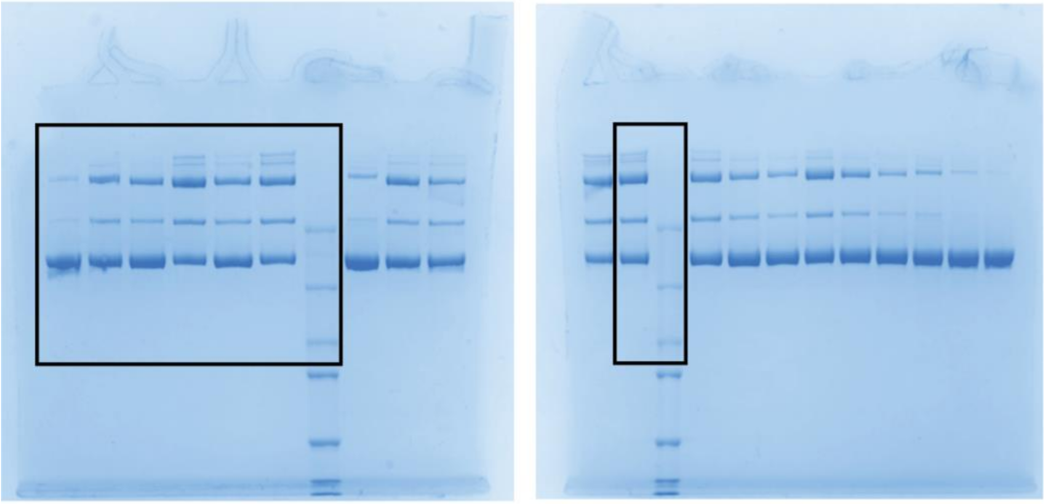

Panel B

**Raw Data 3:** Uncropped, raw gel images that were used to create Fig. S3. A black box indicates where the gel image was cropped for its presentation in the Figure.

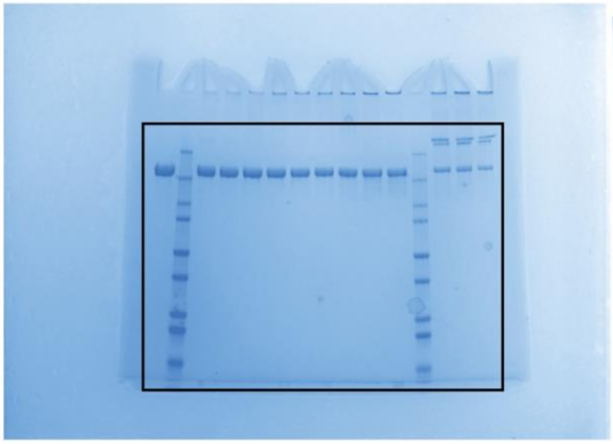

Panel A

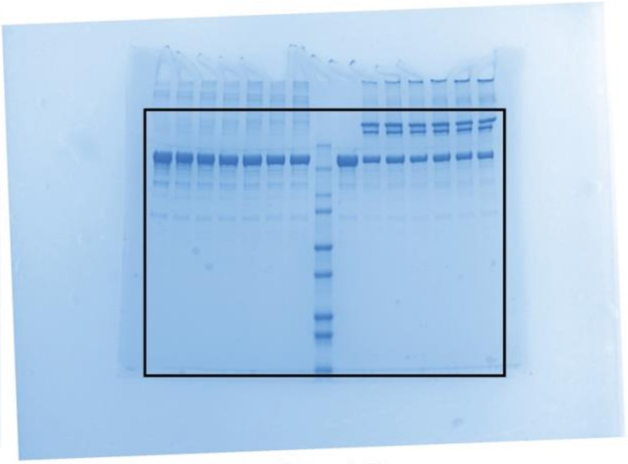

Panel B

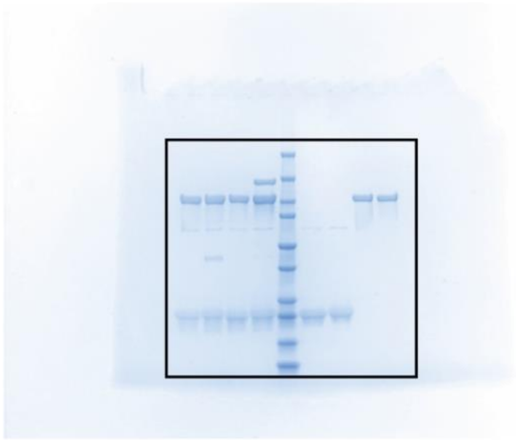

Panel C

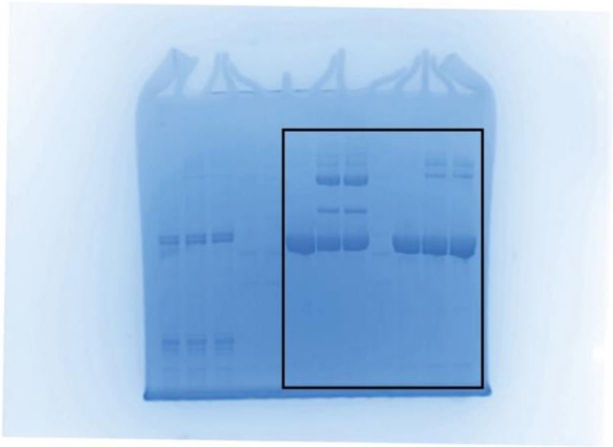

Panel D

**Raw Data 4:** Uncropped, raw gel images that were used to create Fig. S4. A black box indicates where the gel image was cropped for its presentation in the Figure.

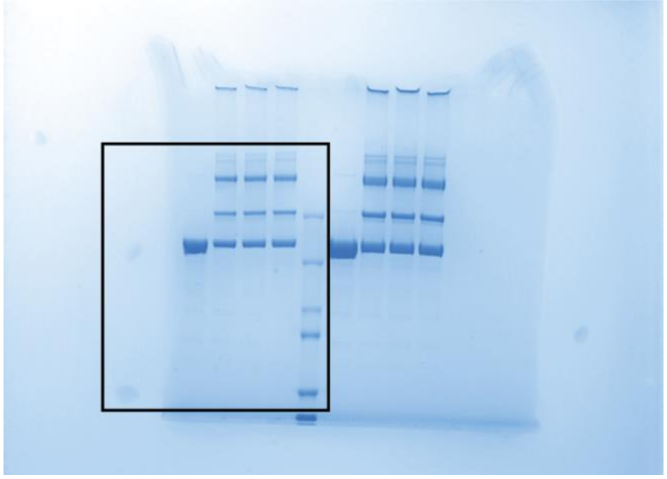

**Raw Data 5:** Uncropped, raw gel images that were used to create Fig. S5. A black box indicates where the gel image was cropped for its presentation in the Figure.

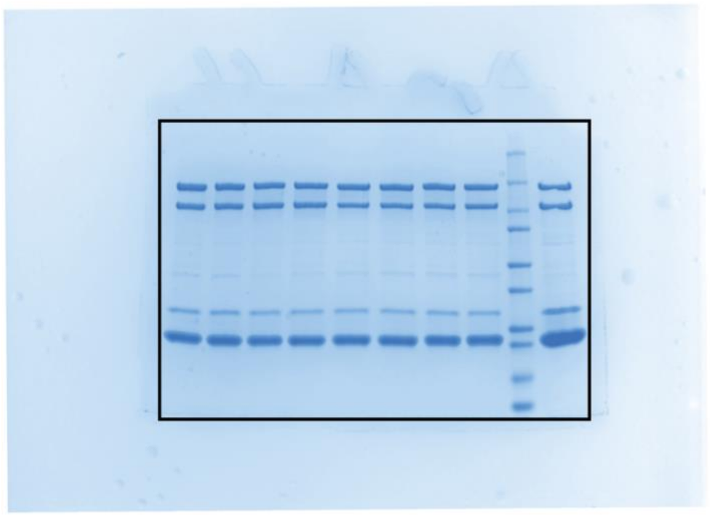

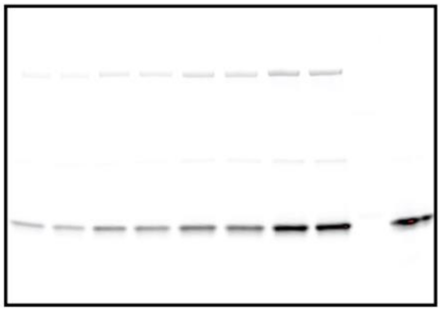

**Raw Data 6:** Uncropped, raw gel images that were used to create Fig. S6. A black box indicates where the gel image was cropped for its presentation in the Figure.

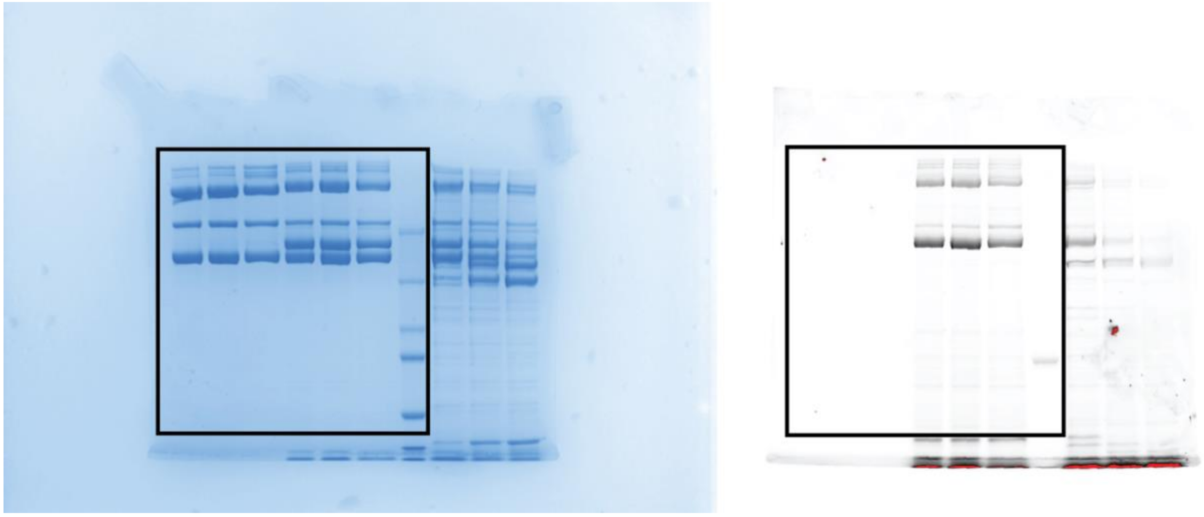

**Raw Data 7:** Uncropped, raw gel images that were used to create Fig. S7. A black box indicates where the gel image was cropped for its presentation in the Figure.

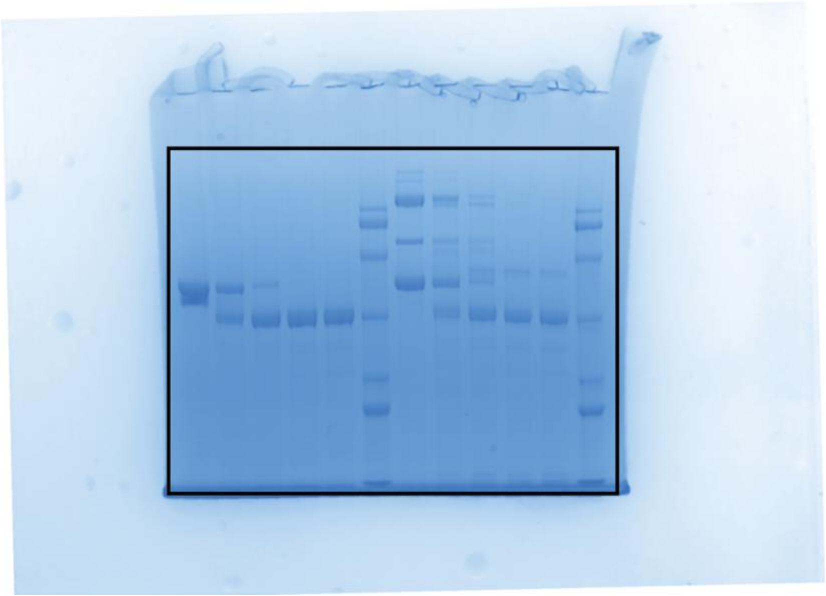

**Raw Data 8:** Uncropped, raw gel images that were used to create Fig. S9. A black box indicates where the gel image was cropped for its presentation in the Figure.

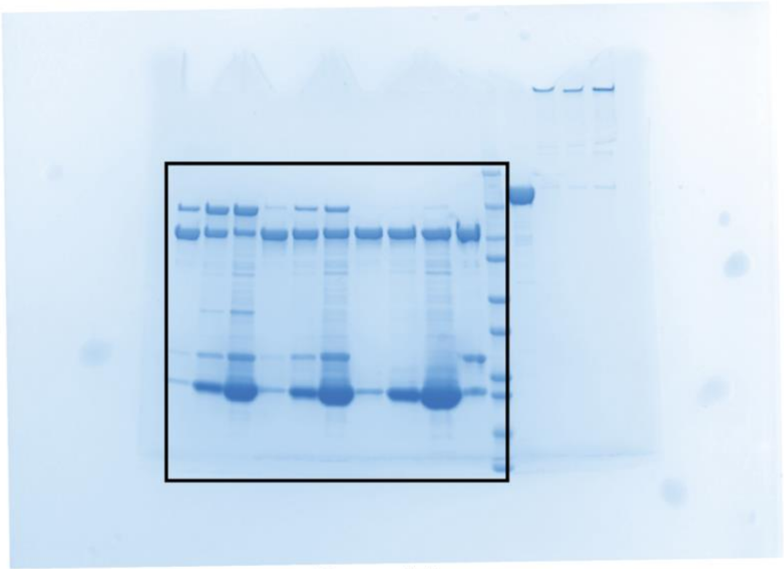

Panel A

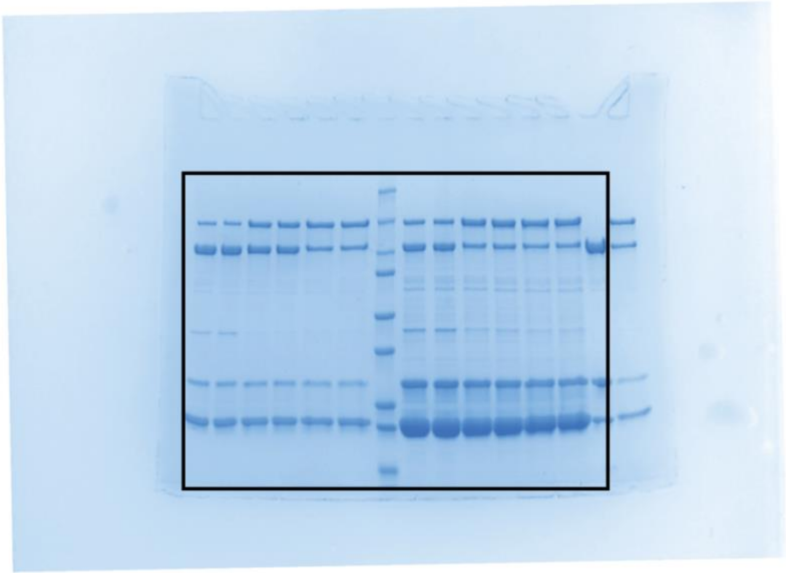

Panel B

**Raw Data 9:** Uncropped, raw gel images that were used to create Fig. S10. A black box indicates where the gel image was cropped for its presentation in the Figure.

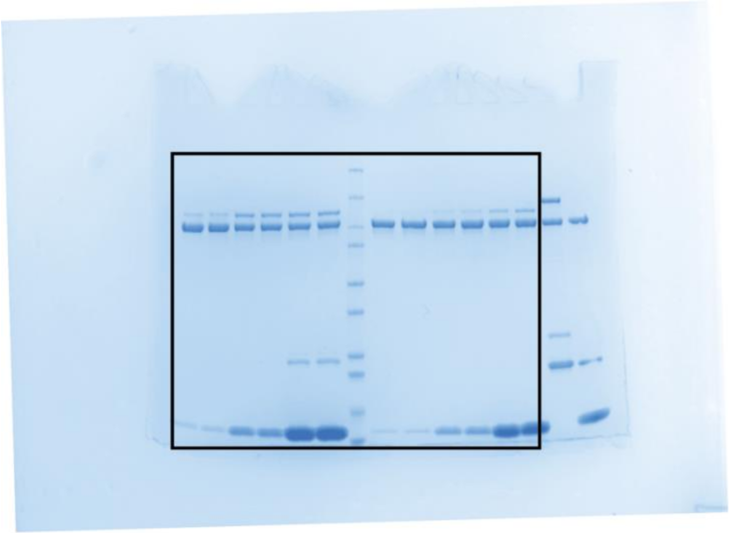
Panel A

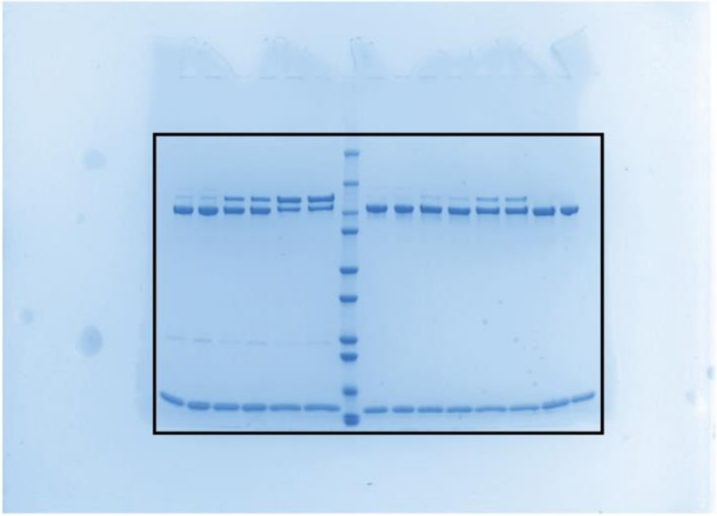
Panel B

**Raw Data 10:** Uncropped, raw gel images that were used to create Fig. S11. A black box indicates where the gel image was cropped for its presentation in the Figure.

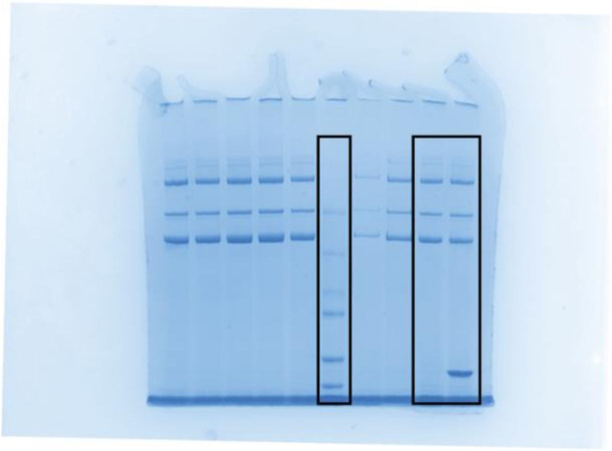

Panel A

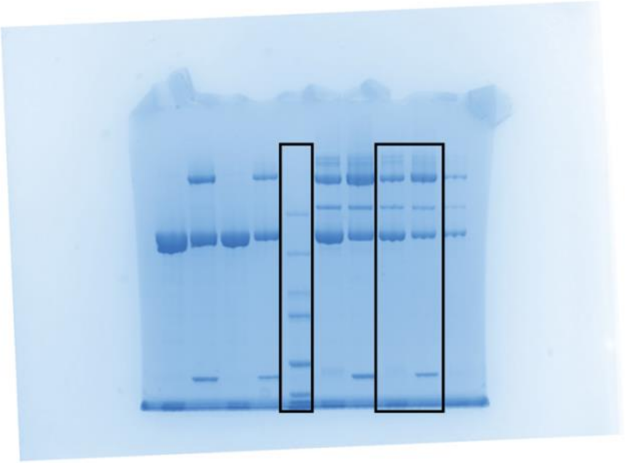

Panel B

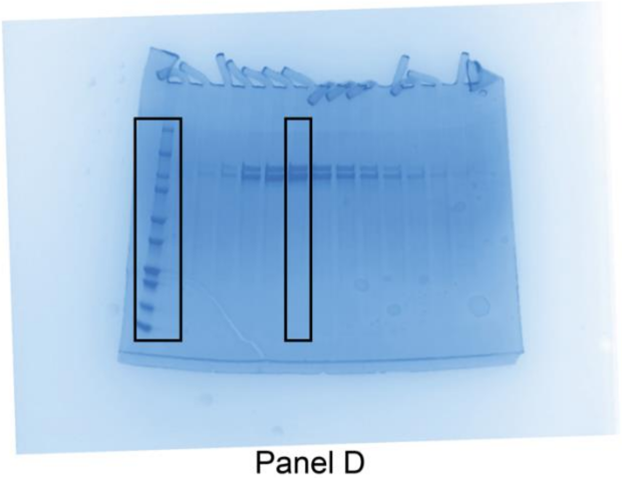

Panel D

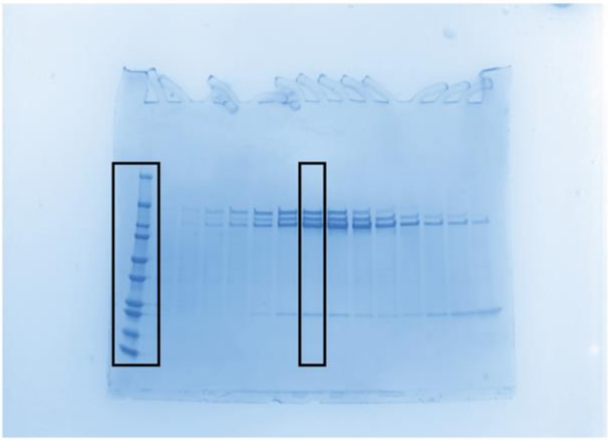

Panel E

**Raw Data 11:** Uncropped, raw gel images that were used to create Fig. S12. A black box indicates where the gel image was cropped for its presentation in the Figure.

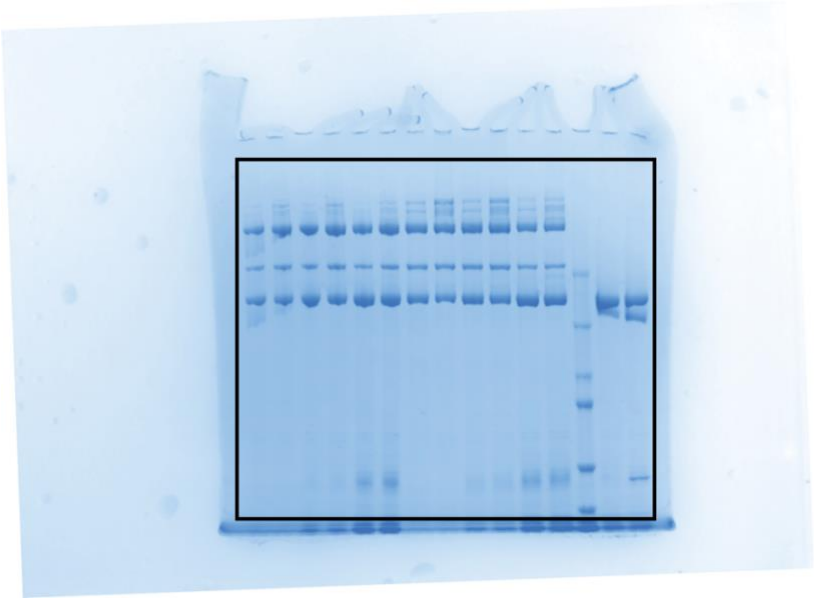

**Raw Data 12:** Uncropped, raw gel images that were used to create Fig. S16. A black box indicates where the gel image was cropped for its presentation in the Figure.

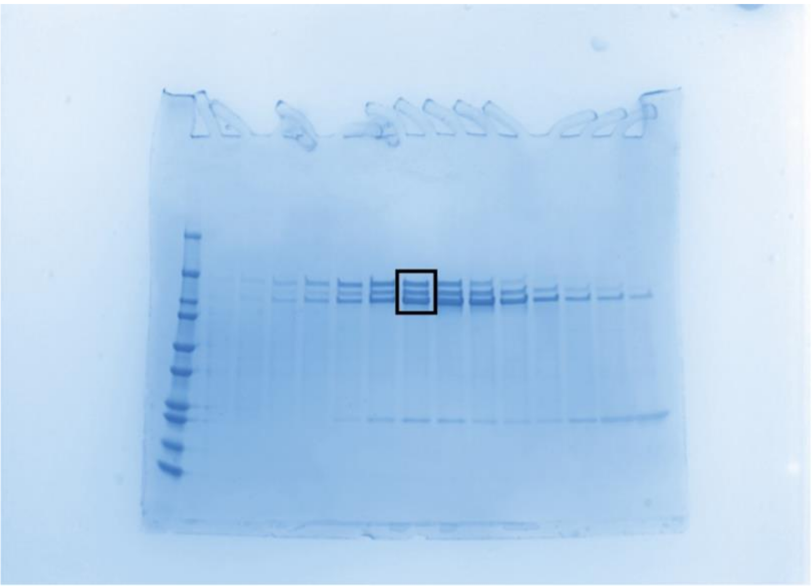

**Raw Data 13:** Uncropped, raw gel images that were used to create Fig. S27. A black box indicates where the gel image was cropped for its presentation in the Figure.
